# μMap-uAA: Photocatalytic proximity labeling targeted with single-residue precision

**DOI:** 10.64898/2026.08.05.742576

**Authors:** Min Sun Kang, Chun Li, Benito F. Buksh, David F. Fernández, David W. C. MacMillan

## Abstract

Mapping protein microenvironments with residue-level precision in living cells remains challenging. We report μMap-uAA, a genetically encoded proximity labeling platform that uses unnatural amino acid incorporation to install a tetrazine-quenched iridium photocatalyst at user-defined protein sites through click chemistry. Photocatalysis is activated by covalent attachment to the incorporated uAA, enabling localized catalytic labeling of proximal biomolecules. Applied to membrane receptors, μMap-uAA captures domain- and time-dependent GPCR interaction networks with residue-level spatial precision.

## Introduction

Cellular signaling is highly dependent on spatial organization, with protein function governed not only by molecular identity but also by local cellular context. Proteins often operate within heterogeneous microenvironments, where distinct structural domains can engage different interaction partners, regulatory factors, and biomolecules. However, methods capable of mapping these local protein environments with site-specific precision in living cells remain limited.

This challenge is especially pronounced in membrane proteins, whose functions are shaped by highly organized and dynamic membrane microenvironments. G protein-coupled receptors (GPCRs) provide a prominent example: extracellular regions mediate ligand recognition, while intracellular domains recruit signaling and trafficking machinery, and spatially distinct receptor surfaces coordinate cellular responses (**Figure 1A**). GPCRs also represent one of the largest and most intensively exploited drug target classes, with more than 500 approved drugs acting on 121 receptors and collectively accounting for approximately 36% of all marketed therapeutics.^1^ Indeed, several GPCR-targeted drugs are included on the World Health Organization Model List of Essential Medicines, and recent blockbuster therapies such as semaglutide and tirzepatide act on the glucagon-like peptide-1 receptor (GLP-1R) for the treatment of diabetes and obesity.^2^ Despite their central importance in pharmacology, the molecular composition and organization of GPCR microenvironments, particularly at nonterminal receptor regions, remain incompletely understood.

**Figure 1.**
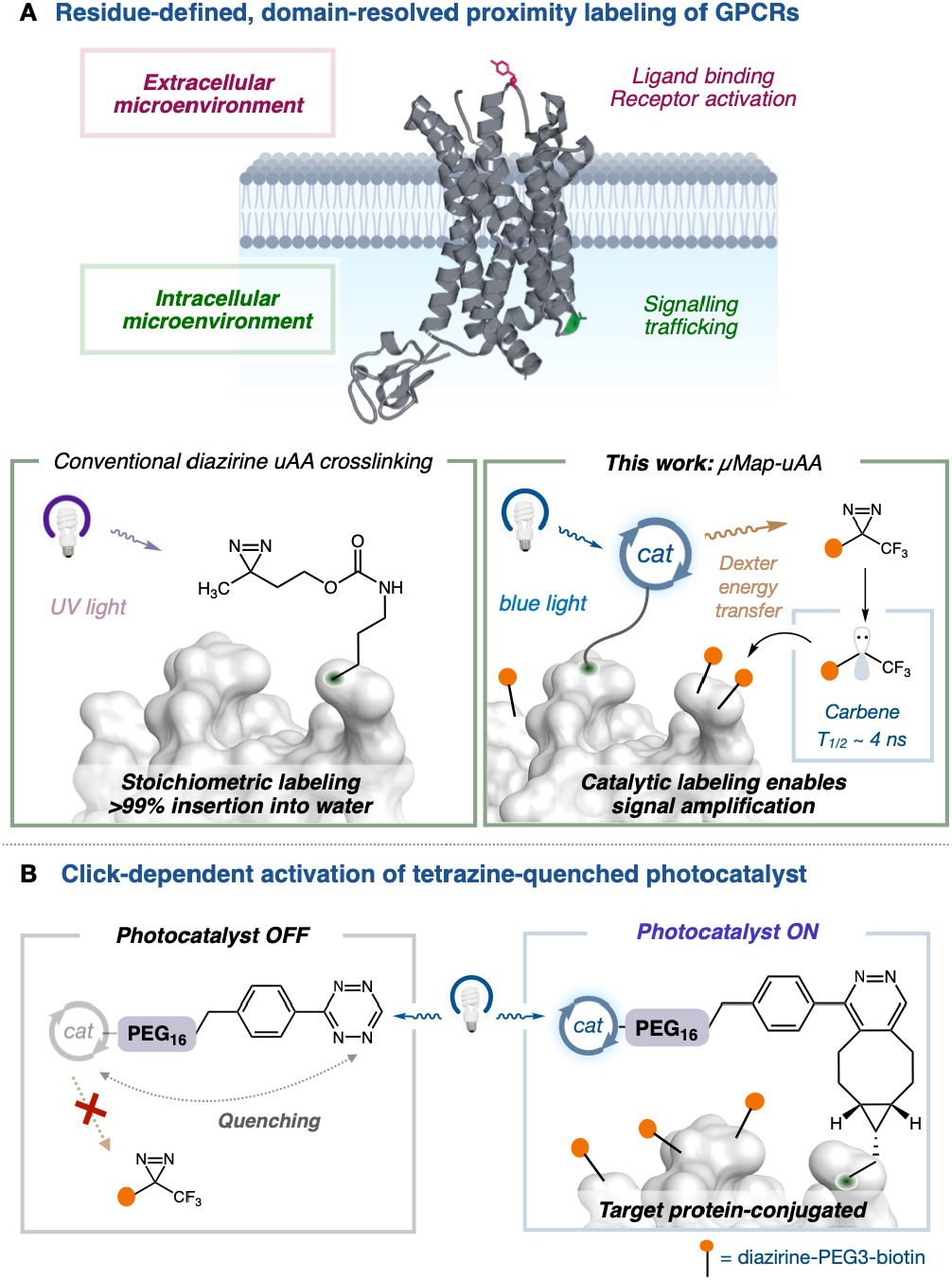
μMap–uAA enables residue-defined, domain-resolved mapping of GPCR microenvironments. A. Spatially distinct GPCR domains and comparison with conventional diazirine–uAA crosslinking. B. Click activation of a tetrazine-quenched photocatalyst on the target protein.

Our laboratory recently introduced μMap, a photocatalytic proximity labeling platform that enables spatially confined protein labeling through light-triggered generation of short-lived reactive intermediates.^3^ This approach has been successfully applied to a range of biological contexts, including profiling cell-surface protein–protein interactions, mapping immunological synapses,^4^ and interrogating dynamic intracellular microenvironments with precise temporal control over labeling.^5^ In parallel, conceptually related photocatalytic proximity labeling strategies have also been described, underscoring the broader utility of light-activated approaches for mapping spatially restricted interaction networks in living systems.^6,7^

Despite these advances, achieving residue-defined proximity labeling on complex membrane proteins remains challenging. Enzyme-fusion approaches can map receptor-proximal interaction networks, but labeling is often dictated by the fusion site and therefore biased toward receptor termini.^8^ Genetically encoded photo-crosslinking amino acids offer an orthogonal strategy for placing reactive groups at defined residues, including nonterminal domains such as extracellular loops.^9^ However, conventional diazirine-containing unnatural amino acids (uAAs) rely on direct UV activation and typically support only a single labeling event per incorporated amino acid, limiting signal amplification and sensitivity (**Figure 1A**).^10^

We therefore sought to combine the residue-level programmability of genetic code expansion with the catalytic amplification of μMap (**Figure 1B**). We hypothesized that an uAA-enabled microenvironment mapping strategy could enable amino acid-resolved, site-specific proximity labeling at defined positions across GPCR domains. In this platform, a bicyclononyne (BCN)-bearing uAA is genetically encoded at selected positions on the receptor, while a tetrazine-quenched iridium photocatalyst remains catalytically inactive until it is covalently attached through inverse electron demand Diels-Alder (IEDDA) reaction. This reaction-dependent installation restores localized photocatalytic activity at the receptor surface, enabling light-triggered generation of short-lived reactive intermediates for catalytic labeling of proximal biomolecules.

Tetrazines are widely exploited as bioorthogonal quenchers owing to their low-lying π ^*^ orbitals, which efficiently accept resonance energy transfer from proximal excited chromophores.^11^ In addition to energy-transfer pathways such as Förster Resonance Energy Transfer (FRET) or Through-Bond Energy Transfer (TBET),^12^ electron-deficient tetrazines can participate in photoinduced electron transfer (PeT) with sufficiently reducing excited states, providing a second nonradiative deactivation pathway.^13^ These mechanisms have been documented for tetrazine-linked fluorophores and, more directly, for cyclometalated iridium(III)–tetrazine complexes, whose strongly attenuated emission is restored upon IEDDA reaction with BCN.^14^ We therefore reasoned that appending a tetrazine to an iridium photocatalyst would suppress its phosphorescence and attenuate the excited-state pathways required for photocatalytic sensitization in solution. In this design, reaction with the BCN-bearing uAA is expected to eliminate the tetrazine-based quenching pathway and restore the long-lived excited state of protein-localized iridium catalyst species (**Figure 1B**).

## Results and Discussion

Consistent with this design, the tetrazine–iridium photocatalyst (**1**) displayed markedly attenuated emission relative to the BCN-reacted product (**2**) (**Figure 2A**). Upon addition of increasing equivalents of BCN-OH (**3**), we observed a progressive increase in emission intensity as the tetrazine was converted to the corresponding pyridazine adduct. This turn-on response supports a Click-depenent photocatalyst activation model in which tetrazine-mediated excited-state quenching suppresses photocatalytic activity prior to ligation, while IEDDA adduct formation relieves quenching and restores the emissive, catalytically competent iridium excited state. To determine whether tetrazine-mediated emission quenching translated into functional suppression of photocatalytic activity, we next evaluated diazirine sensitization using an *in vitro* bovine serum albumin (BSA) labeling assay adapted from the original μMap platform (**Figure 2B**).^3^ In this assay, blue-light excitation of an active iridium photocatalyst promotes energy transfer to a biotin-diazirine probe, generating a short-lived carbene that labels nearby protein residues. We reasoned that if the appended tetrazine attenuates the photoexcited state of iridium, **1** should nearby protein residues. We reasoned that if the appended tetrazine attenuates the photoexcited state of iridium, **1** should exhibit reduced diazirine sensitization relative to BSA was incubated with biotin-diazirine and irradiated with blue light in the presence of increasing concentrations of either **1** or **2**. The ligated catalyst **2** produced robust, concentration-dependent protein biotinylation, consistent with efficient iridium-mediated diazirine activation. In contrast, **1** generated substantially lower labeling across the same concentration range, indicating that the tetrazine-linked catalyst is functionally suppressed prior to ligation. Importantly, mixtures containing **1** and **2** at matched total iridium concentration did not increase labeling beyond that observed with **2** alone, arguing against nonspecific optical or concentration-dependent artifacts (**Figure S2**). Together, these results support a reaction-dependent mechanism for μMap-uAA.

**Figure 2.**
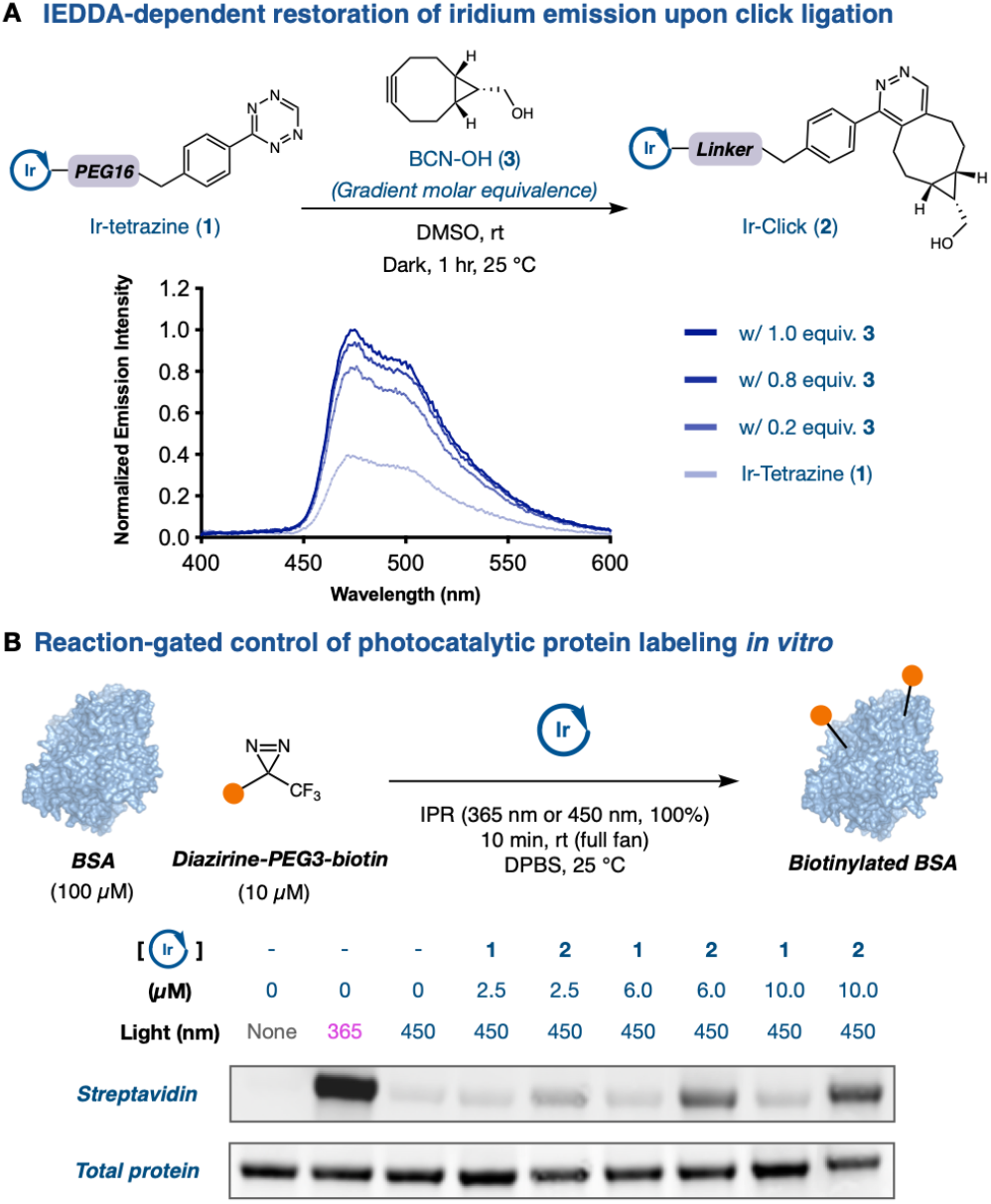
(A) Emission spectra of **1** after incubation with 0, 0.2, 0.8 and an equimolar equivalent of **3.** IEDDA restores iridium emission, indicating relief of tetrazine-mediated quenching. (B) *In vitro* BSA labeling assay comparing the photocatalytic activity of **1** and **2**. BSA was incubated with diazirine–biotin in the absence or presence of the indicated iridium photocatalyst and irradiated with UV or blue light, as shown. Biotinylation was detected by streptavidin blot, with total protein shown as a loading control. Band intensities were quantified by densitometry (n=3 biological replicates); see **Figure S2.C** for quantification. BSA structure shown from PDB 4F5S.

With a reaction-dependent photocatalytic system in hand, we sought to determine whether genetically encoded BCN could support live-cell catalyst installation across distinct protein contexts. We first examined β-actin and the nuclear protein NONO (Non-POU domain-containing octamer-binding protein) to assess whether **1** could engage genetically encoded BCN handles in living cells. Pulse-chase analysis demonstrated that **1** crossed the plasma membrane and covalently engaged the genetically encoded BCN handle on intracellular β-actin in live cells (**Figure S1.A**). Similarly, BCN-containing NONO variant showed robust iridium-dependent self-labeling by immunoblot analysis (**Figure S1.B**), consistent with successful catalyst installation and proximity labeling in a nuclear protein context.

Having established catalyst engagement on non-receptor proteins in live cells, we next evaluated the μMap-uAA platform for residue-defined interactome mapping using C-C chemokine receptor type 5 (CCR5) as a model GPCR. CCR5 is a therapeutically relevant class A GPCR with established roles in immune signaling and as a co-receptor for human immunodeficiency virus (HIV) entry.^15^ In addition, genetic code expansion has previously been applied to CCR5,^16,17^ supporting its use as a model system for site-specific catalyst installation (**Figure 3A**). To enable defined photocatalyst localization, we first established amber suppression in CCR5 by incorporating a BCN-lysine (**4**) at two positions: F96^*^, positioned in extracellular loop 1 (ECL1), and L352^*^, located in the C-terminal tail on the cytoplasmic face of the receptor. Both sites have been reported to tolerate mutation without substantial perturbation of native CCR5 function, and confocal microscopy confirmed that BCN incorporation at either position did not disrupt plasma membrane localization (**Figure S3.F**).

**Figure 3.**
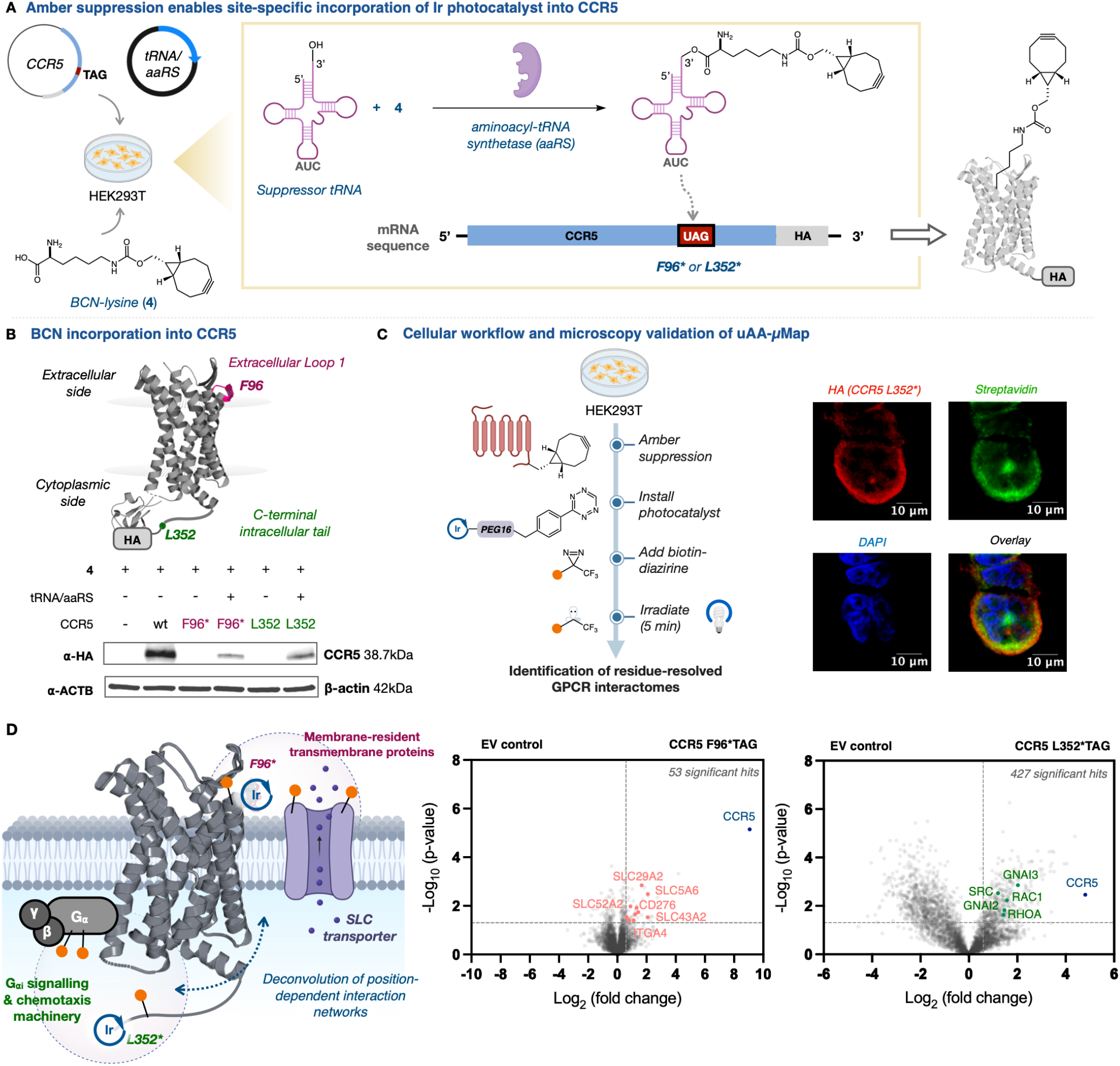
(A) Schematic overview of amber suppression for site-specific incorporation of BCN into CCR5. HEK293T cells were transfected with CCR5 bearing an amber codon at either F96 or L352 together with an orthogonal suppressor tRNA/aaRS pair. (B) Selection and validation of CCR5 amber suppression sites. F96 is positioned in extracellular loop 1, whereas L352 is located in the C-terminal intracellular tail, enabling photocatalyst placement on spatially distinct receptor domains. CCR5 structure shown from PDB 4MBS. Because 4MBS does not resolve the full C-terminal tail, L352 was depicted manually. HA western blot confirmed robust expression of wild-type (wt) CCR5 and production of full-length F96^*^ and L352^*^ CCR5 only when **4** and the tRNA/aaRS pair were present. (C) Live-cell μMap-uAA workflow and microscopy validation. Robust streptavidin signal in HA-positive cells supports successful proximity labeling from a site-specifically installed photocatalyst in living cells. Scale bars, 10 μm. (D) Quantitative proteomics of CCR5-proximal proteins labeled from F96^*^ or L352^*^ . Volcano plots compare each CCR5 amber mutant against the empty-vector control. F96^*^ labeling enriched membrane-resident and transmembrane proteins, including SLC transporters and cell-surface proteins, whereas L352^*^ labeling enriched intracellular CCR5-associated proteins, including G protein subunits and signaling-associated factors. Three biological replicates were performed for each condition. Significance was determined using a two-tailed Student’s t-test. Significant hits are shown above the indicated threshold: log_2_(fold change) > 1 and -log(p-value) > 1.3.

Successful unnatural amino acid incorporation was first assessed by immunoblotting against the C-terminal HA tag. For both F96^*^ and L352^*^ CCR5 mutants, full-length receptor was detected only in the presence of **4** and the engineered aminoacyl-tRNA synthetase (aaRS)/suppressor tRNA pair, whereas omission of the amber suppression machinery abolished the full-length HA-positive band, consistent with translation termination at the amber codon (**Figure 3B**). BCN incorporation was further validated using a BCN-selective tetrazine–TAMRA probe. Treatment of cell lysates with increasing concentrations of tetrazine–TAMRA produced a dose-dependent fluorescent band at the expected molecular weight of CCR5 (**Figure S3.B**,**D**). Subsequent HA immunoblotting confirmed overlap between the HA-positive and TAMRA-positive signals, supporting selective labeling of full-length CCR5 (**Figure S3.C**,**E**).

Having validated site-specific incorporation of **4** into CCR5, we next evaluated whether this handle could support live-cell installation of the iridium photocatalyst and proximity labeling from defined receptor domains. HEK293T cells expressing BCN-containing CCR5 were incubated with **1** to covalently install the photocatalyst by IEDDA, followed by addition of biotin–diazirine and blue-light irradiation for 5 min (**Figure 3C**). Confocal analysis was performed on fixed cells using DAPI to mark nuclei, anti-HA staining to visualize CCR5, and streptavidin staining to detect biotinylated proteins. Under our μMap-uAA conditions, robust streptavidin signal was observed in CCR5-expressing cells, confirming successful live-cell proximity labeling (**Figure 3C**). For CCR5 L352^*^, streptavidin signal was observed throughout intracellular receptor-positive regions, indicating catalyst placement at the cytoplasmic C-terminal tail and labeling from an intracellular receptor domain. In contrast, labeling from CCR5 F96^*^ showed a more peripheral, membrane-localized biotinylation pattern, consistent with photocatalyst installation at the extracellular loop region (**Figure S3.G**). As an independent validation of the live-cell labeling workflow, streptavidin enrichment followed by anti-HA immunoblotting confirmed recovery of full-length CCR5 for both L352^*^ and F96^*^ constructs (**Figure S3.H, I**).

Although the presence or absence of the tRNA/aaRS pair provided a convenient control for initial validation, this comparison is less suitable for quantitative proteomics. The tRNA/aaRS pair can incorporate **4** into endogenous proteins at native amber codons regardless of target protein expression, generating target-independent labeling background. We therefore used a matched control design in which both samples contained **4** and the tRNA/aaRS pair, but only the directed sample expressed CCR5; the control received empty vector. This design controls for suppression -dependent background, allowing enrichment to be attributed specifically to CCR5-proximal labeling.

Using this refined control design, μMap-uAA and subsequent data-independent acquisition (DIA)-based label-free quantitative proteomics identified CCR5 as the top enriched protein in both the extracellular and intracellular labeling datasets (**Figure 3D**). In the extracellular F96^*^ dataset, enriched proteins were biased toward plasma membrane-associated factors, including multipass solute carriers such as SLC5A6, SLC43A2, and SLC22A5, the ABC transporter ABCC1, and cell-surface proteins including CD276 and IL17RA. Notably, the dataset also recovered ITGA4, an adhesion receptor co-expressed with CCR5 in migratory immune-cell populations and implicated in leukocyte trafficking.^19^ By contrast, the intracellular L352^*^ dataset was selectively enriched for heterotrimeric G protein subunits, including GNAI2 and GNAI3, consistent with the established coupling of CCR5 to inhibitory G_α_ proteins at its cytoplasmic tail.^20^ The larger number of enriched proteins observed from L352^*^ labeling likely reflects the greater protein density and richer signaling and trafficking network at the cytoplasmic face of CCR5. The enrichment of canonical intracellular CCR5 effectors in the L352^*^ dataset, together with their absence from the F96^*^ pulldown, supports the site-resolved spatial precision of μMap-uAA.

We then investigated whether μMap-uAA could resolve inhibitor-induced remodeling of the CCR5-proximal proteome. Maraviroc, the active ingredient in Selzentry™, is a clinically used CCR5 antagonist that allosterically blocks CCR5-dependent HIV entry by stabilizing a conformation incompatible with viral engagement.^21^ Maraviroc has also been reported to inhibit ligand-induced CCR5 internalization,^22^ yet the mechanism of antagonist binding remodels receptor-proximal endocytic machinery at the cell surface remains less well defined. Given that μMap-uAA enables residue-defined photocatalyst placement outside the maraviroc-binding pocket (**Figure 4A**), we reasoned that this platform could capture antagonist-dependent changes in CCR5-proximal interaction networks in living cells (**Figure 4B**). In the F96^*^ labeling dataset, quantitative proteomics comparing untreated and maraviroc-treated cells (100 nM, 1 hr) revealed marked depletion of AP-2 adaptor complex subunits from the CCR5-proximal proteome, particularly AP2S1 and AP2M1 (**Figure 4C**). This decrease is consistent with maraviroc-induced uncoupling of CCR5 from clathrin-mediated endocytic machinery. In parallel, L352^*^ labeling identified maraviroc-sensitive intracellular CCR5-proximal proteins, including CADM1, ITGA4, and SERINC5, suggesting antagonist-dependent remodeling of cell-adhesion and membrane-associated interaction networks from the cytoplasmic-facing receptor environment (**Figure S4.A**,**B**). Confocal imaging further showed reduced overlap between wild-type CCR5 and AP2S1 following maraviroc treatment (500 nM, 1 hr), providing orthogonal validation that μMap-uAA captures antagonist-induced remodeling of the native CCR5 endocytic environment (**Figure 4C**).

**Figure 4.**
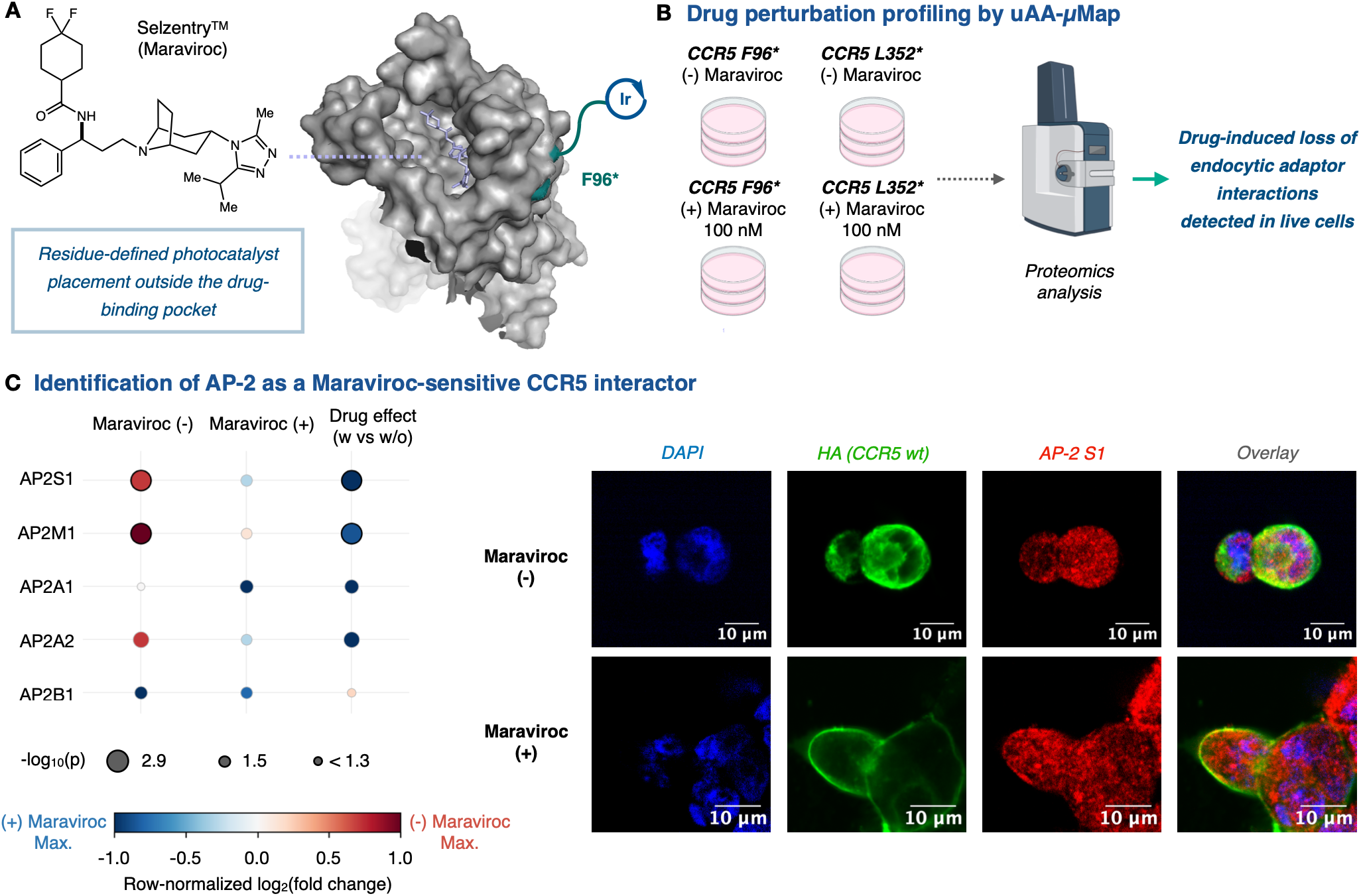
(A) Maraviroc binds within an allosteric pocket of CCR5, whereas the μMap-uAA photocatalyst is installed outside the drug-binding site. CCR5–maraviroc structure shown from PDB 4MBS. (B) Experimental workflow for drug perturbation profiling. Cells expressing CCR5 F96^*^ or L352^*^ were treated with or without maraviroc and subjected to μMap-uAA followed by quantitative proteomics. (C) Identification of AP-2 adaptor complex subunits as maraviroc-sensitive CCR5-proximal proteins. Bubble plot shows AP-2 subunit enrichment in the absence or presence of Maraviroc (100 nM, 1 hr) and the drug effect calculated by comparing maraviroc-treated versus untreated conditions. Bubble size indicates statistical significance, and color indicates log_2_(fold change). Confocal imaging using wild-type (wt) CCR5, rather than the amber mutant, showed reduced overlap between CCR5 and AP2S1 following maraviroc treatment (500 nM, 1 hr), providing orthogonal validation. Scale bars, 10 μm.

We next asked whether μMap–uAA could capture agonist-driven remodeling of CCR5-proximal biology. CCL5/RANTES stimulation induced time-dependent changes in the CCR5-proximal proteome, with gene ontology analysis revealing distinct enrichment patterns across basal, early, and prolonged ligand exposure (**Figure S4.C**). Prolonged stimulation increased enrichment of ARRB2, consistent with ligand-induced CCR5 internalization, and also revealed progressive enrichment of GNAO1, a candidate CCR5-proximal signaling component. Confocal microscopy further supported increased CCR5–GNAO1 proximity after 2 h of CCL5 treatment (**Figure S4.D**). These findings highlight the ability of μMap–uAA to detect ligand-induced GPCR interactome remodeling with temporal precision.

To evaluate the generality of the platform, we next applied μMap-uAA to GLP-1R. Recent ligand-directed proximity labeling of GLP-1R using a GLP-1-peroxidase conjugate provides a useful framework for profiling receptor-associated membrane proteins during agonist stimulation.^23^ As an orthogonal strategy, we asked whether residue-defined photocatalyst placement by μMap-uAA could access GLP-1R microenvironments independently of ligand-directed catalyst recruitment. We selected F61 as the catalyst-installation site, because amber-mutagenesis mapping has revealed that F61 is in the GLP-1R extracellular domain, does not directly contact the GPL-1R agonist exendin-4, and tolerates uAA incorporation without compromising exendin-4–stimulated signaling.^24^ Thus, F61 enabled extracellular photocatalyst placement while minimizing perturbation of the peptide-binding interface (**Figure 5A**). The μMap-uAA and subsequent quantitative DIA proteomics identified GLP-1R as the most strongly enriched protein, validating receptor-directed labeling from the F61^*^ site (**Figure 5B**). The basal interactome recovered known GLP-1R-associated and membrane-proximal proteins, including heterotrimeric G proteins,^25^ endocytic machinery,^26^ RAB5-family early endosomal trafficking regulators,^27^ CD81,^28^ TFRC,^29^ and zinc-importing ZIP transporters SLC39A6 and SLC39A10.^30^

**Figure 5.**
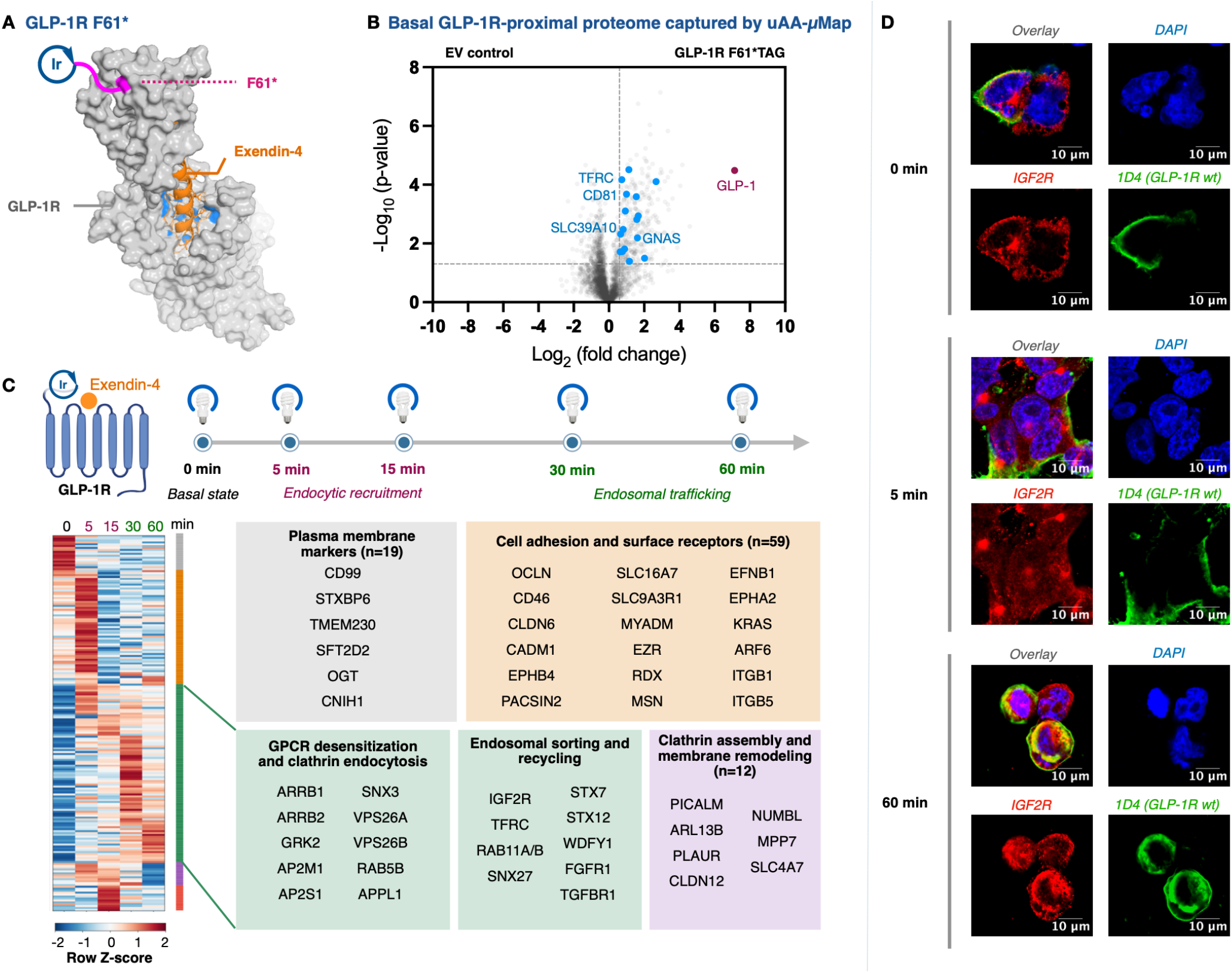
μMap-uAA captures basal and agonist-regulated GLP-1R microenvironments. (A) Structural positioning of GLP-1R F61^*^ relative to the exendin-4 binding interface. Exendin-4 is shown in orange within the receptor, with proximal binding-site residues highlighted in blue. The F61 site, shown in magenta, is located distal to the ligand-binding pocket. Structure from PDB 7LLL. (B) Quantitative proteomics identifies basal GLP-1R-proximal proteins captured by μMap-uAA from F61^*^ . Volcano plot compares GLP-1R F61^*^ labeling against the empty-vector control. Blue data points represent known GLP-1R interactors from the BioGRID database.^18^ (C) Time-resolved μMap-uAA captures 100 nM exendin-4-induced remodeling of the GLP-1R-proximal proteome, including recruitment of endocytic, endosomal, sorting/recycling, and membrane-remodeling proteins. Heat map shows row Z-scored enrichment profiles across the 0–60 min stimulation time course. (D) Confocal microscopy validates time-dependent GLP-1R and IGF2R redistribution following exendin-4 (500 nM) stimulation. GLP-1R was visualized using the C-terminus 1D4 epitope tag, and IGF2R was used as a late endosomal marker. Scale bars, 10 μm.

We next examined agonist-induced remodeling of the GLP-1R-proximal proteome. Cells expressing GLP-1R F61^*^ were stimulated with 100 nM exendin-4 and subjected to time-resolved μMap-uAA over 0–60 min (**Figure 5C**). Hierarchical clustering revealed temporally ordered remodeling, with early enrichment of plasma membrane, receptor desensitization, and clathrin-mediated endocytic factors, including GRK2, ARRB1/2, and AP-2 complex subunits. Later time points showed increased enrichment of endosomal sorting, recycling, and membrane-remodeling proteins, including RAB family members, SNX proteins, VPS26A/B, IGF2R, and TFRC. Consistent with this late endosomal trafficking signature, confocal microscopy showed increased intracellular overlap between internalized GLP-1R and IGF2R after prolonged exendin-4 stimulation, particularly at 60 min (**Figure 5D**).

## Conclusion

In conclusion, we disclose a chemogenetic, residue-defined proximity labeling platform for mapping protein microenvironments in live cells. By combining genetic code expansion with a bioorthogonally activated iridium photocatalyst, μMap-uAA enables photocatalyst placement at user-defined amino acid positions on a protein of interest. Because catalyst installation is programmed at the DNA level through amber codon mutagenesis, this strategy is readily adaptable to new protein sites and systems. We show that tetrazine-mediated photocatalyst quenching suppresses background labeling prior to ligation, while reaction with a BCN-bearing unnatural amino acid restores photocatalytic activity and enables localized biotin–diazirine activation. Although the chemistry is generalizable, membrane receptors provide a particularly powerful setting for μMap-uAA because distinct extracellular loops, intracellular tails, and other nonterminal regions remain difficult to interrogate using conventional proximity labeling approaches. Applied to CCR5 and GLP-1R, μMap-uAA enables interrogation of extracellular and intracellular receptor environments from defined residue positions, revealing position-dependent interactomes as well as ligand- and drug-induced remodeling of receptor-proximal proteomes. This work establishes μMap-uAA as a residue-defined and operationally accessible strategy for probing dynamic protein microenvironments in live cells, with utility for studying GPCR signaling, trafficking, pharmacology, and ligand discovery.

## AUTHOR INFORMATION

### Authors

**Min Sun Kang** *- Merck Center for Catalysis at Princeton University, Princeton, New Jersey, 08544, United States; Department of Chemistry, Princeton University, Princeton, New Jersey 08544, United States*

**Chun Li** *- Merck Center for Catalysis at Princeton University, Princeton, New Jersey, 08544, United States; Department of Chemistry, Princeton University, Princeton, New Jersey 08544, United States*

**Benito F. Buksh** *- Merck Center for Catalysis at Princeton University, Princeton, New Jersey, 08544, United States; Department of Chemistry, Princeton University, Princeton, New Jersey 08544, United States*

**David F. Fernández** *- Merck Center for Catalysis at Princeton University, Princeton, New Jersey, 08544, United States; Department of Chemistry, Princeton University, Princeton, New Jersey 08544, United States*

### Author Contributions

The manuscript was written through contributions of all authors. / All authors have given approval to the final version of the manuscript.

### Notes

D.W.C.M. declares an ownership interest in Dexterity Pharma, which has commercialized materials used in this work.

## ACKNOWLEDGMENT

Research reported in this work was supported by the Princeton Branch of the Ludwig Institute for Cancer Research. We would like to thank Gary S. Laevsky and Sha Wang of the Confocal Imaging Facility, a Nikon Center of Excellence, in the Department of Molecular Biology at Princeton University for instrument use and technical advice. Illustrations were created with BioRender.com.

---

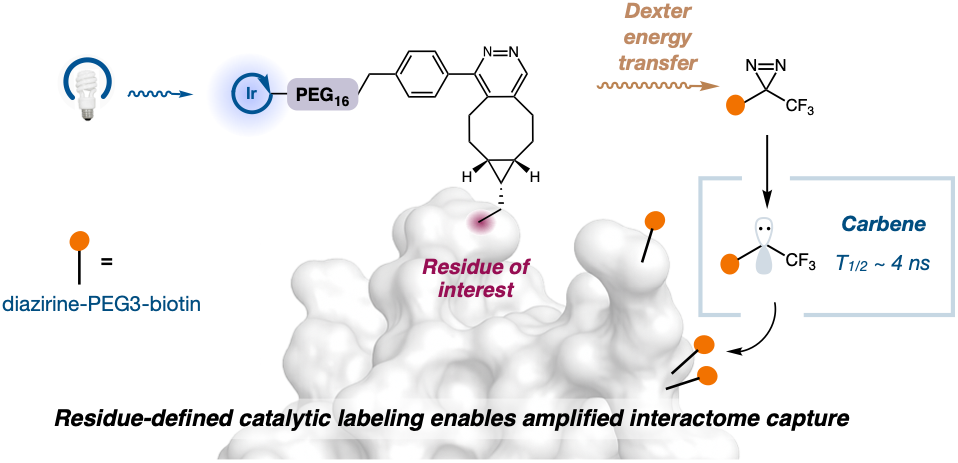

## REFERENCES

(1) Lorente, J. S.; Sokolov, A. V.; Ferguson, G.; Schiöth, H. B.; Hauser, A. S.; Gloriam, D. E. GPCR Drug Discovery: New Agents, Targets and Indications. Nat Rev Drug Discov 2025, 24 (6), 458–479. 10.1038/s41573-025-01139-y.

(2) Drucker, D. J. GLP-1-Based Therapies for Diabetes, Obesity and Beyond. Nat Rev Drug Discov 2025, 24 (8), 631–650. 10.1038/s41573-025-01183-8.

(3) Geri, J. B.; Oakley, J. V.; Reyes-Robles, T.; Wang, T.; McCarver, S. J.; White, C. H.; Rodriguez-Rivera, F. P.; Parker, D. L.; Hett, E. C.; Fadeyi, O. O.; Oslund, R. C.; MacMillan, D. W. C. Microenvironment Mapping via Dexter Energy Transfer on Immune Cells. Science 2020, 367 (6482), 1091–1097. 10.1126/science.aay4106.

(4) Huth, S. W.; Geri, J. B.; Oakley, J. V.; MacMillan, D. W. C. μMap-Interface: Temporal Photoproximity Labeling Identifies F11R as a Functional Member of the Transient Phagocytic Surfaceome. J. Am. Chem. Soc. 2024, 146 (47), 32255–32262. 10.1021/jacs.4c11058.

(5) Pan, C.; Knutson, S. D.; Huth, S. W.; MacMillan, D. W. C. μMap Proximity Labeling in Living Cells Reveals Stress Granule Disassembly Mechanisms. Nat Chem Biol 2025, 21 (4), 490–500. 10.1038/s41589-024-01721-2.

(6) Huang, Z.; Liu, Z.; Xie, X.; Zeng, R.; Chen, Z.; Kong, L.; Fan, X.; Chen, P. R. Bioorthogonal Photocatalytic Decaging-Enabled Mitochondrial Proteomics. J. Am. Chem. Soc. 2021, 143 (44), 18714–18720. 10.1021/jacs.1c09171.

(7) Zhai, Y.; Huang, X.; Zhang, K.; Huang, Y.; Jiang, Y.; Cui, J.; Zhang, Z.; Chiu, C. K. C.; Zhong, W.; Li, G. Spatiotemporal-Resolved Protein Networks Profiling with Photoactivation Dependent Proximity Labeling. Nat Commun 2022, 13 (1), 4906. 10.1038/s41467-022-32689-z.

(8) Samavarchi-Tehrani, P.; Samson, R.; Gingras, A.-C. Proximity Dependent Biotinylation: Key Enzymes and Adaptation to Proteomics Approaches* . Molecular & Cellular Proteomics 2020, 19 (5), 757–773. 10.1074/mcp.R120.001941.

(9) Shah, U. H.; Toneatti, R.; Gaitonde, S. A.; Shin, J. M.; González-Maeso, J. Site-Specific Incorporation of Genetically Encoded Photo-Crosslinkers Locates the Heteromeric Interface of a GPCR Complex in Living Cells. Cell Chemical Biology 2020, 27 (10), 1308–1317.e4. 10.1016/j.chembiol.2020.07.006.

(10) Dubinsky, L.; Krom, B. P.; Meijler, M. M. Diazirine Based Photoaffinity Labeling. Bioorganic & Medicinal Chemistry 2012, 20 (2), 10.1016/j.bmc.2011.06.066. 554–570.

(11) Carlson, J. C. T.; Meimetis, L. G.; Hilderbrand, S. A.; Weissleder, R. BODIPY–Tetrazine Derivatives as Superbright Bioorthogonal Turn-on Probes. Angewandte Chemie 2013, 125 (27), 7055–7058. 10.1002/ange.201301100.

(12) Loredo, A.; Tang, J.; Wang, L.; Wu, K.-L.; Peng, Z.; Xiao, H. Tetrazine as a General Phototrigger to Turn on Fluorophores. Chem. Sci. 2020, 11 10.1039/D0SC01009J.(17), 4410–4415.

(13) Shen, T.; Li, X.; Liu, X. Photoinduced Electron Transfer Endows Fluorogenicity in Tetrazine-Based near-Infrared Labels. Mater. Chem. Front. 2024, 8 10.1039/D3QM01217D.(9), 2135–2141.

(14) Li, S. P.-Y.; Yip, A. M.-H.; Liu, H.-W.; Lo, K. K.-W. Installing an Additional Emission Quenching Pathway in the Design of Iridium(III)-Based Phosphorogenic Biomaterials for Bioorthogonal Labelling and Imaging. Biomaterials 2016, 103, 305–313. 10.1016/j.biomaterials.2016.06.065.

(15) Alkhatib, G.; Combadiere, C.; Broder, C. C.; Feng, Y.; Kennedy, P. E.; Murphy, P. M.; Berger, E. A. CC CKR5: A RANTES, MIP-1α, MIP-1β Receptor as a Fusion Cofactor for Macrophage-Tropic HIV-1. Science 1996, 272 (5270), 1955–1958. 10.1126/science.272.5270.1955.

(16) Ye, S.; Köhrer, C.; Huber, T.; Kazmi, M.; Sachdev, P.; Yan, E. C. Y.; Bhagat, A.; RajBhandary, U. L.; Sakmar, T. P. Site-Specific Incorporation of Keto Amino Acids into Functional G Protein-Coupled Receptors Using Unnatural Amino Acid Mutagenesis. Journal of Biological Chemistry 2008, 283 (3), 1525–1533. 10.1074/jbc.M707355200.

(17) Mattheisen, J. M.; Wollowitz, J. S.; Huber, T.; Sakmar, T. P. Genetic Code Expansion to Enable Site-specific Bioorthogonal Labeling of Functional G Protein-coupled Receptors in Live Cells. Protein Science 2023, 32 (2), e4550. 10.1002/pro.4550.

(18) Stark, C.; Breitkreutz, B.-J.; Reguly, T.; Boucher, L.; Breitkreutz, A.; Tyers, M. BioGRID: A General Repository for Interaction Datasets. Nucleic Acids Research 2006, 34 (suppl_1), D535–D539. 10.1093/nar/gkj109.

(19) Deffner, M.; Schneider-Hohendorf, T.; Schulte-Mecklenbeck, A.; Falk, S.; Lu, I.-N.; Ostkamp, P.; Müller-Miny, L.; Schumann, E. M.; Goelz, S.; Cahir-McFarland, E.; Thakur, K. T.; De Jager, P. L.; Klotz, L.; Meyer Zu Hörste, G.; Gross, C. C.; Wiendl, H.; Grauer, O. M.; Schwab, N. Chemokine-Mediated Cell Migration into the Central Nervous System in Progressive Multifocal Leukoencephalopathy. Cell Reports Medicine 2024, 5 (7), 101622. 10.1016/j.xcrm.2024.101622.

(20) Zhao, J.; Ma, L.; Wu, Y.-L.; Wang, P.; Hu, W.; Pei, G. Chemokine Receptor CCR5 Functionally Couples to Inhibitory G Proteins and Undergoes Desensitization. Journal of Cellular Biochemistry 1998, 71 (1), 36–45. 10.1002/(SICI)1097-4644(19981001)71:1%3C36::AID-JCB4%3E3.0.CO;2-2.

(21) Dorr, P.; Westby, M.; Dobbs, S.; Griffin, P.; Irvine, B.; Macartney, M.; Mori, J.; Rickett, G.; Smith-Burchnell, C.; Napier, C.; Webster, R.; Armour, D.; Price, D.; Stammen, B.; Wood, A.; Perros, M. Maraviroc (UK-427,857), a Potent, Orally Bioavailable, and Selective Small-Molecule Inhibitor of Chemokine Receptor CCR5 with Broad-Spectrum Anti-Human Immunodeficiency Virus Type 1 Activity. Antimicrob Agents Chemother 2005, 49 (11), 4721–4732. 10.1128/AAC.49.11.4721-4732.2005.

(22) Nakata, H.; Kruhlak, M.; Kamata, W.; Ogata-Aoki, H.; Li, J.; Maeda, K.; Ghosh, A. K.; Mitsuya, H. Effects of CC Chemokine Receptor 5 (CCR5) Inhibitors on the Dynamics of CCR5 and CC-Chemokine–CCR5 Interactions. Antiviral Therapy 2010, 15 (3), 321–331. 10.3851/IMP1529.

(23) Dang, T.; Yu, J.; Cao, Z.; Zhang, B.; Li, S.; Xin, Y.; Yang, L.; Lou, R.; Zhuang, M.; Shui, W. Endogenous Cell Membrane Interactome Mapping for the GLP-1 Receptor in Different Cell Types. Nat Chem Biol 2025, 21 (2), 256–267. 10.1038/s41589-024-01714-1.

(24) Koole, C.; Reynolds, C. A.; Mobarec, J. C.; Hick, C.; Sexton, P. M.; Sakmar, T. P. Genetically Encoded Photocross-Linkers Determine the Biological Binding Site of Exendin-4 Peptide in the N-Terminal Domain of the Intact Human Glucagon-like Peptide-1 Receptor (GLP-1R). Journal of Biological Chemistry 2017, 292 (17), 7131–7144. 10.1074/jbc.M117.779496.

(25) Graaf, C. de; Donnelly, D.; Wootten, D.; Lau, J.; Sexton, P. M.; Miller, L. J.; Ahn, J.-M.; Liao, J.; Fletcher, M. M.; Yang, D.; Brown, A. J. H.; Zhou, C.; Deng, J.; Wang, M.-W. Glucagon-Like Peptide-1 and Its Class B G Protein–Coupled Receptors: A Long March to Therapeutic Successes. Pharmacological Reviews 2016, 68 (4), 954–1013. 10.1124/pr.115.011395.

(26) Buenaventura, T.; Kanda, N.; Douzenis, P. C.; Jones, B.; Bloom, S. R.; Chabosseau, P.; Corrêa, I. R., Jr.; Bosco, D.; Piemonti, L.; Marchetti, P.; Johnson, P. R.; Shapiro, A. M. J.; Rutter, G. A.; Tomas, A. A Targeted RNAi Screen Identifies Endocytic Trafficking Factors That Control GLP-1 Receptor Signaling in Pancreatic β-Cells. Diabetes 2017, 67 (3), 385–399. 10.2337/db17-0639.

(27) Girada, S. B.; Kuna, R. S.; Bele, S.; Zhu, Z.; Chakravarthi, N. R.; DiMarchi, R. D.; Mitra, P. Gαs Regulates Glucagon-Like Peptide 1 Receptor-Mediated Cyclic AMP Generation at Rab5 Endosomal Compartment. Molecular Metabolism 2017, 6 (10), 1173–1185. 10.1016/j.molmet.2017.08.002.

(28) Huang, X.; Dai, F. F.; Gaisano, G.; Giglou, K.; Han, J.; Zhang, M.; Kittanakom, S.; Wong, V.; Wei, L.; Showalter, A. D.; Sloop, K. W.; Stagljar, I.; Wheeler, M. B. The Identification of Novel Proteins That Interact With the GLP-1 Receptor and Restrain Its Activity. Molecular Endocrinology 2013, 27 (9), 1550–1563. 10.1210/me.2013-1047.

(29) Zhang, M.; Robitaille, M.; Showalter, A. D.; Huang, X.; Liu, Y.; Bhattacharjee, A.; Willard, F. S.; Han, J.; Froese, S.; Wei, L.; Gaisano, H. Y.; Angers, S.; Sloop, K. W.; Dai, F. F.; Wheeler, M. B. Progesterone Receptor Membrane Component 1 Is a Functional Part of the Glucagon-like Peptide-1 (GLP-1) Receptor Complex in Pancreatic β Cells* . Molecular & Cellular Proteomics 2014, 13 (11), 3049–3062. 10.1074/mcp.M114.040196.

(30) Liu, Y.; Batchuluun, B.; Ho, L.; Zhu, D.; Prentice, K. J.; Bhattacharjee, A.; Zhang, M.; Pourasgari, F.; Hardy, A. B.; Taylor, K. M.; Gaisano, H.; Dai, F. F.; Wheeler, M. B. Characterization of Zinc Influx Transporters (ZIPs) in Pancreatic β Cells. Journal of Biological Chemistry 2015, 290 (30), 18757–18769. 10.1074/jbc.M115.640524.

