## Supporting Information for "μMap-uAA: Photocatalytic proximity labeling targeted with single-residue precision"

### Supporting Figures

#### General procedures

##### General materials

All buffers and synthetic starting materials were used as received from commercial sources unless otherwise noted. Blue-light irradiation of samples was performed using a PennOC Photoreactor (PennOC, Pennsburg, PA, Model M2). Ascorbic acid (BP321-500) and ethanol (BP2818100) were purchased from Fisher Scientific (Pittsburgh, PA). Bovine serum albumin (BSA; A7906) and Eppendorf Protein LoBind tubes (Z666505) were purchased from Sigma-Aldrich. Sodium azide (14314) was purchased from Alfa Aesar (Haverhill, MA). RIPA buffer (89900), 1× DPBS (14190144), Pierce BCA Protein Assay Kit (23227), iBright Prestained Protein Ladder (LC5615), Trypsin-EDTA (Gibco, 25300054), TrypLE Express (Gibco, 12604021), Streptavidin Magnetic Beads (Pierce, 88816), trifluoroacetic acid (Optima grade), acetonitrile (Optima grade), water (Optima grade), acetic acid (Optima grade), urea (Pierce, Sequanal grade, 29700), and DTT (R0862) were purchased from Thermo Fisher Scientific. TBST (IBB-S18-581X) was purchased from Boston BioProducts (Ashland, MA). Axygen 1.5 mL Maximum Recovery tubes (MCT-150-LC) were purchased from Axygen Scientific (Union City, CA). Sodium hydroxide (567530), triethylammonium bicarbonate solution (1 M; 90360), hydroxylamine solution (50%; 438227), ammonium bicarbonate (LiChropur, Merck, 5438350), iodoacetamide (I1149), and cOmplete EDTA-free protease inhibitor cocktail (Roche, 11873580001) were purchased from Sigma-Aldrich. Criterion TGX 4–20% Tris-glycine polyacrylamide gel cassettes (5671094) and 4× Laemmli sample buffer (161-0747) were purchased from Bio-Rad (Hercules, CA). Biotin-$PEG_3$-diazirine (S2) was prepared as described previously.[4]

##### General Western Blot Protocol

Gel electrophoresis was performed using a Bio-Rad Criterion Vertical Electrophoresis Cell tank, Bio-Rad PowerPac Basic Power Supply, and Criterion TGX tris-glycine polyacrylamide gel cassettes (SDS/Tris). After electrophoresis, gels were transferred from precast cassettes to nitrocellulose membranes using an iBlot 2 gel transfer device (Thermo Fisher, IB21001, IB23001) and washed with water. The membranes were then immersed in REVERT total protein stain (Li-Cor, 926-11011) for 5 minutes. Excess stain was decanted, membranes washed with 6.7:30:63.3 AcOH:MeOH:$H_2O$ and imaged using a Li-Cor Odyssey CLx scanner in the 700 nm channel. The membranes were washed with water, then immersed in Odyssey Blocking

Buffer (Li-Cor, 927-50000) and incubated for 1 hour. The blocking solution was then decanted, and 35 mL of fresh blocking buffer containing 70 µL of Tween 20 was added. This mixture was rocked for 5 minutes. Afterwards, 1.5 µL of IRDye 800CW streptavidin (Li-Cor, 926-32230) was added and the mixture incubated for 1 hour. The blocking buffer was then decanted, and the membranes were washed with 1X TBST (3 x 5 min) and water before imaging via Li-Cor Odyssey CLx scanner in the 800 nm channel. Pixel densitometry was performed using Image Studio Lite V. 5.2 (Li-Cor). The streptavidin 800 channel pixel density was then divided by the total protein stain 700 channel pixel density to provide a normalized biotinylation signal for each protein band.

**General Proteomics Protocol**

Label-free data-independent acquisition (DIA) proteomics was performed on a Bruker timsTOF Pro 2 mass spectrometer coupled to a nanoElute LC system. For each sample, 1 µL of enriched peptide sample was injected onto a C18 trap column (PepMap, 5 µm particle size, 5 mm length, 300 µm internal diameter), followed by separation on a C18 analytical column (ReproSil AQ, 1.9 µm particle size, 100 mm length, 75 µm internal diameter). Peptides were eluted using a MeCN/water gradient at a column temperature of 40 °C with buffer A consisting of 0.1% formic acid in water and buffer B consisting of 0.1% formic acid in MeCN. The flow rate was 0.5 µL/min. The gradient started at 2% B, increased to 35% B over 20 min, increased to 95% B over 30 s, and was held at 95% B for 2.25 min. Scans were performed in positive-ion dia-PASEF mode over an m/z range of 100–1700, with a ramp time of 100 ms, accumulation time of 100 ms, duty cycle of 100%, ramp rate of 9.43 Hz, and MS averaging set to 1. Absolute thresholds were set to 5000 for mobility peaks and 10 for MS peaks.

The resulting raw data files (.d) were processed using DIA-NN v. 2.0 with the following parameters: trypsin/P digestion, up to 3 missed cleavages, up to 3 variable modifications, N-terminal methionine excision, methionine oxidation, N-terminal acetylation, cysteine carbamidomethylation, peptide length range of 7–30 amino acids, precursor charge range of 1–4, precursor m/z range of 300–1800, fragment ion m/z range of 200–1800, MS1 and MS2 mass accuracies set to 10 ppm, and precursor FDR set to 1%. The following DIA-NN settings were applied: "Use isotopologues," "MBR," "No shared spectra," and "Heuristic protein inference." An in silico spectral library was generated in DIA-NN from the UniProt human proteome FASTA file (UP000005640) using the "FASTA digest for library-free search/library generation" and "Deep learning-based spectra, RTs and IMs prediction" options, with all other parameters as described above.

The resulting matrix.pg files were processed in Perseus v. 2.0.7.0. Protein intensities were imported as main values, while the remaining descriptors were imported as categorical values. Intensities were $log_2$-transformed and annotated according to the corresponding experimental conditions. Data were normalized by median subtraction, and volcano plots were generated using a two-sided t-test for statistical analysis. Final volcano plots were prepared in GraphPad Prism 9. Gene ontology analysis performed by taking top enriched proteins and analyzing them via Metascape.[5]

**Cell culture**

HEK293T cells were obtained from the American Type Culture Collection (ATCC) and maintained in Dulbecco's modified Eagle medium (DMEM; Gibco, 11995065 and 31053036) supplemented with 10% fetal bovine serum (FBS; Gibco, 10437028) and 1% penicillin-streptomycin (Gibco, 15070063). Cells were cultured in 10-cm tissue culture dishes at 37 °C in a humidified incubator with 5% $CO_2$.

**Immunofluorescence Microscopy**

HEK293T cells were seeded into glass-bottom 8-well chamber slides (Thermo Fisher, 155409) pre-coated with poly-D-lysine (Cultrex, 3439-100-01) and allowed to recover overnight. µMap-uAA labeling was performed as described above. Following labeling, the solution was removed, and cells were washed twice with DPBS for 30 min each to remove unbound photocatalyst. Cells were then fixed with 4% paraformaldehyde in DPBS for 15 min at room temperature, washed once with DPBS, and permeabilized with 0.1% Triton X-100 in DPBS for 10 min. Samples were blocked for 1 h at room temperature in blocking buffer consisting of 3% BSA and 0.05% Tween-20 in DPBS. Primary antibodies, including anti-1D4 antibody (Santa Cruz Biotechnology, sc-57432) or anti-HA antibody (Invitrogen, 26183), were diluted in blocking buffer according to the manufacturer's recommended conditions and applied overnight at 4 °C. Samples were then washed three times with 0.05% Tween-20 in DPBS and incubated with secondary antibody solution in blocking buffer for 1 h at room temperature. Hoechst 33342 (1:10,000; Tocris, 5117) was used for nuclear staining. Goat anti-mouse Alexa Fluor 488 (1:1,000; Invitrogen, A11001) and goat anti-rabbit Alexa Fluor 555 (1:1,000; Invitrogen, A21428) were used to detect the corresponding primary antibodies, and Streptavidin Alexa Fluor 488 (1:2,000; Invitrogen, S11223) was used to detect biotinylation signal where appropriate. Samples were washed three times with 0.05% Tween-20 in DPBS and stored in DPBS before imaging. Images were acquired on a Nikon A1R-Si HD confocal microscope using NIS-Elements AR v4.60.00 software. Image processing was performed using ImageJ2/FIJI on Mac OS X.

**General procedure for uAA–μMap labeling of amber-mutant proteins in live HEK293T cells**

HEK293T cells were seeded into poly-D-lysine-coated 6 cm dishes (Gibco, A3890401) for each condition and grown to 40–60% confluency on the day of amber suppression. For each protein of interest (POI), amber suppression conditions were optimized separately in 6 cm dishes prior to labeling experiments, as expression efficiencies varied across constructs. Optimization parameters included the transfection reagent, the ratio of POI plasmid containing the amber codon mutation to the tRNA/aminoacyl-tRNA synthetase plasmid, and the expression time. On the day of labeling, cells were washed with complete DMEM (2 × 30 min) to remove unreacted endo BCN-L-lysine (**4**, SiChem, SC-8014). Cells were then incubated with iridium–tetrazine (**1**, 1 μM) in phenol red-free DMEM without FBS for 2 h, followed by an wash with complete DMEM (1 × 30 min, 37 °C). Next, cells were incubated with 500 μM biotin–diazirine (**S1**) in phenol red-free complete DMEM (Gibco, 31053028) for 30 min. After biotin–diazirine incubation, cells were directly subjected to 5 min irradiation using M2 photoreactors (Acceled) equipped with a 450 nm LED plate at 100% intensity (2.2 W output). Cells were then collected, washed with room temperature DPBS (Gibco, 14190136), pelleted, and stored at −80 °C until further use. Cell pellets were resuspended in RIPA lysis buffer (Thermo Fisher Scientific, 89901) and vortexed for 10–15 s three times, with 5 min cooling intervals on ice between each cycle. Samples were then sonicated using a Bioruptor Plus (Diagenode, B01020014) for 10 cycles (30 s on, 30 s off) at 4 °C and clarified by centrifugation at 18,000g for 15 min at 4 °C. The supernatants were transferred to fresh tubes, and protein concentrations were determined by bicinchoninic acid (BCA) assay (Thermo Fisher Scientific, 23227) using a BioTek Gen5 v2.05 plate reader. For streptavidin bead enrichment, 100 μL of lysate was added to a 1.5 mL LoBind tube containing Pierce streptavidin magnetic beads (Thermo Fisher Scientific, 88817) that had been pre-washed 2x with 1 mL RIPA buffer. Magnetic beads were added at a ratio of 1 μL beads per 10 μg total protein in the lysate. Samples were incubated overnight on a rotisserie at 4 °C, after which the beads were pelleted on a magnetic rack and the supernatant was removed. The beads were washed 3x with 500 μL 1% SDS in DPBS, 2x with 500 μL 1 M NaCl in DPBS, and 1x with 500 μL 10% EtOH in DPBS. The beads were then resuspended in 0.5 mL RIPA and transferred to a fresh 1.5 mL LoBind tube. After removal of the supernatant, the beads were resuspended in 50 μL elution buffer (6 M urea, 2 M thiourea, 30 mM biotin, 2% SDS in DPBS, 25% 4× Laemmli buffer with β-mercaptoethanol, pH 11.5). Samples were heated at 95 °C for 15 min with shaking at 700 rpm. The beads were pelleted on a magnetic rack, and 20 µL of the resulting supernatant was loaded onto a 4–20% Criterion TGX Precast SDS–PAGE gel (Bio-Rad, 5671094). Gels were run in freshly prepared Tris-glycine running buffer for 1 h at 160 V using a Criterion Cell and PowerPac system (Bio-Rad, 1656019).

For Western blot analysis, membranes were probed with the relevant epitope-tag primary antibody overnight at 4 °C. Membranes were then washed 3x with TBST (5 min each) and 3x with water, followed by incubation in blocking buffer containing secondary reagents (1:8000 dilution) for 1 h at room temperature. Secondary reagents included goat anti-mouse IRDye 680CW and streptavidin IRDye 800LT (LI-COR). Membranes were washed 3x with TBST (5 min each) and 3x with water prior to imaging on an Odyssey CLx imaging system (LI-COR, 9140).

For proteomics experiments, the same streptavidin enrichment procedure was used, except that each condition (± POI expression) was prepared in triplicate and the streptavidin beads were processed according to the General Proteomics Procedure prior to addition of the elution buffer.

#### Figure S1. µMap-uAA with NONO and ACTB

##### Figure S1.A: β-Actin pulse-chase assay for live-cell Ir–tetrazine engagement

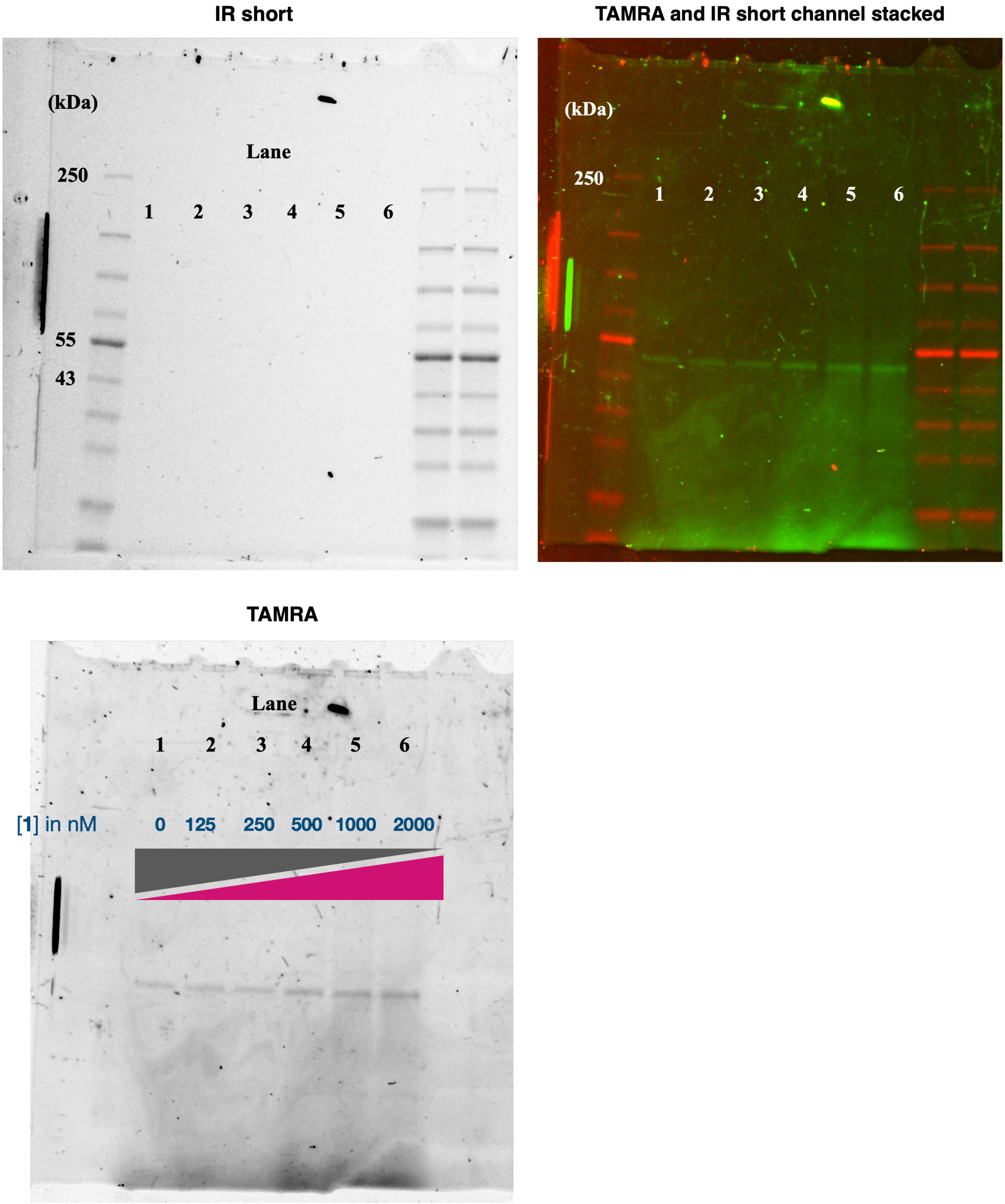

β-Actin was selected as an initial intracellular test protein for µMap-uAA because a β-actin K118TAG construct has been previously reported for site-specific incorporation of clickable noncanonical amino acids, including BCN-lysine (**4**).[1] This established β-actin K118TAG system_provided a well-defined starting point to evaluate whether genetically encoded BCN could support live-cell installation of Iridium-tetrazine (**1**). The amber suppression conditions optimized for β-actin served as the experimental basis for subsequent μMap-uAA labeling of additional intracellular and membrane protein targets described in this study.

HEK293T cells were seeded in 6-well Nunc cell-culture plates (Thermo Fisher Scientific, cat. no. 140675) that had been pre-coated with poly-D-lysine. For coating, 1 mL of room-temperature poly-D-lysine solution was added to each well and incubated for 1 h at room temperature. The solution was then aspirated, and wells were washed twice with DPBS prior to cell seeding. On the day of amber suppression, the culture medium was aspirated and replaced with phenol red-free DMEM containing **4** at a final concentration of 0.5 mM. 6 μg β-actin K118TAG plasmid, 6 μg M21 tRNA/aaRS plasmid, and 24 μL P3000 reagent were diluted in 1200 μL Opti-MEM and incubated for 5 min at room temperature to generate the plasmid master mix. In a separate tube, 36 μL Lipofectamine 3000 was diluted in 1200 μL Opti-MEM. The plasmid master mix was then added to the Lipofectamine 3000 solution and incubated for 10 min at room temperature. The resulting transfection mixture was added dropwise to cells at 400 μL per well.

After 36 h, cells were washed twice by incubation with 2 mL complete DMEM for 1 h at 37 °C 5% $CO_2$ per wash to remove residual free **4** from the culture medium. Cells were then incubated with **1** in phenol red-free DMEM without FBS at the indicated concentrations (2000, 1000, 500, 250, 125, or 0 nM) for 2 h at 37 °C 5% $CO_2$. Following live-cell incubation with **1**, cells were washed twice with DPBS and lysed in 100 μL RIPA buffer supplemented with protease inhibitor cocktail.

Lysate protein concentrations were determined by BCA assay and equalized prior to the chase reaction. To label remaining unreacted BCN-containing β-actin, Tetrazine–PEG4–TAMRA (BroadPharm, BP-22940, **S2**) was added to each lysate at a final concentration of 1 μM. Samples were incubated for 1 h in the dark with rotation. Following incubation, 4× Laemmli sample buffer was added, and samples were analyzed by SDS-PAGE. Gels were imaged by in-gel fluorescence to detect TAMRA-labeled β-actin. A decrease in TAMRA signal with increasing [**1**] indicates dose-dependent engagement of genetically encoded BCN in β-actin by **1** in living cells.

**Table S1**: Iridium-tetrazine (**1**) and **S2** concentrations used for the β-actin pulse-chase assay.

| Lane | [**1**] nM | [**S2**] added to lysates, nM |
|---|---|---|
| 1 | 2000 | 1000 |

| | | |
|---|---|---|
| 2 | 1000 | 1000 |
| 3 | 500 | 1000 |
| 4 | 250 | 1000 |
| 5 | 125 | 1000 |
| 6 | 0 | 1000 |

**Figure S1.B: µMap-uAA of NONO-258TAG**

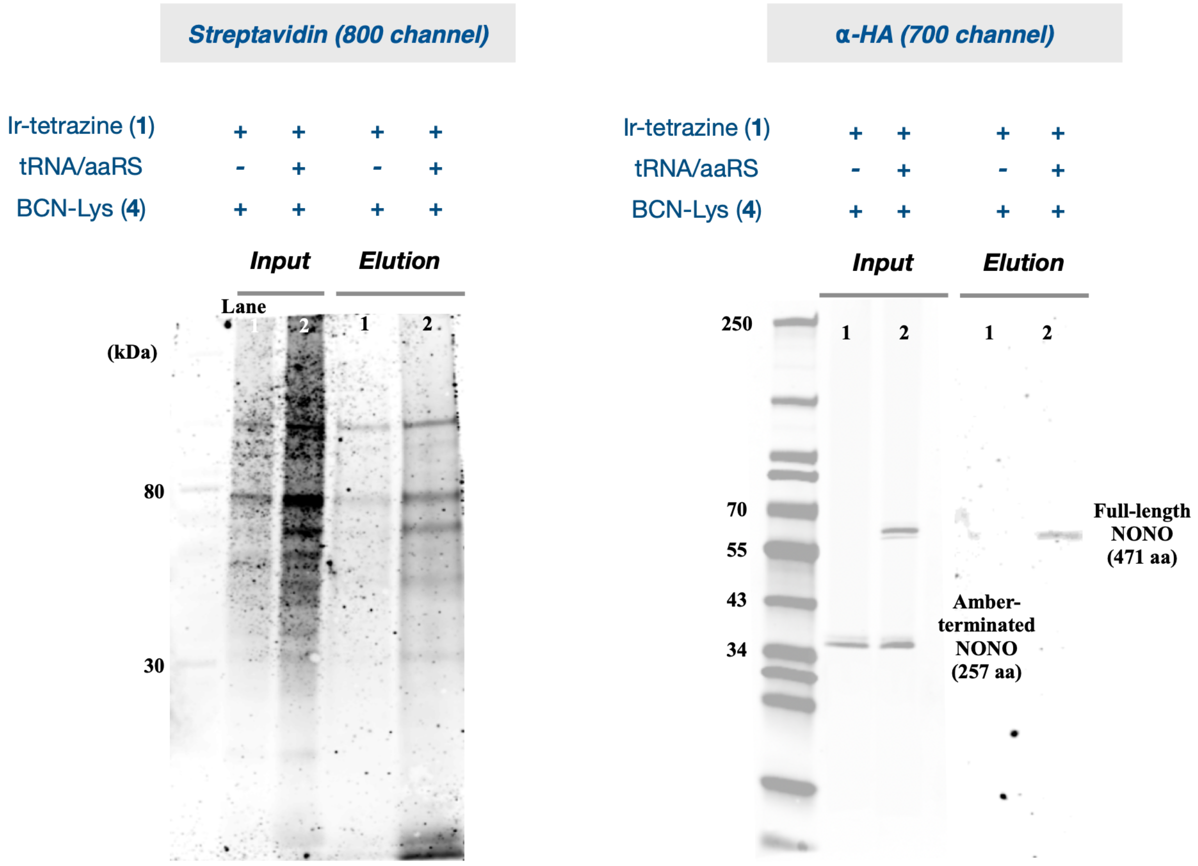

HEK293T cells were seeded in two 6 cm plates one day prior to transfection. One plate was transfected with plasmids encoding N-terminally HA-tagged NONO-258TAG (1 µg) and the M21 tRNA/aaRS amber suppression machinery (2 µg) using P3000 reagent (6 µL) and Lipofectamine 3000 (9 µL) according to the manufacturer's protocol (Lane 2). The second plate was transfected with N-terminally HA-tagged NONO-258TAG plasmid only (1 µg), using P3000 reagent (2 µL) and Lipofectamine 3000 (3 µL), and served as the no-amber-suppression control (Lane 1).

After 36 h, both plates were subjected to the µMap-uAA labeling protocol, harvested, and lysed in RIPA buffer supplemented with protease inhibitor cocktail. Lysate protein concentrations were determined by BCA assay and equalized prior to streptavidin enrichment. A portion of each lysate was reserved as input. Input lysates and streptavidin-enriched samples were analyzed by western blotting using streptavidin 800 nm and anti-HA 700 nm channels.

Because the HA tag is positioned at the N-terminus, both the amber-terminated HA–NONO product and the full-length, BCN-incorporated HA–NONO product are detected in the input lysate. In the streptavidin 800 nm channel, global biotinylation level was higher in the presence of the M21 tRNA/aaRS machinery, supporting click-dependent labeling. Following streptavidin enrichment, only full-length HA–NONO was detected by anti-HA western blot, consistent with selective enrichment of the amber-suppressed, BCN-containing NONO product.

#### Figure S2: Click-dependent μMap-uAA

##### Steady-State Emission Measurements and *in vitro* BSA labeling

Steady-state fluorescence emission spectra were recorded on an Agilent Cary Eclipse fluorescence spectrophotometer using 10 mm quartz cuvettes. All measurements were conducted at room temperature.

A 1 mM stock solution of iridium–tetrazine photocatalyst was prepared in DMSO. For emission measurements of the unreacted catalyst, 40 μL of the 1 mM Ir–tetrazine stock solution was diluted into 3.96 mL acetonitrile (MeCN) to give a final concentration of 10 μM Ir–tetrazine in a total volume of 4.00 mL.

For click-reacted samples, Ir–tetrazine and BCN–OH (**3**, BroadPharm, Cat. BP-26285) were first allowed to react prior to dilution for fluorescence measurement. Briefly, 25 μL of a 2 mM Ir–tetrazine solution in DMSO was combined with 25 μL of **3** solution at the desired concentration to achieve the specified molar equivalents relative to **1**. The reaction mixture was allowed to react at room temperature to form the corresponding pyridazine adduct. For the HRMS analysis of click conversion, please refer to the below section.

Following the click reaction, 40 μL of the reaction mixture was diluted into 3.96 mL MeCN, affording a final Ir concentration of 10 μM in 4.00 mL total volume for fluorescence analysis.

Emission spectra were collected with excitation at $\lambda_{ex}$ = 400 nm, and emission was recorded from 400–600 nm.

**Table S2**: Reaction and dilution conditions for fluorescence analysis of **1** following click reaction with **3**

| sample | **1** stock (2 mM) | BCN-OH stock | Reaction mixture prepared | Aliquot used | MeCN added | Final Ir concentration |
|---|---|---|---|---|---|---|
| **1** alone | 40 µL of 1mM **1** | N/A | N/A | 40 µL | 3960 µL | 10 µM |
| 0.2 eq. **3** | 25 µL (2 mM **1**) | 25 µL **3** (0.4 mM) | 50 µL | 40 µL | 3960 µL | 10 µM |
| 0.8 eq. **3** | 25 µL (2 mM **1**) | 25 µL **3** (1.6 mM) | 50 µL | 40 µL | 3960 µL | 10 µM |
| 1.0 eq. **3** | 25 µL (2 mM **1**) | 25 µL **3** (2.0 mM) | 50 µL | 40 µL | 3960 µL | 10 µM |

**HRMS analysis of click conversion.**

**1** was incubated with **3** at the indicated molar equivalents in DMSO for 1 h in the dark at room temperature, and the reaction mixtures were analyzed by HRMS. Reaction progress was evaluated by integrating the extracted-ion chromatograms corresponding to the **1** (**m/z 1966.7**) and the clicked pyridazine adduct (**2**, **m/z 2088.8**). The relative ratio of product-to-starting-material signal was used as a semi-quantitative measure of conversion. As the molar equivalence of **3** increased, the signal corresponding to the starting material decreased while the product signal increased, consistent with progressive click ligation. Because ionization efficiencies of the precursor and product were not independently calibrated, these values are reported as relative LC–MS responses rather than absolute percent conversion.

**Figure S2.A: HRMS analysis of click conversion**

1 alone

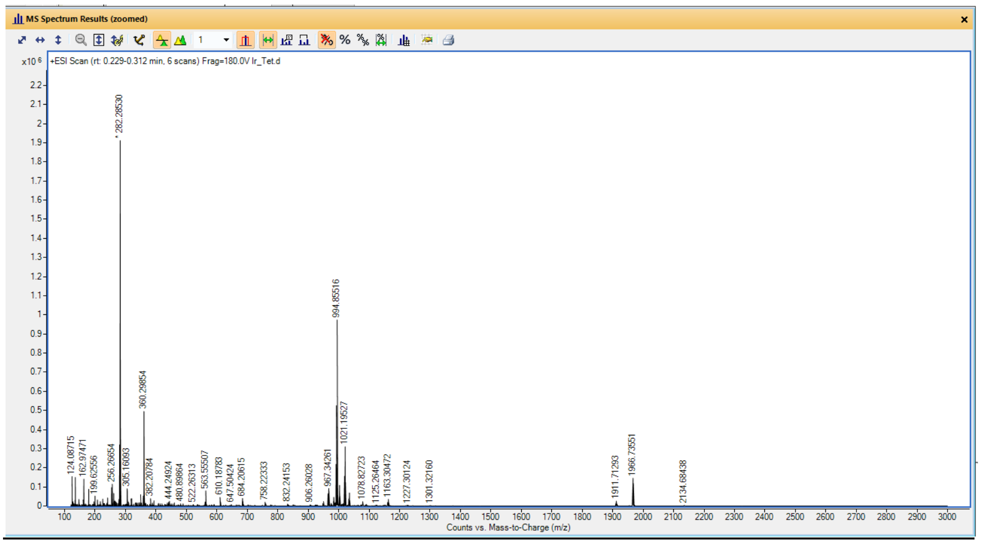

#### With 0.2 molar equivalence of **3**

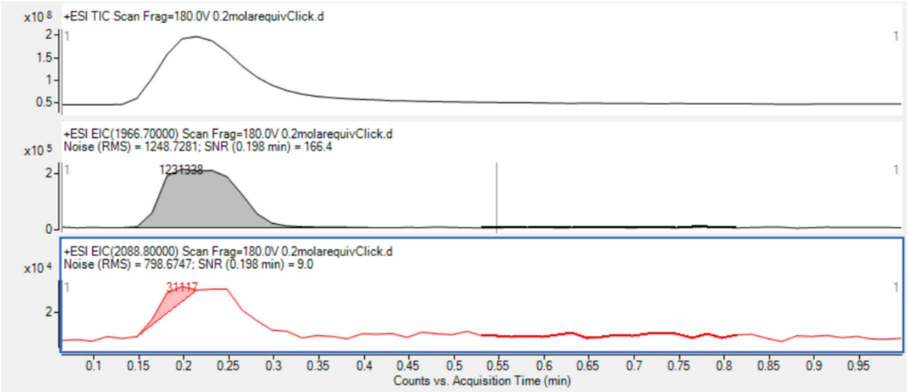

#### With 0.8 molar equivalence of **3**

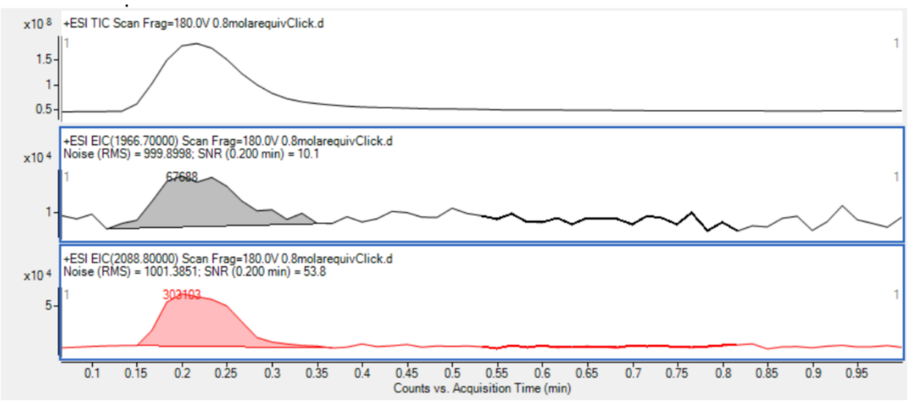

ESI scan of **2** (full conversion)

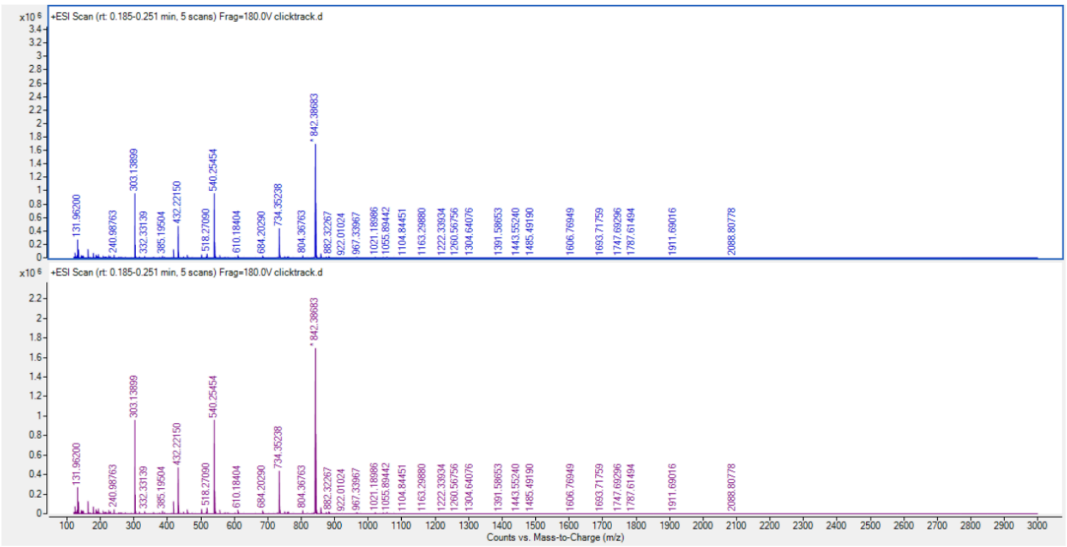

**Biotinylation: Catalytic Diazirine Sensitization and BSA Labeling**

**1** was combined with biotin-diazirine (**S1**) and BSA in DPBS to afford reaction mixtures with 100 µL total solution volume and desired component concentrations (see table below). These samples were then either placed in the dark, irradiated with UV (365 nm) light, or irradiated with visible (450 nm) light in a biophotoreactor for 10 minutes at 100% intensity. 30 µL samples were then removed, combined with 10 µL of 4x reducing Laemmli sample buffer (5% ß-mercaptoethanol), vortexed, and heated at 95 °C for 10 minutes. 10 µL of each sample was then analyzed by Western blot according to the Western Blot General Protocol described above.

**Table S3**: reaction conditions for BSA biotinylation using biotin-diazirine with **1** and **2**

| Lane | [Biotin-Diazirine] (µM) | [Ir] (µM) | | [BSA] (µM) | Light |
|---|---|---|---|---|---|
| | | [**1**] | [**2**] | | |
| 1 | 100 | 0.0 | | 10.0 | None |
| 2 | 100 | 0.0 | | 10.0 | 365 nm |
| 3 | 100 | 0.0 | | 10.0 | 450 nm |
| 4 | 100 | 2.5 | 0.0 | 10.0 | 450 nm |
| 5 | 100 | 0.0 | 2.5 | 10.0 | 450 nm |
| 6 | 100 | 6.0 | 0.0 | 10.0 | 450 nm |
| 7 | 100 | 0.0 | 6.0 | 10.0 | 450 nm |
| 8 | 100 | 10.0 | 0.0 | 10.0 | 450 nm |
| 9 | 100 | 0.0 | 10.0 | 10.0 | 450 nm |

**Figure S2.B:** Representative western blots for *in vitro* BSA labeling with **1** and **2**

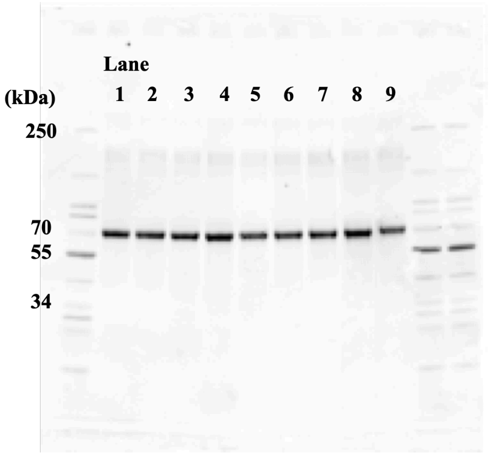

700 channel (total protein).

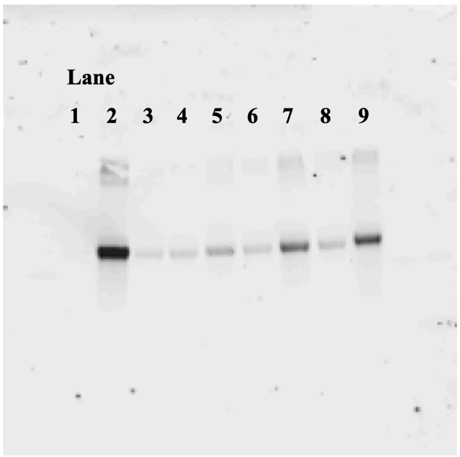

800 channel (streptavidin)

**Figure S2.C:** Densitometric quantification of *in vitro* BSA biotinylation with 1 and 2

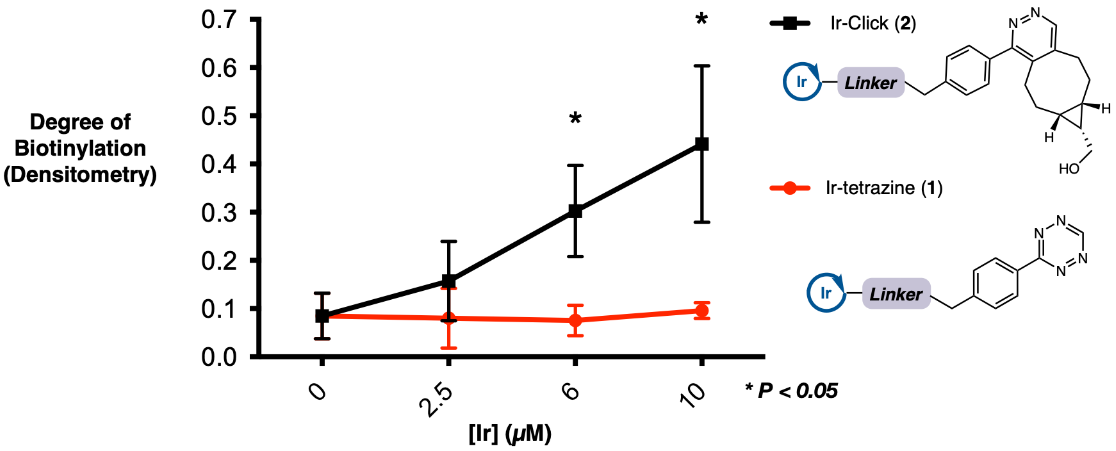

**Results**: The extent of protein biotinylation increases with increasing catalyst concentration for Ir–Click, whereas minimal change is observed for Ir–Tetrazine, consistent with reaction-gated activation of photocatalytic labeling.

**Panel**: Normalized intensity values were obtained by densitometry analysis of Western blot bands using ImageJ (Fiji). Streptavidin (800 channel) signal was normalized to total protein signal. Error bars represent standard deviation (n = 3).

<u>Assessment of the inner filter effect</u>

**Figure S2.D**: Evaluation of inner-filter effects on photocatalytic BSA biotinylation

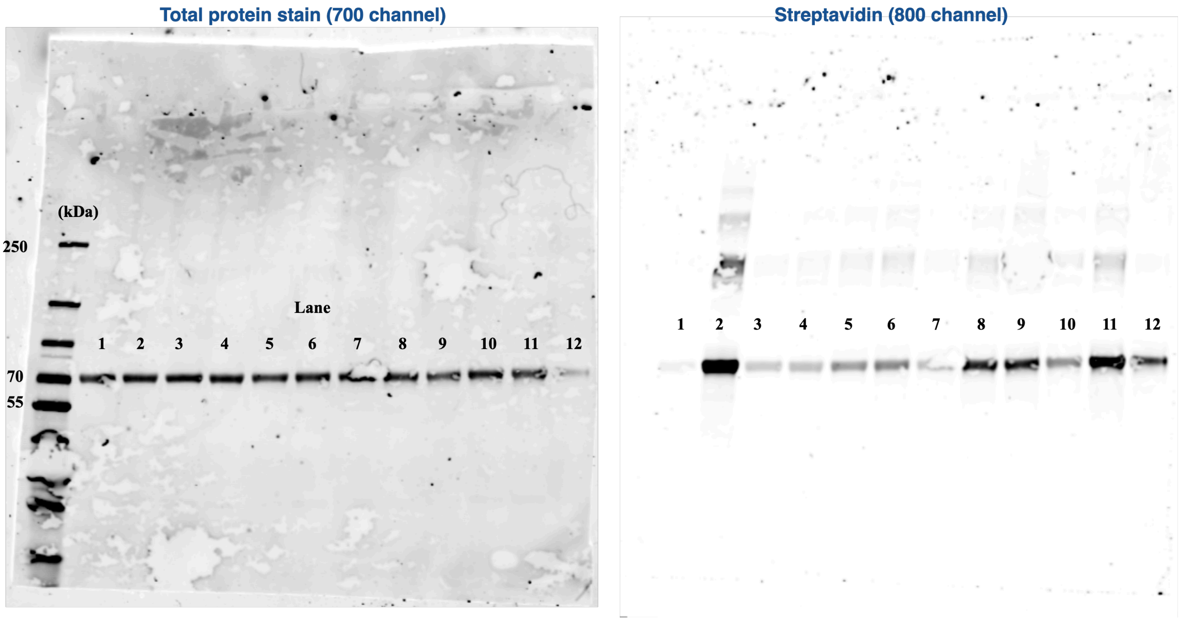

**Table S4**: reaction conditions for BSA biotinylation using biotin-diazirine with **1**, **2** and mixture of **1** and **2**

| Lane | [Biotin-Diazirine] (μM) | [Ir] (μM) | | [BSA] (μM) | Light |
|---|---|---|---|---|---|
| | | [**1**] | [**2**] | | |
| 1 | 100 | 0.0 | | 10.0 | None |
| 2 | 100 | 0.0 | | 10.0 | 365 nm |
| 3 | 100 | 0.0 | | 10.0 | 450 nm |
| 4 | 100 | 2.5 | 0.0 | 10.0 | 450 nm |
| 5 | 100 | 0.0 | 2.5 | 10.0 | 450 nm |
| 6 | 100 | 2.5 | 2.5 | 10.0 | 450 nm |
| 7 | 100 | 6.0 | 6.0 | 10.0 | 450 nm |
| 8 | 100 | 0.0 | 6.0 | 10.0 | 450 nm |
| 9 | 100 | 6.0 | 6.0 | 10.0 | 450 nm |
| 10 | 100 | 10.0 | 0.0 | 10.0 | 450 nm |
| 11 | 100 | 0.0 | 10.0 | 10.0 | 450 nm |
| 12 | 100 | 10.0 | 10.0 | 10.0 | 450 nm |

To determine whether increased iridium concentration attenuated photocatalytic labeling through an inner-filter effect, BSA was irradiated with blue light in the presence of Ir–click (**2**) either alone (lanes 5, 8, and 11) or together with an equimolar concentration of Ir–tetrazine (**1**) (lanes 6, 9, and 12). Thus, the mixed samples contained twice the total concentration of iridium while maintaining the same concentration of photoactive Ir–click (**2**). Comparable BSA biotinylation was observed between the Ir–click-only and mixed-catalyst conditions at each concentration tested, indicating that the increased total iridium concentration did not measurably affect photocatalytic labeling under these conditions.

#### Figure S3: Proof-of-concept with CCR5

**Figure S3A: Validation of amber suppression at CCR5 residues F96TAG and L352TAG under the final optimized transfection conditions.**

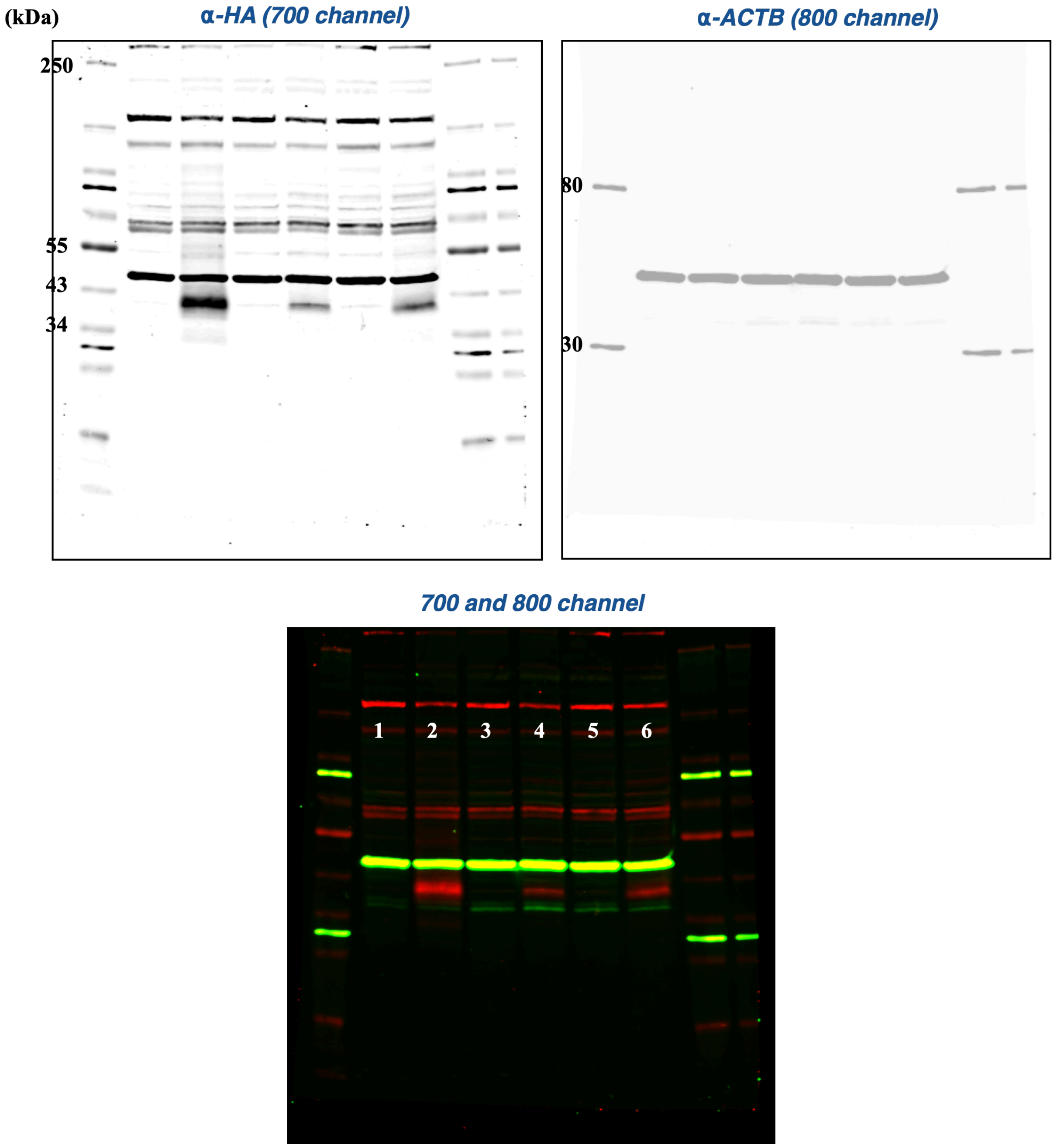

HEK293T cells were seeded in 6-well Nunc cell-culture plates (Thermo Fisher Scientific, cat. no. 140675) one day prior to transfection. The following day, cells were transfected with plasmids encoding the indicated CCR5 constructs, with or without the M21 tRNA/aaRS amber suppression machinery, in the presence of BCN–lysine (**4,** 0.5 mM final concentration in medium). Amber suppression and transfection conditions were optimized prior to this 6-well validation assay using a matrix-based optimization approach. Parameters evaluated included total plasmid loading per well, transfection reagent identity, transfection duration, and the ratio of the protein of interest amber mutant plasmid to the M21 tRNA/aaRS amber suppression plasmid. The optimized transfection conditions used for the 6-well amber suppression validation assay are summarized in Table S5.

For CCR5 F96* transfection, plasmid DNA encoding the indicated CCR5 F96TAG construct, without (well 3) or with (well 4) the M21 tRNA/aaRS plasmid, was diluted in Opti-MEM together with P3000 reagent (2xDNA) to a final volume of 400 μL. In a separate tube, Lipofectamine 3000 (3X DNA) was diluted in 200 μL Opti-MEM. After 5 min incubation at room temperature, the mastermix was combined with the Lipofectamine 3000 mixture and incubated for an additional 10 min at room temperature. The resulting transfection mixture was then added dropwise to the corresponding well.

For CCR5 L352* transfection, plasmid DNA encoding the indicated CCR5 construct, without (well 5) or with (well 6) the M21 tRNA/aaRS plasmid, was diluted in 300 μL Opti-MEM. FuGENE HD was added at a 3:1 FuGENE HD:DNA ratio, and the mixture was incubated for 5-10 min at room temperature before being added dropwise to cells.

After 36 h of expression, cells were harvested and lysed in 1x RIPA buffer supplemented with protease inhibitor cocktail. Lysates were clarified by centrifugation, and protein concentrations were determined by BCA assay. Equal amounts of total protein were separated by SDS-PAGE and analyzed by western blotting using an anti-HA antibody to detect HA-tagged CCR5 constructs. β-Actin was used as a loading control.

Expression of CCR5 amber mutants was observed only when the amber mutant construct was co-transfected with the M21 tRNA/aaRS machinery in the presence of **4**, consistent with successful amber suppression.

**Table S5**: Optimized transfection conditions for CCR5 amber suppression validation.

| Lane | CCR5 construct | **4** | tRNA/aaRS | Transfection condition |
|---|---|---|---|---|
| 1 | N/A | + | - | |
| 2 | CCR5 wt (1 μg) | + | - | |
| 3 | CCR5 F96* (1 μg) | + | - | P3000: 2 μL<br>Lipo3000: 3 μL |
| 4 | CCR5 F96* | + | + | P3000: 4 μL |

| | | | | |
|---|---|---|---|---|
| | (1 µg) | | (1 µg) | Lipo3000: 6 µL |
| 5 | CCR5 L352* (1 µg) | + | - | Fugene HD: 3 µL |
| 6 | CCR5 L352* (1 µg) | + | + (1 µg) | Fugene HD: 6 µL |

**Figure S3.B.: Dose-dependent in-lysate labeling of BCN-containing CCR5 F96* with S2.**

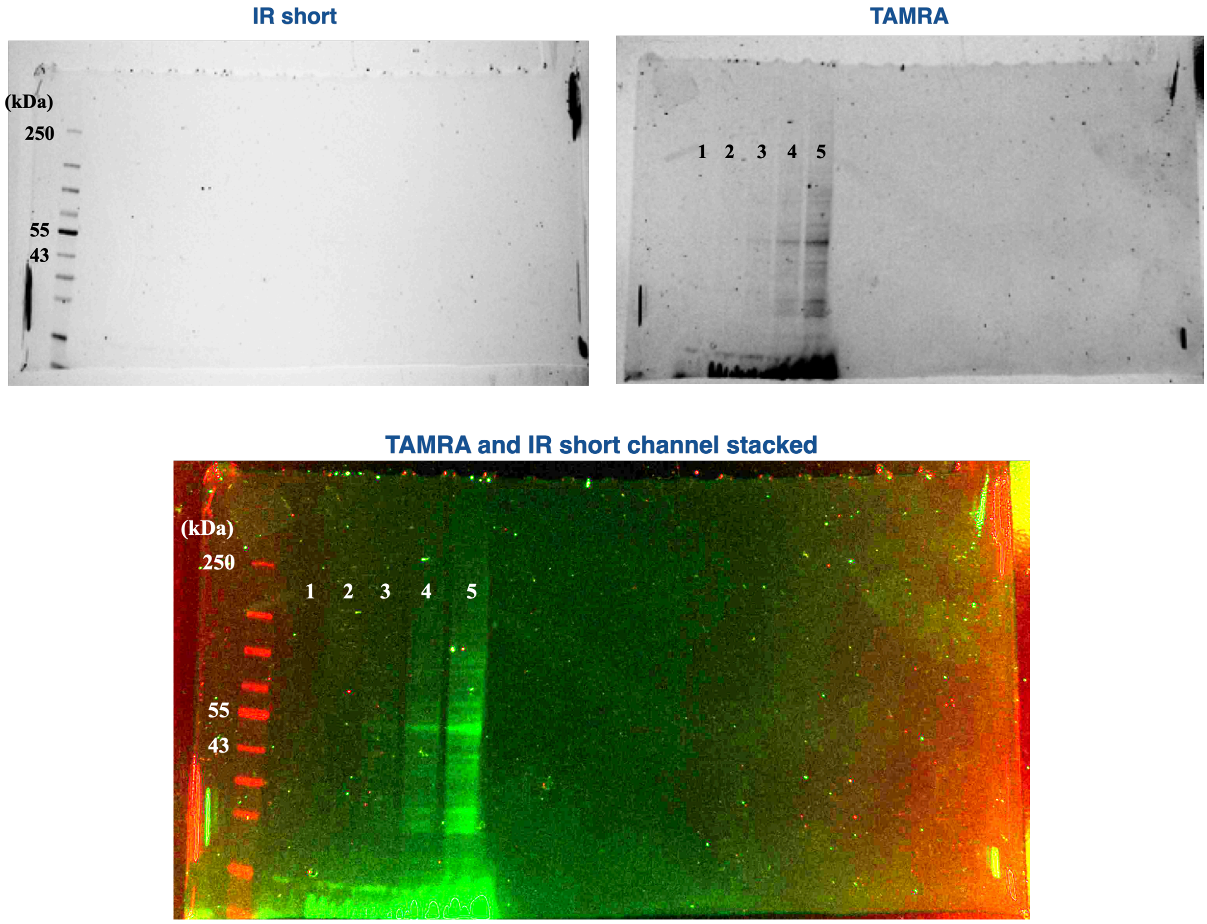

**Figure S3.C: Western blot validation of sample loading for S2 in-lysate labeling of CCR5 F96***

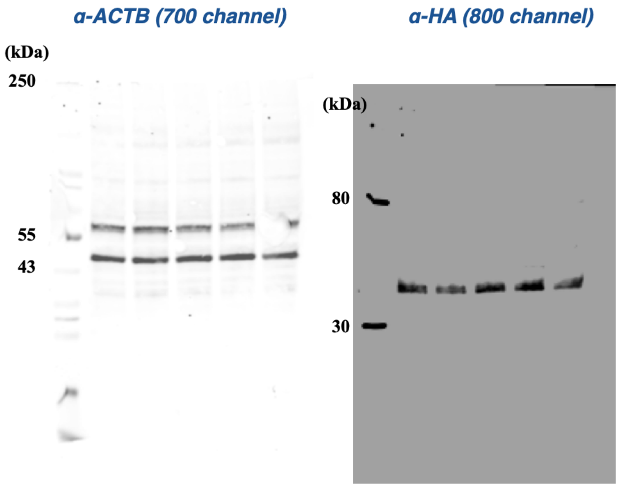

**Figure S3.D: Dose-dependent in-lysate labeling of BCN-containing CCR5 L352* with S2.**

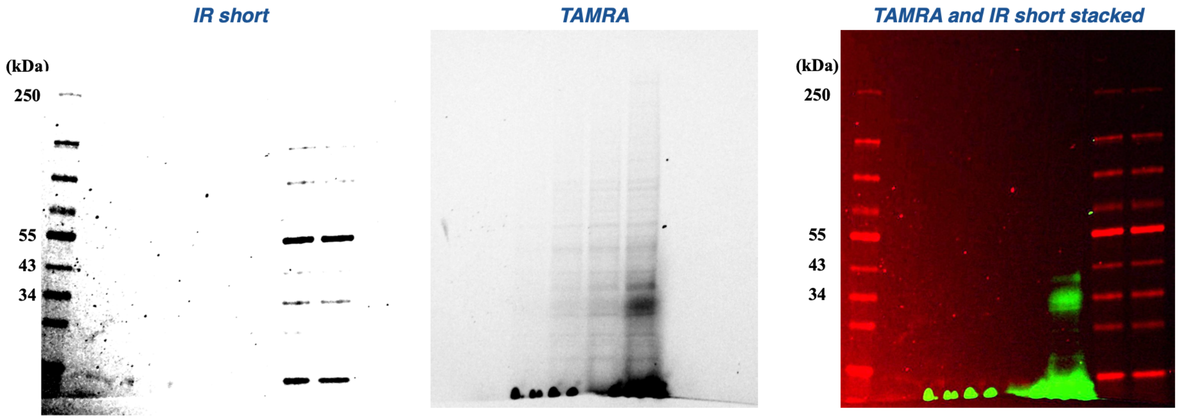

**Figure S3.E: Western blot validation of sample loading for S2 in-lysate labeling of CCR5 L352***

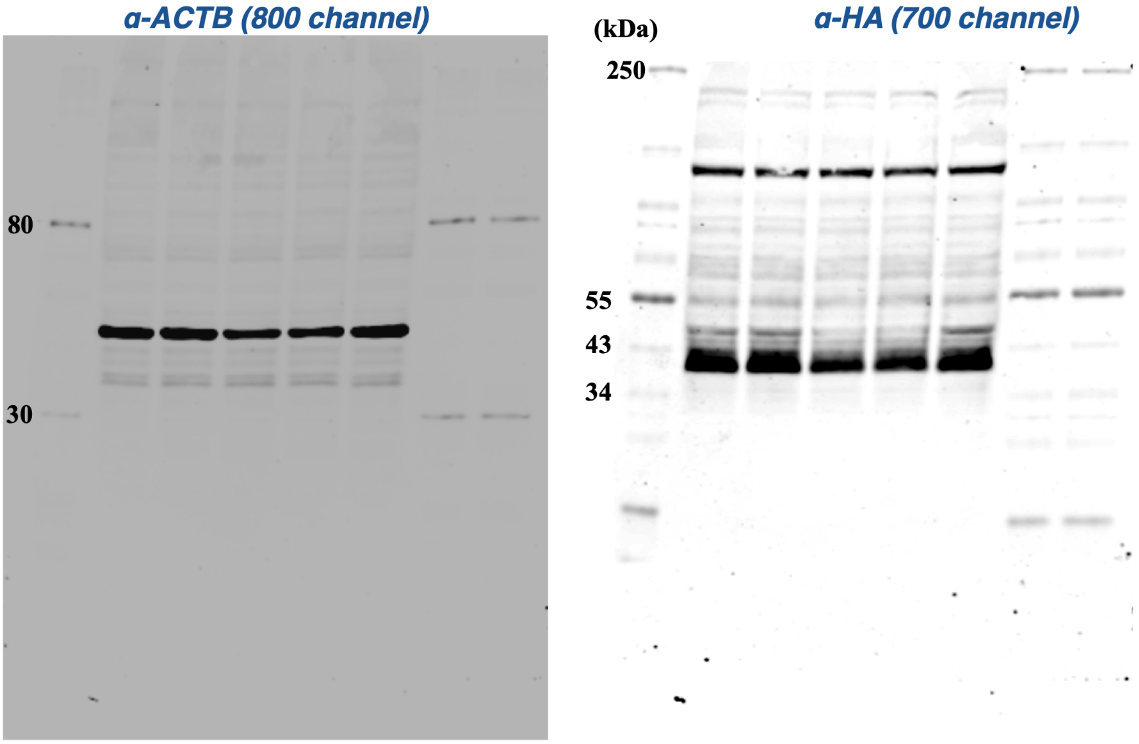

HEK293T cells were seeded in a 6 cm plate and transfected with plasmids encoding CCR5 F96TAG-HA (1 μg) and the orthogonal tRNA/tRNA synthetase pair (1 μg) using Lipofectamine 3000 (6 μL) and P3000 reagent (4 μL) according to the manufacturer's protocol (Figure S3B, S3C). Amber suppression with L352TAG is in Figure S3D, S3E.

Cells were harvested and lysed in 120 μL 1× RIPA buffer supplemented with protease inhibitor cocktail. Lysates were clarified by centrifugation and divided into five equal aliquots (20 μL each). Tetrazine–PEG4–TAMRA (BroadPharm, BP-22940, **S2**) was dissolved in DMSO to prepare 10, 25, 50, and 100 μM stock solutions. Lysate aliquots were treated with 0.2 μL of each stock solution to give final **S2** concentrations of 0, 100, 250, 500, and 1000 nM (lane 1 to 5 respectively); a control aliquot was treated with 0.2 μL DMSO. Samples were incubated for 1 h at room temperature in the dark.

Following incubation, 6.7 μL of 4× Laemmli sample buffer was added to each sample, and samples were briefly vortexed (10 s) without heating prior to SDS–PAGE. Heating was avoided to preserve TAMRA fluorescence and to minimize multimer formation during GPCR analysis. Proteins were resolved by SDS–PAGE, and the same gel was first imaged by in-gel fluorescence scanning in the TAMRA channel and for total protein ladder in the IR short channel, then directly transferred to a membrane for Western blot analysis (Figure S3C, S3E). Membranes were probed with anti-HA antibody to detect CCR5 expression and anti-ACTB antibody as a loading control. Overlay of the in-gel TAMRA band with the HA-positive CCR5 band confirmed receptor labeling. A dose-dependent increase in TAMRA fluorescence was observed at the molecular weight corresponding to CCR5, while HA signal remained consistent across samples, supporting successful incorporation of the BCN-unnatural amino acid at F96* and L352* and its reactivity toward tetrazine.

**Figure S3.F: Plasma membrane localization of CCR5 F96* and L352* following BCN incorporation**

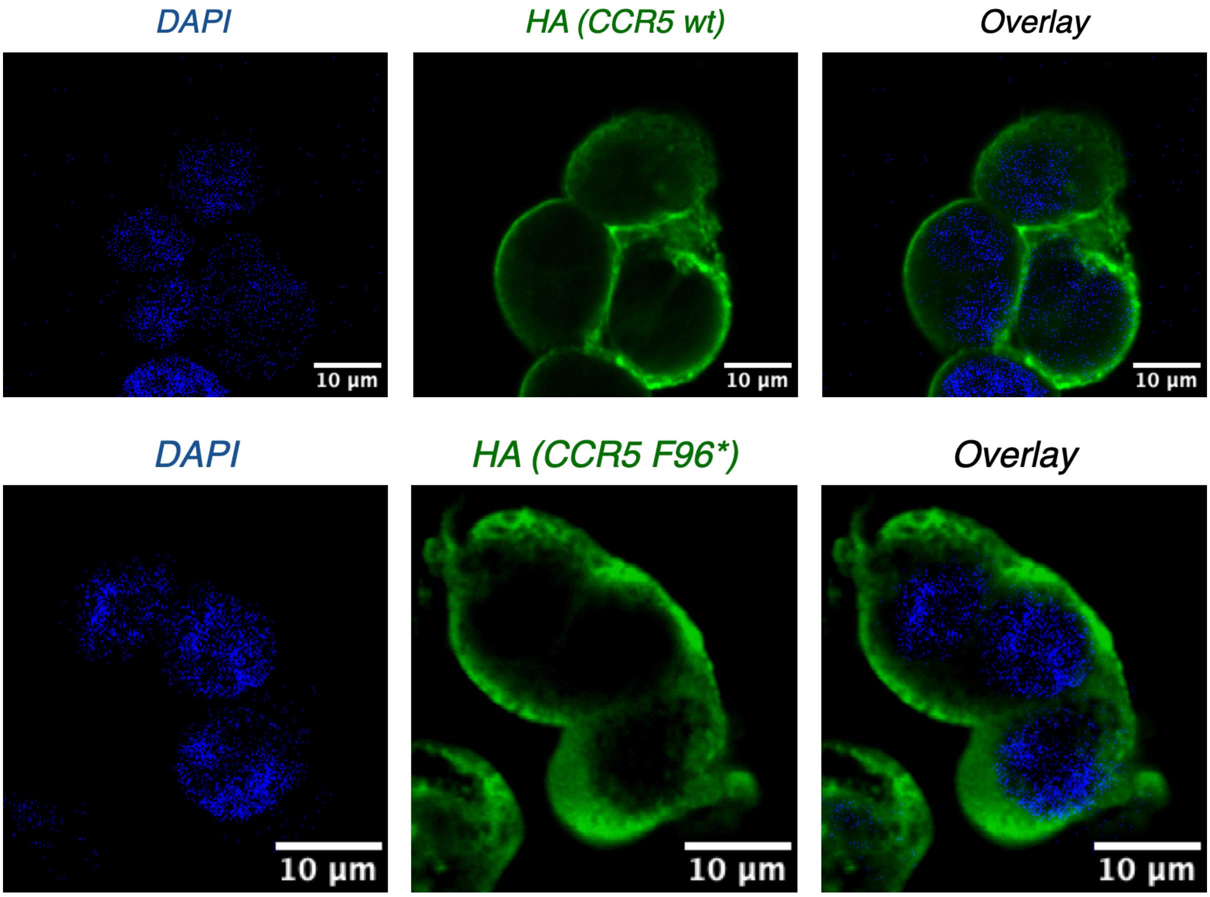

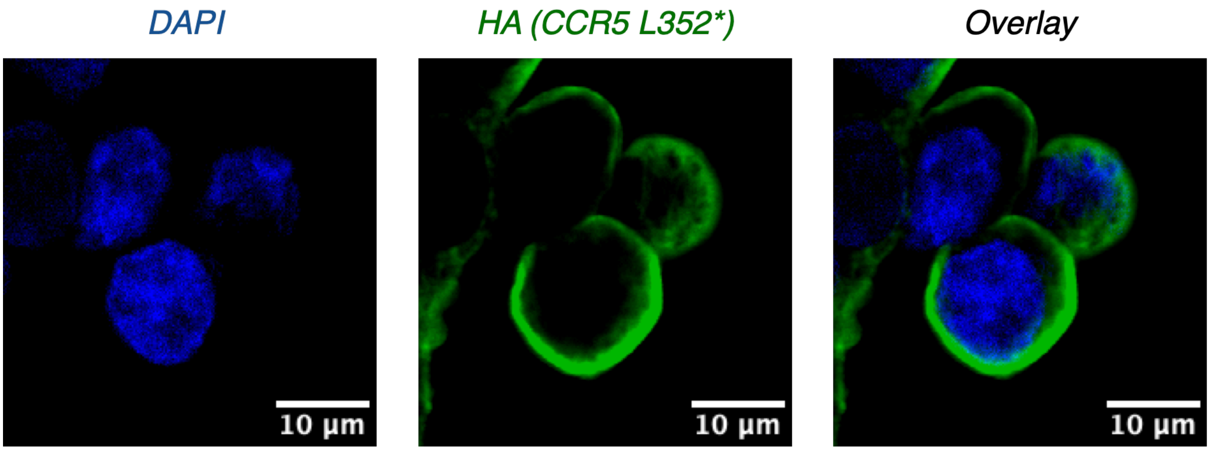

Representative confocal fluorescence images of HEK293T cells expressing HA-tagged wild-type CCR5, CCR5 F96*, or CCR5 L352* under amber-suppression conditions in the presence of BCN–Lys. CCR5 localization was visualized using mouse anti-HA antibody (Invitrogen, Cat. No. 26183; 1:1,000), and nuclei were stained with DAPI. BCN–Lys incorporation at either position retained plasma membrane localization comparable to wild-type CCR5. Scale bars, 10 µm.

**Figure S3.G: Confocal assay with CCR5 F96* amber suppression with BCN**

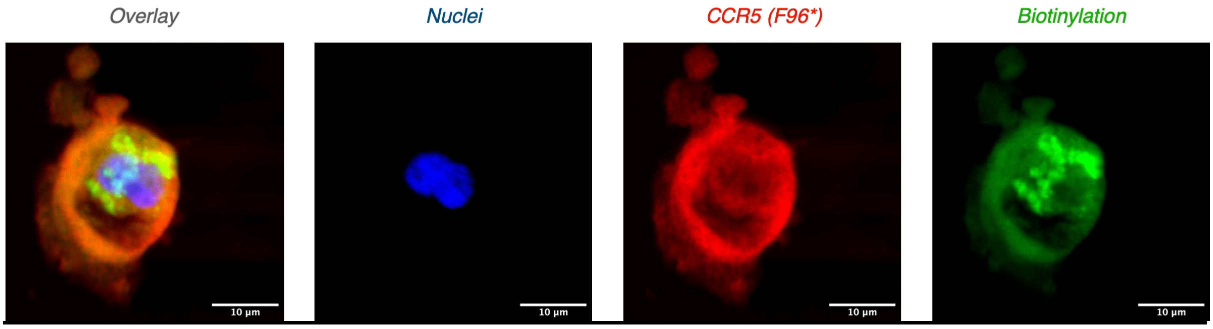

Representative confocal fluorescence images of HEK293T cells expressing HA-tagged CCR5 F96* following µMap–uAA labeling. CCR5 was detected using rabbit anti-HA antibody (Abcam, ab9110), biotinylated proteins were visualized using Alexa Fluor 488-conjugated streptavidin (Invitrogen, S11223), and nuclei were stained with DAPI. Biotinylation exhibited a predominantly peripheral pattern consistent with photocatalyst installation at the extracellular F96* site. Scale bars, 10 µm.

**Figure S3.H: Western blot analysis of CCR5 F96* labeling and enrichment**

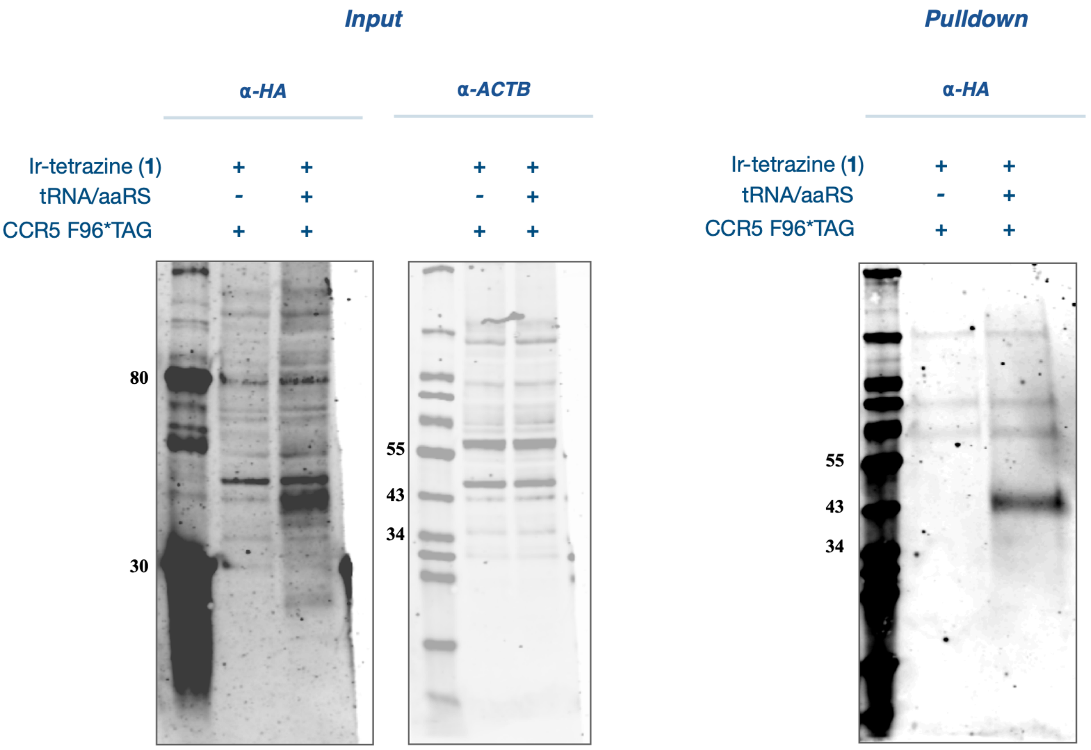

**Figure S3.I: Western blot analysis of CCR5 L352* labeling and enrichment**

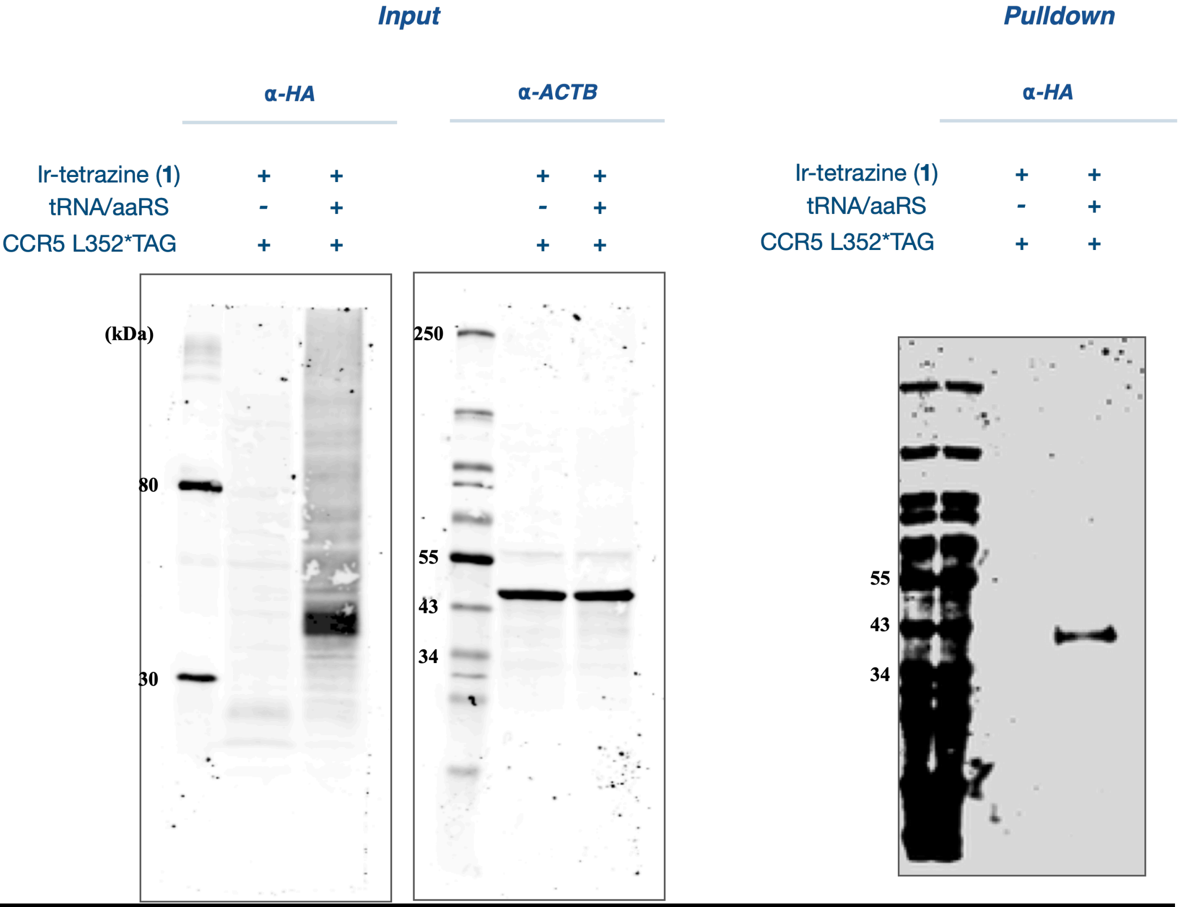

HEK293T cells expressing HA-tagged CCR5 F96*TAG (Figure S3.H) or CCR5 L352*TAG (Figure S3.I) were treated with **1** (1 µM) in the absence or presence of the orthogonal tRNA/aaRS amber suppression machinery. Input lysates were analyzed by western blot using rabbit anti-HA antibody (Abcam, ab9110), with ACTB as an input loading control (Cell Signaling Technology, 8H10D10). Streptavidin-enriched pulldown eluates were analyzed by western blot using anti-HA antibody (Invitrogen, 26183).

#### Figure S4: CCR5 validation

##### Figure S4A: F96* extracellular µMap-uAA identifies maraviroc-sensitive CCR5-proximal proteins

Procedure for Maraviroc-treatment during µMap-uAA labeling

For maraviroc-treated samples, µMap-uAA labeling was performed as described above with the following modification. Following 2 h incubation with iridium–tetrazine photocatalyst, the medium was aspirated and replaced with phenol red-free complete DMEM containing 100 nM maraviroc (MedChemExpress, HY-13004). Cells were incubated for 30 min at 37 °C and 5% CO2. The medium was then replaced with phenol red-free complete DMEM containing 500 µM biotin–diazirine and 100 nM maraviroc, and cells were incubated for an additional 30 min prior to blue-light irradiation.

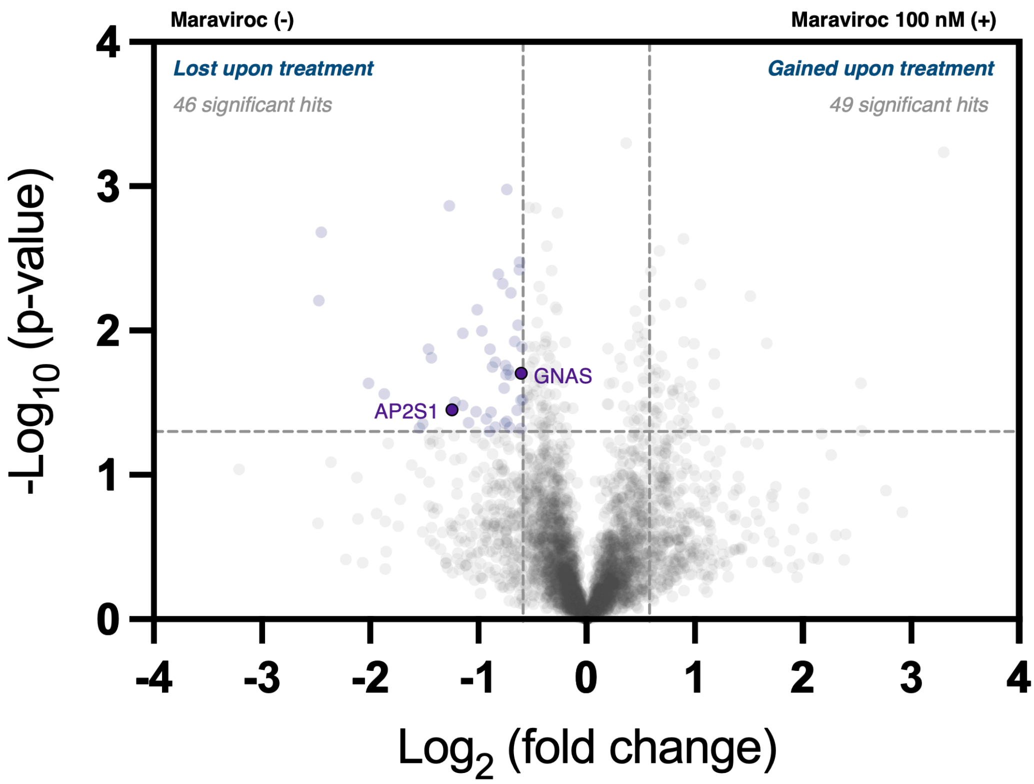

μMap-uAA was performed from the extracellular CCR5 F96* site to compare the receptor-proximal proteome in untreated and maraviroc-treated cells. In the untreated condition, F96* labeling enriched a set of CCR5-proximal membrane-associated proteins. Upon maraviroc treatment, several proteins showed reduced enrichment, including GNAS and AP2S1. Loss of GNAS and AP2S1 is consistent with antagonist-dependent remodeling of signaling- and endocytosis-associated proteins within the CCR5-proximal microenvironment.

**Figure S4B: L352* intracellular μMap-uAA identifies maraviroc-sensitive CCR5-proximal proteins**

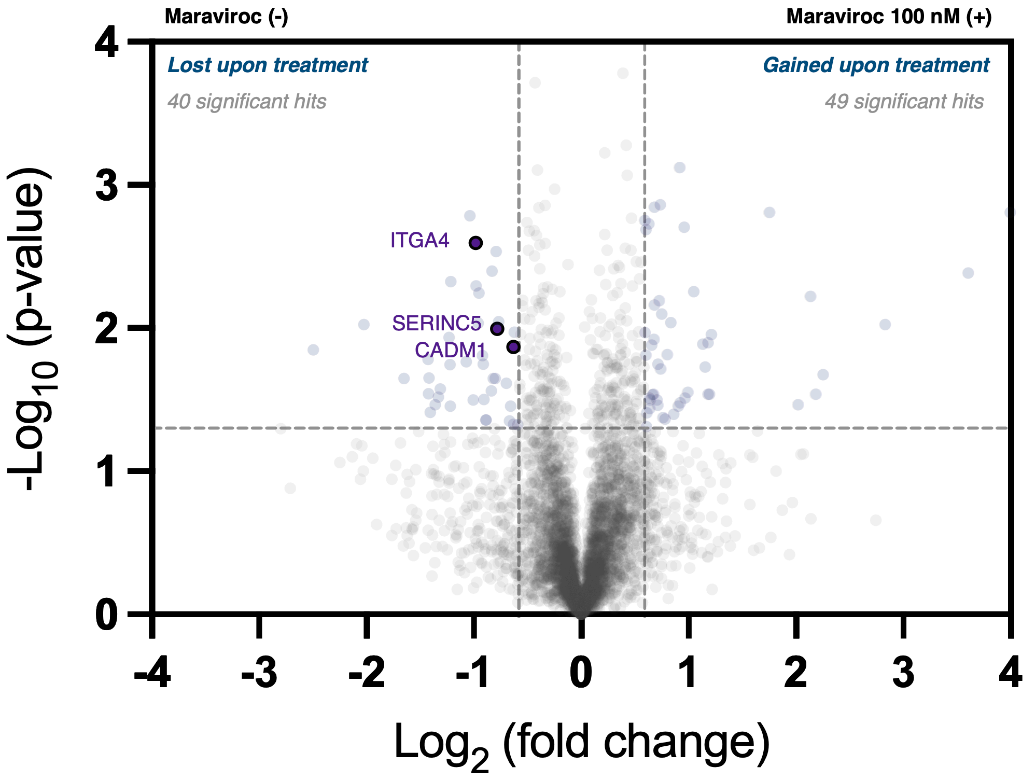

μMap-uAA was performed from the intracellular CCR5 L352* site to compare the receptor-proximal proteome in untreated and maraviroc-treated cells. In the untreated condition, L352* labeling enriched a set of intracellular and membrane-associated CCR5-proximal proteins. Upon maraviroc treatment, several proteins showed reduced enrichment, including CADM1, ITGA4, and SERINC5. These results further demonstrate that residue-defined μMap-uAA can capture antagonist-dependent remodeling of CCR5-proximal interactions from an intracellular receptor site.

**Figure S4C: CCL5/RANTES-induced remodeling of the CCR5-proximal interactome captured by μMap-uAA labeling**

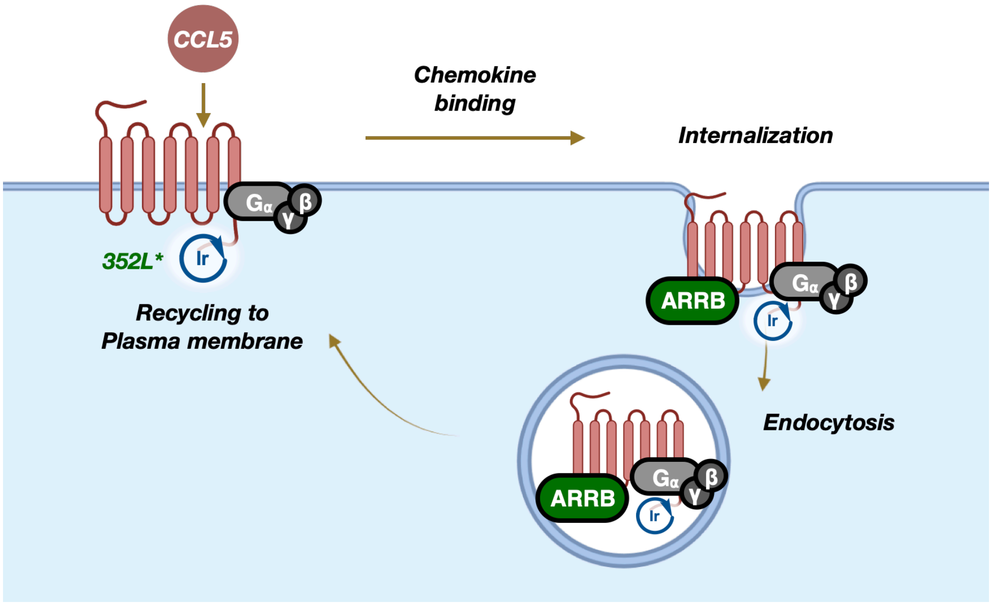

For CCL5/RANTES-treated samples, μMap–uAA labeling with CCR5 L352* was performed using separate plates for each stimulation condition. Following incubation with Ir–tetrazine (1 μM) for 2 h, cells were treated as described below. CCL5/RANTES (MedChemExpress, HY-P70450) was prepared as a 100 μM stock solution in $ddH_2O$ and used at a final concentration of 100 nM.

For the 0 min condition, no CCL5/RANTES was added. Cells were incubated with biotin–diazirine (**S1**, 500 μM) in phenol red-free complete DMEM for 30 min at 37 °C and immediately subjected to blue-light irradiation.

For the 5 min condition, cells were first incubated with biotin–diazirine (500 μM) for 25 min. CCL5/RANTES-containing medium was then added directly to achieve a final CCL5/RANTES concentration of 100 nM. After 5 min of stimulation, cells were immediately subjected to blue-light irradiation.

For the 120 min condition, cells were initially incubated with CCL5/RANTES (100 nM) for 90 min. The medium was then replaced with phenol red-free complete DMEM containing both CCL5/RANTES (100 nM) and biotin–diazirine (500 μM). Following an additional 30 min incubation, cells were immediately subjected to blue-light irradiation. Thus, the total CCL5/RANTES stimulation time before irradiation was 120 min.

**Figure S4D: CCR5-GNAO1 proximity after 2 h of CCL5 treatment identified by µMap-uAA**

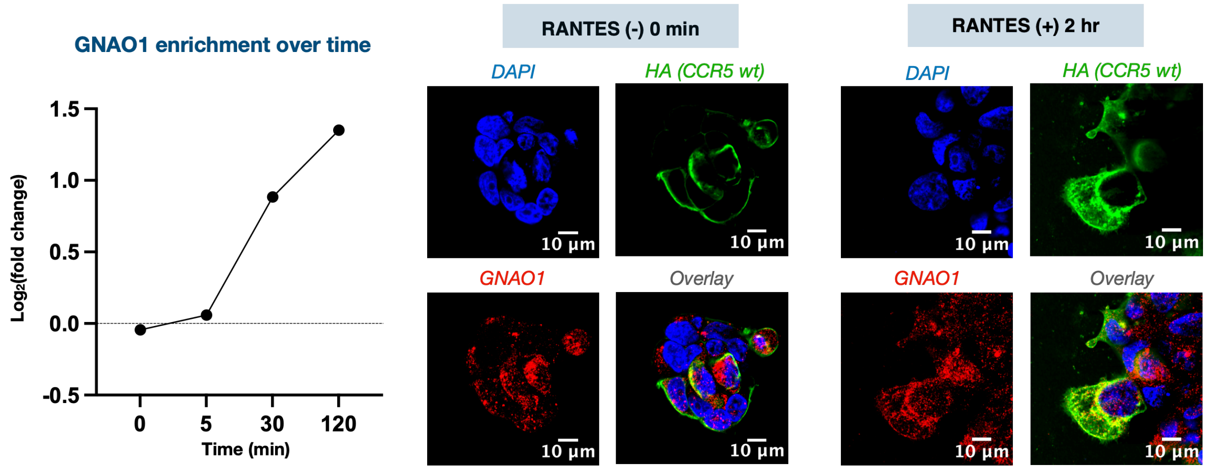

Left: GNAO1 enrichment in the CCR5-proximal proteome increased over the CCL5/RANTES stimulation time course, with the strongest enrichment observed after 2 h treatment. Right: Representative confocal microscopy images of cells expressing HA-tagged CCR5, stained for HA and GNAO1, in the absence or presence of CCL5/RANTES stimulation for 2 h. Increased overlap between CCR5 and GNAO1 was observed after prolonged ligand treatment. Although a functional relationship between CCR5 and GNAO1 remains to be established, these data further support the ability of µMap–uAA to detect ligand-induced remodeling of CCR5-proximal interactomes with temporal precision. Scale bars, 10 µm.

#### Figure S5: GLP-1R μMap-uAA labeling

##### Figure S5A: Western blot analysis of GLP-1R F61* labeling and enrichment

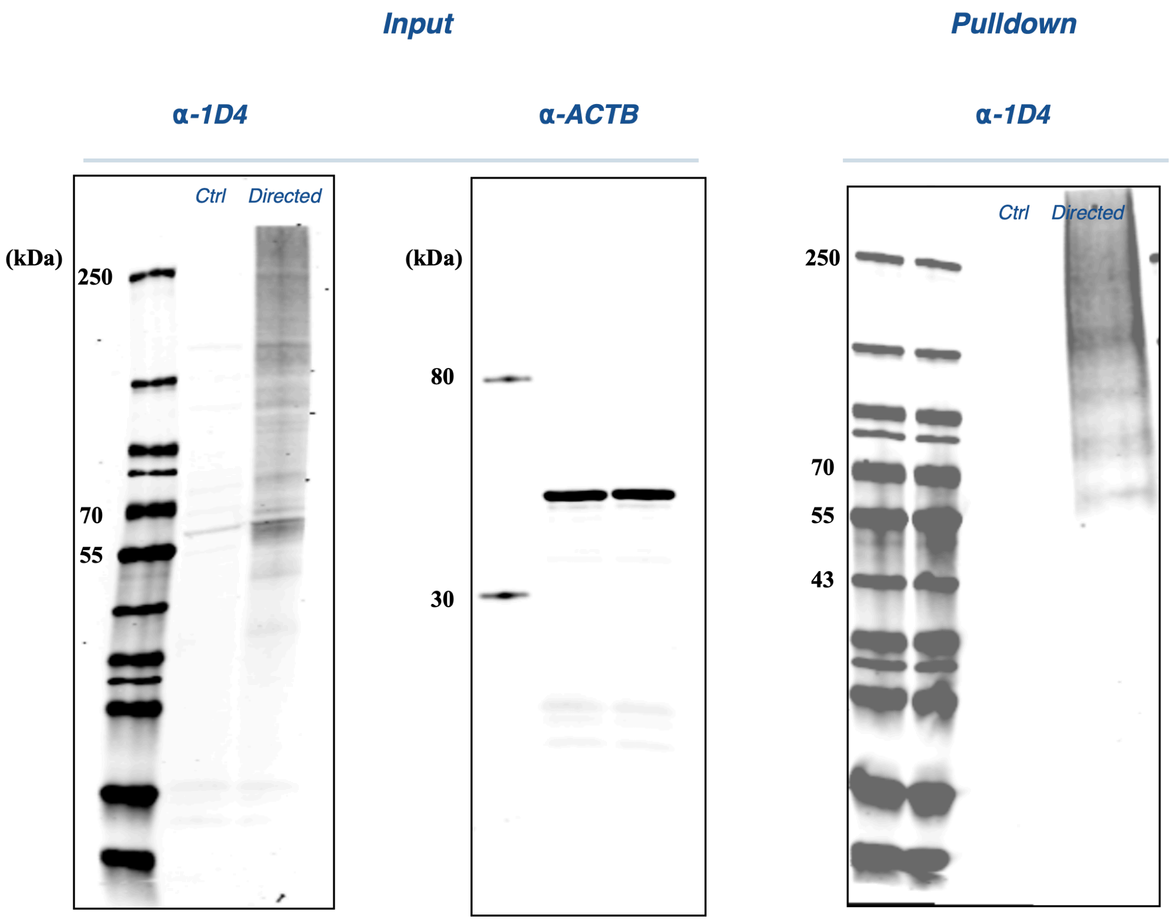

HEK293T cells were subjected to amber suppression in 6 cm dishes using either empty vector together with the amber suppression machinery (ctrl) or GLP-1R F61TAG together with the amber suppression machinery (directed) under the optimized conditions for GLP-1R F61TAG (GLP-1R F61TAG:tRNA/aaRS = 1:2, FuGENE HD, 30 h). Both groups were then subjected to iridium–tetrazine (1 μM) conjugation, live-cell photocatalytic labeling, and streptavidin bead enrichment. Input lysates and streptavidin elution fractions were analyzed by Western blot. Membranes were probed with anti-1D4 antibody (Santa Cruz Biotechnology, sc-57432, 1:200 dilution), followed by goat anti-mouse IRDye 680 CW secondary antibody (LI-COR), to detect the C-terminal epitope tag on GLP-1R. Detection of the 1D4 tag is indicative of full-length receptor expression rather than truncated protein arising from incomplete amber suppression. Input lysates were additionally probed for ACTB as a loading control; comparable ACTB signal intensities indicate similar protein input across ctrl and directed samples. 1D4 enrichment was observed in the streptavidin elution fraction, indicating successful self-labeling and enrichment of GLP-1R.

**Figure S5B: Confocal microscopy analysis of GLP-1R F61* μMap-uAA labeling**

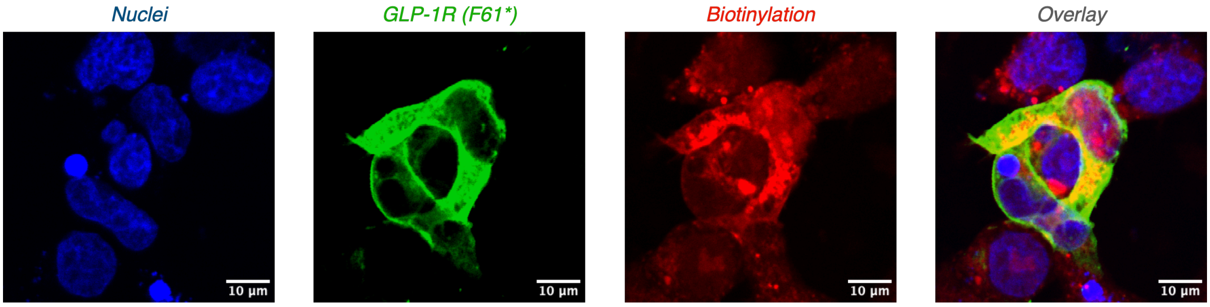

HEK293T cells were seeded into poly-D-lysine-coated glass-bottom 8-well chamber slides and allowed to recover overnight. Cells were transfected with plasmid encoding GLP-1R F61*TAG (0.2 μg) and the M21 tRNA/aaRS plasmid (0.4 μg) using FuGENE HD transfection reagent (1.8 μL) per well. μMap-uAA labeling and immunofluorescence microscopy were then performed according to the standard protocols described above.

#### Plasmids

##### Subcloning of β-actin from mCherry-C18-actin

The β-actin coding sequence was subcloned from mCherry-C18-actin (Addgene #54967) to generate a standalone actin expression construct lacking the upstream mCherry coding sequence. The cloning strategy used AgeI and BamHI restriction sites flanking the actin insert.

Insert PCR: The β-actin coding sequence was amplified from mCherry-C18-actin template plasmid DNA in a 50 μL PCR containing 25 μL 2× Phusion Master Mix, 2.5 μL each of forward and reverse primers (10 μM), 0.1 μL plasmid DNA, 1 μL DMSO, and 18.5 μL nuclease-free water.

The forward primer introduced an AgeI restriction site (ACCGGT), Kozak sequence (GCCACC), and ATG start codon upstream of the β-actin coding sequence. The reverse primer introduced a BamHI restriction site (GGATCC) downstream of the β-actin coding sequence. PCR cycling conditions were as follows: 98 °C for 30 s; 30 cycles of 98 °C for 10 s, 66 °C for 30 s, and 72 °C for 45 s; followed by 72 °C for 7 min and hold at 4 °C. The expected ~1.2 kb PCR product was gel-purified.

Backbone preparation: The mCherry-C18-actin plasmid was double-digested with AgeI and BamHI in a 50 µL reaction containing 5 µL 10× rCutSmart Buffer, 1 µL AgeI, 1 µL BamHI, plasmid DNA, and nuclease-free water. The digested backbone fragment was gel-purified.

Insert digestion: The purified β-actin PCR product was digested with AgeI and BamHI in a 50 µL reaction containing 5 µL 10× rCutSmart Buffer, 1 µL AgeI, 1 µL BamHI, PCR product, and nuclease-free water. The digested insert was purified by PCR cleanup.

Ligation and verification: The digested β-actin insert and AgeI/BamHI-digested backbone were ligated using T4 DNA Ligase at an approximately 3:1 insert:vector molar ratio. The ligation product was transformed into competent *E. coli*, and plasmid DNA was isolated from individual colonies by miniprep. The final construct was verified by Sanger sequencing.

**General procedure for plasmid construction and sequence verification**

Amber mutant constructs were generated by site-directed mutagenesis from the corresponding parent plasmids. Site-specific introduction of amber codons was performed using the Q5 Site-Directed Mutagenesis Kit (New England Biolabs) according to the manufacturer's protocol. Following mutagenesis, plasmids were transformed into competent *E. coli*, amplified, and purified using the QIAprep Spin Miniprep Kit (Qiagen, cat. no. 27104). All constructs were verified by whole-plasmid sequencing through Plasmidsaurus to confirm successful introduction of the desired TAG mutation and the absence of undesired mutations within the coding sequence. Parent plasmids, mutation sites, and primer sequences are listed in Supplementary Table S6.

Orthogonal tRNA/tRNA synthetase pair

The M21 plasmid encoding pNEU-chBCNRS2020/tRNAM15 was reconstructed in-house by molecular cloning according to the previously reported plasmid map.[1]

ACTB constructs

ACTB (K118TAG)-HA

tacggggtcattagttcatagcccatatatggagttccgcgttacataacttacggtaaatggcccgcctggctgaccgcccaacgacccccgcccattgacgtcaataatgacgtat
gttcccatagtaacgccaatagggactttccattgacgtcaatgggtggagtatttacggtaaactgcccacttggcagtacatcaagtgtatcatatgccaagtacgccccctattgac
gtcaatgacggtaaatggcccgcctggcattatgcccagtacatgaccttatgggactttcctacttggcagtacatctacgtattagtcatcgctattaccatggtgatgcggttttggca
gtacatcaatgggcgtggatagcggtttgactcacggggatttccaagtctccaccccattgacgtcaatgggagtttgttttggcaccaaaatcaacgggactttccaaaatgtcgtaa
caactccgccccattgacgcaaatgggcggtaggcgtgtacggtgggaggtctatataagcagagctggtttagtgaaccgtcagatccgctagcgctaccggtcgccaccACC
CATACGATGTTCCAGATTACGCTatggatgatgatatcgccgcgctcgtcgtcgacaacggctccggcatgtgcaaggccggcttcgcgggcgacgatgccccccg
ggccgtcttccccctccatcgtggggcgccccaggcaccagggcgtgatggtgggcatgggtcagaaggattcctatgtgggcgacgaggcccagagcaagagaggcatcctca
ccctgaagtaccccatcgagcacggcatcgtcaccaactgggacgacatggagaaaatctggcaccacaccttctacaatgagctgcgtgtggctcccgaggagcaccccgtgc
tgctgaccgaggcccccctgaaccccaaggccaaccgcgagTAGatgacccagatcatgtttgagaccttcaacaccccagccatgtacgttgctatccaggctgtgctatccctg
tacgcctctggccgtaccactggcatcgtgatggactccggtgacggggtcacccacactgtgcccatctacgaggggtatgccctcccccatgccatcctgcgtctggacctggctg
gccgggacctgactgactacctcatgaagatcctcaccgagcgcggctacagcttcaccaccacggccgagcgggaaatcgtgcgtgacattaaggagaagctgtgctacgtcg
ccctggacttcgagcaagagatggccacggctgcttccagctcctccctggagaagagctacgagctgcctgacggccaggtcatcaccattggcaatgagcggttccgctgccct

gaggcactcttccagccttccttcctgggcatggagtcctgtggcatccacgaaactaccttcaactccatcatgaagtgtgacgtggacatccgcaaagacctgtacgccaacaca
gtgctgtctggcggcaccaccatgtaccctggcattgccgacaggatgcagaaggagatcactgccctggcacccagcacaatgaagatcaagatcattgctcctcctgagcgca
agtactccgtgtggatcggcggctccatcctggcctcgctgtccaccttccagcagatgtggatcagcaagcaggagtatgacgagtccggcccctccatcgtccaccgcaaatgct
tcTACCCATACGATGTTCCAGATTACGCTtagggatccaccggatctagataactgatcataatcagccataccacatttgtagaggttttacttgctttaaaaaacctc
ccacacctccccctgaacctgaaacataaaatgaatgcaattgttgttgttaacttgtttattgcagcttataatggttacaaataaagcaatagcatcacaaatttcacaaataaagcat
ttttttcactgcattctagttgtggtttgtccaaactcatcaatgtatcttaaggcgtaaattgtaagcgttaatattttgttaaaattcgcgttaaatttttgttaaatcagctcattttttaaccaata
ggccgaaatcggcaaaatcccttataaatcaaaagaatagaccgagatagggttgagtgttgttccagtttggaacaagagtccactattaaagaacgtggactccaacgtcaaag
ggcgaaaaaccgtctatcagggcgatggcccactacgtgaaccatcaccctaatcaagttttttggggtcgaggtgccgtaaagcactaaatcggaaccctaaagggagcccccg
atttagagcttgacggggaaagccggcgaacgtggcgagaaaggaagggaagaaagcgaaaggagcgggcgctagggcgctggcaagtgtagcggtcacgctgcgcgta
accaccacacccgccgcgcttaatgcgccgctacagggcgcgtcaggtggcacttttcggggaaatgtgcgcggaacccctatttgtttatttttctaaatacattcaaatatgtatccg
ctcatgagacaataaccctgataaatgcttcaataatattgaaaaaggaagagtcctgaggcggaaagaaccagctgtggaatgtgtgtcagttagggtgtggaaagtccccagg
ctccccagcaggcagaagtatgcaaagcatgcatctcaattagtcagcaaccatagtcccgcccctaactccgcccatcccgcccctaactccgcccagttccgcccattctccgcc
ccatggctgactaattttttttatttatgcagaggccgaggccgcctcggcctctgagctattccagaagtagtgaggaggcttttttggaggcctaggcttttgcaaagatcgatcaaga
gacaggatgaggatcgtttcgcatgattgaacaagatggattgcacgcaggttctccggccgcttgggtggagaggctattcggctatgactgggcacaacagacaatcggctgctc
tgatgccgccgtgttccggctgtcagcgcaggggcgcccggttctttttgtcaagaccgacctgtccggtgccctgaatgaactgcaagacgaggcagcgcggctatcgtggctggc
cacgacgggcgttccttgcgcagctgtgctcgacgttgtcactgaagcgggaagggactggctgctattgggcgaagtgccggggcaggatctcctgtcatctcaccttgctcctgcc
gagaaagtatccatcatggctgatgcaatgcggcggctgcatacgcttgatccggctacctgcccattcgaccaccaagcgaaacatcgcatcgagcgagcacgtactcggatgg
aagccggtcttgtcgatcaggatgatctggacgaagagcatcaggggctcgcgccagccgaactgttcgccaggctcaaggcgagcatgcccgacggcgaggatctcgtcgtga
cccatggcgatgcctgcttgccgaatatcatggtggaaaatggccgcttttctggattcatcgactgtggccggctgggtgtggcggaccgctatcaggacatagcgttggctacccgt
gatattgctgaagagcttggcggcgaatgggctgaccgcttcctcgtgctttacggtatcgccgctcccgattcgcagcgcatcgccttctatcgccttcttgacgagttcttctgagcggg
actctggggttcgaaatgaccgaccaagcgacgcccaacctgccatcacgagatttcgattccaccgccgccttctatgaaaggttgggcttcggaatcgttttccgggacgccggct
ggatgatcctccagcgcggggatctcatgctggagttcttcgcccaccctagggggaggctaactgaaacacggaaggagacaataccggaaggaacccgcgctatgacggca
ataaaaagacagaataaaacgcacggtgttgggtcgtttgttcataaacgcggggttcggtcccagggctggcactctgtcgataccccaccgagaccccattggggccaatacgc
ccgcgtttcttccttttccccaccccaccccccaagttcgggtgaaggcccagggctcgcagccaacgtcggggcggcaggccctgccatagcctcaggttactcatatatactttag
attgatttaaaacttcatttttaatttaaaaggatctaggtgaagatcctttttgataatctcatgaccaaaatcccttaacgtgagttttcgttccactgagcgtcagaccccgtagaaaaga
tcaaaggatcttcttgagatcctttttttctgcgcgtaatctgctgcttgcaaacaaaaaaaccaccgctaccagcggtggtttgtttgccggatcaagagctaccaactctttttccgaagg
taactggcttcagcagagcgcagataccaaatactgttcttctagtgtagccgtagttaggccaccacttcaagaactctgtagcaccgcctacatacctcgctctgctaatcctgttacc
agtggctgctgccagtggcgataagtcgtgtcttaccgggttggactcaagacgatagttaccggataaggcgcagcggtcgggctgaacggggggttcgtgcacacagcccagc
ttggagcgaacgacctacaccgaactgagatacctacagcgtgagctatgagaaagcgccacgcttcccgaagggagaaaggcggacaggtatccggtaagcggcagggtc
ggaacaggagagcgcacgagggagcttccagggggaaacgcctggtatctttatagtcctgtcgggtttcgccacctctgacttgagcgtcgatttttgtgatgctcgtcaggggggc
ggagcctatggaaaaacgccagcaacgcggcctttttacggttcctggccttttgctggccttttgctcacatgttctttcctgcgttatcccctgattctgtggataaccgtattaccgccatg
cattagttattaatagtaatcaat

#### NONO constructs

Human NONO cDNA bearing an N-terminal HA tag was obtained from Addgene (plasmid #127655; pcDNA3.1-HA-NONO).

##### HA-NONO (A258TAG)

gacggatcgggagatctcccgatcccctatggtgcactctcagtacaatctgctctgatgccgcatagttaagccagtatctgctccctgcttgtgtgttggaggtcgctgagtagtgcgc
gagcaaaatttaagctacaacaaggcaaggcttgaccgacaattgcatgaagaatctgcttagggttaggcgttttgcgctgcttcgcgatgtacgggccagatatacgcgttgacat
tgattattgactagttattaatagtaatcaattacggggtcattagttcatagcccatatatggagttccgcgttacataacttacggtaaatggcccgcctggctgaccgcccaacgacc
cccgcccattgacgtcaataatgacgtatgttcccatagtaacgccaatagggactttccattgacgtcaatgggtggagtatttacggtaaactgcccacttggcagtacatcaagtgt
atcatatgccaagtacgccccctattgacgtcaatgacggtaaatggcccgcctggcattatgcccagtacatgaccttatgggactttcctacttggcagtacatctacgtattagtcat
cgctattaccatggtgatgcggttttggcagtacatcaatgggcgtggatagcggtttgactcacggggatttccaagtctccaccccattgacgtcaatgggagtttgttttggcaccaa
aatcaacgggactttccaaaatgtcgtaacaactccgccccattgacgcaaatgggcggtaggcgtgtacggtgggaggtctatataagcagagctctctggctaactagagaacc
cactgcttactggcttatcgaaattaatacgactcactatagggagacccaagcttagccaccaatatgtacccatacgatgttccagattacgctggagcaagggatccaatgcag
agtaataaaacttttaacttggagaagcaaaaccatactccaagaaagcatcatcaacatcaccaccagcagcagcaccaccagcagcaacagcagcagccgccaccaccg
ccaatacctgcaaatgggcaacaggccagcagccaaaatgaaggcttgactattgacctgaagaattttagaaaaccaggagagaagaccttcacccaacgaagccgtcttttttg
tgggaaatcttcctcccgacatcactgaggaagaaatgaggaaactatttgagaaatatggaaaggcaggcgaagtcttcattcataaggataaaggatttggctttatccgcttgga
aacccgaaccctagcggagattgccaaagtggagctggacaatatgccactccgtggaaagcagctgcgtgtgcgctttgcctgccatagtgcatcccttacagttcgaaaccttcc
tcagtatgtgtccaacgaactgctggaagaagccttttctgtgtttggccaggtagagagggctgtagtcattgtggatgatcgaggaaggccctcaggaaaaggcattgttgagttctc
agggaagccagctgctcggaaagctctggacagatgcagtgaaggctccttcctgctaaccacatttcctcgtcctgtgactgtggagcccatggaccagttagatgatgaagagg
gacttccagagaagctggttataaaaaaccagcaatttcacaaggaacgagagcagccacccagattttagcagcctggctcctttgagtatgaatatgccatgcgctggaaggca
ctcattgagatggagaagcagcagcaggaccaagtggaccgcaacatcaaggaggctcgtgagaagctggagatggagatggaagctgcacgccatgagcaccaggtcatg
ctaatgagacaggatttgatgaggcgccaagaagaacttcggaggatggaagagctgcacaaccaagaggtgcaaaaacgaaagcaactggagctcaggcaggaggaag
agcgcaggcgccgtgaagaagagatgcggcggcagcaagaagaaatgatgcggcgacagcaggaaggattcaagggaaccttccctgatgcgagagagcaggagattcg
gatgggtcagatggctatgggaggtgctatgggcataaacaacagaggtgccatgccccctgctcctgtgccagctggtaccccagctcctccaggacctgccactatgatgccgg
atggaactttgggattgaccccaccaacaactgaacgctttggtcaggctgctacaatggaaggaattggggcaattggtggaactcctcctgcattcaaccgtgcagctcctggag
ctgaatttgtcccaaacaaacgtcgccgatactaactcgagcatgcatctagagggccctattctatagtgtcacctaaatgctagagctcgctgatcagcctcgactgtgccttctagtt
gccagccatctgttgtttgcccctcccccgtgccttccttgaccctggaaggtgccactcccactgtcctttcctaataaaatgaggaaattgcatcgcattgtctgagtaggtgtcattctat
tctggggggtggggtggggcaggacagcaagggggaggattgggaagacaatagcaggcatgctggggatgcggtgggctctatggcttctgaggcggaaagaaccagctgg
ggctctagggggtatccccacgcgccctgtagcggcgcattaagcgcggcgggtgtggtggttacgcgcagcgtgaccgctacacttgccagcgccctagcgcccgctcctttcgct
ttcttcccttcctttctcgccacgttcgccggctttccccgtcaagctctaaatcgggggctccctttagggttccgatttagtgctttacggcacctcgaccccaaaaaacttgattagggtg
atggttcacgtagtgggccatcgccctgatagacggtttttcgccctttgacgttggagtccacgttctttaatagtggactcttgttccaaactggaacaacactcaaccctatctcggtct
attcttttgatttataagggattttgccgatttcggcctattggttaaaaaatgagctgatttaacaaaaatttaacgcgaattaattctgtggaatgtgtgtcagttagggtgtggaaagtccc
caggctccccagcaggcagaagtatgcaaagcatgcatctcaattagtcagcaaccaggtgtggaaagtccccaggctccccagcaggcagaagtatgcaaagcatgcatctc
aattagtcagcaaccatagtcccgcccctaactccgcccatcccgcccctaactccgcccagttccgcccattctccgccccatggctgactaattttttttatttatgcagaggccgagg
ccgcctctgcctctgagctattccagaagtagtgaggaggcttttttggaggcctaggcttttgcaaaaagctcccgggagcttgtatatccattttcggatctgatcaagagacaggatg

aggatcgtttcgcatgattgaacaagatggattgcacgcaggttctccggccgcttgggtggagaggctattcggctatgactgggcacaacagacaatcggctgctctgatgccgc
cgtgttccggctgtcagcgcaggggcgcccggttctttttgtcaagaccgacctgtccggtgccctgaatgaactgcaggacgaggcagcgcggctatcgtggctggccacgacgg
gcgttccttgcgcagctgtgctcgacgttgtcactgaagcgggaagggactggctgctattgggcgaagtgccggggcaggatctcctgtcatctcaccttgctcctgccgagaaagt
atccatcatggctgatgcaatgcggcggctgcatacgcttgatccggctacctgcccattcgaccaccaagcgaaacatcgcatcgagcgagcacgtactcggatggaagccggt
cttgtcgatcaggatgatctggacgaagagcatcaggggctcgcgccagccgaactgttcgccaggctcaaggcgcgcatgcccgacggcgaggatctcgtcgtgacccatggc
gatgcctgcttgccgaatatcatggtggaaaatggccgcttttctggattcatcgactgtggccggctgggtgtggcggaccgctatcaggacatagcgttggctacccgtgatattgct
gaagagcttggcggcgaatgggctgaccgcttcctcgtgctttacggtatcgccgctcccgattcgcagcgcatcgccttctatcgccttcttgacgagttcttctgagcgggactctggg
gttcgaaatgaccgaccaagcgacgcccaacctgccatcacgagatttcgattccaccgccgccttctatgaaaggttgggcttcggaatcgttttccgggacgccggctggatgatc
ctccagcgcggggatctcatgctggagttcttcgcccaccccaacttgtttattgcagcttataatggttacaaataaagcaatagcatcacaaatttcacaaataaagcatttttttcact
gcattctagttgtggtttgtccaaactcatcaatgtatcttatcatgtctgtataccgtcgacctctagctagagcttggcgtaatcatggtcatagctgtttcctgtgtgaaattgttatccgctc
acaattccacacaacatacgagccggaagcataaagtgtaaagcctggggtgcctaatgagtgagctaactcacattaattgcgttgcgctcactgcccgctttccagtcgggaaac
ctgtcgtgccagctgcattaatgaatcggccaacgcgcggggagaggcggtttgcgtattgggcgctcttccgcttcctcgctcactgactcgctgcgctcggtcgttcggctgcggcg
agcggtatcagctcactcaaaggcggtaatacggttatccacagaatcaggggataacgcaggaaagaacatgtgagcaaaaggccagcaaaaggccaggaaccgtaaaa
aggccgcgttgctggcgtttttccataggctccgcccccctgacgagcatcacaaaaatcgacgctcaagtcagaggtggcgaaacccgacaggactataaagataccaggcgtt
tccccctggaagctccctcgtgcgctctcctgttccgaccctgccgcttaccggatacctgtccgcctttctcccttcgggaagcgtggcgctttctcatagctcacgctgtaggtatctcag
ttcggtgtaggtcgttcgctccaagctgggctgtgtgcacgaaccccccgttcagcccgaccgctgcgccttatccggtaactatcgtcttgagtccaacccggtaagacacgacttat
cgccactggcagcagccactggtaacaggattagcagagcgaggtatgtaggcggtgctacagagttcttgaagtggtggcctaactacggctacactagaagaacagtatttggt
atctgcgctctgctgaagccagttaccttcggaaaaagagttggtagctcttgatccggcaaacaaaccaccgctggtagcggtggtttttttgtttgcaagcagcagattacgcgcag
aaaaaaaggatctcaagaagatcctttgatcttttctacggggtctgacgctcagtggaacgaaaactcacgttaagggattttggtcatgagattatcaaaaaggatcttcacctagat
ccttttaaattaaaaatgaagttttaaatcaatctaaagtatatatgagtaaacttggtctgacagttaccaatgcttaatcagtgaggcacctatctcagcgatctgtctatttcgttcatcca
tagttgcctgactccccgtcgtgtagataactacgatacgggagggcttaccatctggccccagtgctgcaatgataccgcgagacccacgctcaccggctccagatttatcagcaat
aaaccagccagccggaagggccgagcgcagaagtggtcctgcaactttatccgcctccatccagtctattaattgttgccgggaagctagagtaagtagttcgccagttaatagtttg
cgcaacgttgttgccattgctacaggcatcgtggtgtcacgctcgtcgtttggtatggcttcattcagctccggttcccaacgatcaaggcgagttacatgatcccccatgttgtgcaaaa
aagcggttagctccttcggtcctccgatcgttgtcagaagtaagttggccgcagtgttatcactcatggttatggcagcactgcataattctcttactgtcatgccatccgtaagatgcttttct
gtgactggtgagtactcaaccaagtcattctgagaatagtgtatgcggcgaccgagttgctcttgcccggcgtcaatacgggataataccgcgccacatagcagaactttaaaagtg
ctcatcattggaaaacgttcttcggggcgaaaactctcaaggatcttaccgctgttgagatccagttcgatgtaacccactcgtgcacccaactgatcttcagcatcttttactttcaccag
cgtttctgggtgagcaaaaacaggaaggcaaaatgccgcaaaaaagggaataagggcgacacggaaatgttgaatactcatactcttcctttttcaatattattgaagcatttatcag
ggttattgtctcatgagcggatacatatttgaatgtatttagaaaaataaacaaataggggttccgcgcacatttccccgaaaagtgccacctgacgtc

#### CCR5 constructs

Human CCR5 cDNA was obtained from Addgene (plasmid #98943; pcDNA3-CCR5).

##### CCR5 (F96TAG) HA

Atggattatcaagtgtcaagtccaatctatgacatcaattatacatcggagccctgccaaaaaatcaatgtgaagcaaatcgcagcccgcctcctgcctccgctctactcactggt
gttcatctttggttttgtgggcaacatgctggtcatcctcatcctgataaactgcaaaaggctgaagagcatgactgacatctacctgctcaacctggccatctctgacctgtttttccttctta
ctgtccccttctgggctcactatgctgccgcccagtgggactagggaaatacaatgtgtcaactcttgacagggctctattttataggcttcttctctggaatcttcttcatcatcctcctgaca
atcgataggtacctggctgtcgtccatgctgtgtttgctttaaaagccaggacggtcacctttggggtggtgacaagtgtgatcacttgggtggtggctgtgtttgcgtctctcccaggaatc
atctttaccagatctcaaaaagaaggtcttcattacacctgcagctctcattttccatacagtcagtatcaattctggaagaatttccagacattaaagatagtcatcttggggctggtcctg
ccgctgcttgtcatggtcatctgctactcgggaatcctaaaaactctgcttcggtgtcgaaatgagaagaagaggcacagggctgtgaggcttatcttcaccatcatgattgtttattttctc
ttctgggctccctacaacattgtccttctcctgaacaccttccaggaattctttggcctgaataattgcagtagctctaacaggttggaccaagctatgcaggtgacagagactcttgggat
gacgcactgctgcatcaaccccatcatctatgcctttgtcggggagaagttcagaaactacctcttagtcttcttccaaaagcacattgccaaacgcttctgcaaatgctgttctattttcca
gcaagaggctcccgagcgagcaagctcagtttacacccgatccactggggagcaggaaatatctgtgggcttgTACCCATACGATGTTCCAGATTACGCTtgat
aactcgagtctagagggcccgtttaaacccgctgatcagcctcgactgtgccttctagttgccagccatctgttgtttgcccctcccccgtgccttccttgaccctggaaggtgccactcc
cactgtcctttcctaataaaatgaggaaattgcatcgcattgtctgagtaggtgtcattctattctggggggtggggtggggcaggacagcaagggggaggattgggaagacaatag
caggcatgctggggatgcggtgggctctatggcttctgaggcggaaagaaccagctggggctctagggggtatccccacgcgccctgtagcggcgcattaagcgcggcgggtgt
ggtggttacgcgcagcgtgaccgctacacttgccagcgccctagcgcccgctcctttcgctttcttcccttcctttctcgccacgttcgccggctttccccgtcaagctctaaatcgggggc
tccctttagggttccgatttagtgctttacggcacctcgaccccaaaaaacttgattagggtgatggttcacgtagtgggccatcgccctgatagacggtttttcgccctttgacgttggagt
ccacgttctttaatagtggactcttgttccaaactggaacaacactcaaccctatctcggtctattcttttgatttataagggattttgccgatttcggcctattggttaaaaaatgagctgattta
acaaaaatttaacgcgaattaattctgtggaatgtgtgtcagttagggtgtggaaagtccccaggctccccagcaggcagaagtatgcaaagcatgcatctcaattagtcagcaacc
aggtgtggaaagtccccaggctccccagcaggcagaagtatgcaaagcatgcatctcaattagtcagcaaccatagtcccgcccctaactccgcccatcccgcccctaactccgc
ccagttccgcccattctccgccccatggctgactaattttttttatttatgcagaggccgaggccgcctctgcctctgagctattccagaagtagtgaggaggcttttttggaggcctaggct
tttgcaaaaagctcccgggagcttgtatatccattttcggatctgatcaagagacaggatgaggatcgtttcgcatgattgaacaagatggattgcacgcaggttctccggccgcttgg
gtggagaggctattcggctatgactgggcacaacagacaatcggctgctctgatgccgccgtgttccggctgtcagcgcaggggcgcccggttctttttgtcaagaccgacctgtccg
gtgccctgaatgaactgcaggacgaggcagcgcggctatcgtggctggccacgacgggcgttccttgcgcagctgtgctcgacgttgtcactgaagcgggaagggactggctgct
attgggcgaagtgccggggcaggatctcctgtcatctcaccttgctcctgccgagaaagtatccatcatggctgatgcaatgcggcggctgcatacgcttgatccggctacctgcccat
tcgaccaccaagcgaaacatcgcatcgagcgagcacgtactcggatggaagccggtcttgtcgatcaggatgatctggacgaagagcatcaggggctcgcgccagccgaactg
ttcgccaggctcaaggcgcgcatgcccgacggcgaggatctcgtcgtgacccatggcgatgcctgcttgccgaatatcatggtggaaaatggccgcttttctggattcatcgactgtg
gccggctgggtgtggcggaccgctatcaggacatagcgttggctacccgtgatattgctgaagagcttggcggcgaatgggctgaccgcttcctcgtgctttacggtatcgccgctcc
cgattcgcagcgcatcgccttctatcgccttcttgacgagttcttctgagcgggactctggggttcgaaatgaccgaccaagcgacgcccaacctgccatcacgagatttcgattccac
cgccgccttctatgaaaggttgggcttcggaatcgttttccgggacgccggctggatgatcctccagcgcggggatctcatgctggagttcttcgcccaccccaacttgtttattgcagctt
ataatggttacaaataaagcaatagcatcacaaatttcacaaataaagcatttttttcactgcattctagttgtggtttgtccaaactcatcaatgtatcttatcatgtctgtataccgtcgacc
tctagctagagcttggcgtaatcatggtcatagctgtttcctgtgtgaaattgttatccgctcacaattccacacaacatacgagccggaagcataaagtgtaaagcctggggtgcctaa
tgagtgagctaactcacattaattgcgttgcgctcactgcccgctttccagtcgggaaacctgtcgtgccagctgcattaatgaatcggccaacgcgcggggagaggcggtttgcgtat
tgggcgctcttccgcttcctcgctcactgactcgctgcgctcggtcgttcggctgcggcgagcggtatcagctcactcaaaggcggtaatacggttatccacagaatcaggggataac
gcaggaaagaacatgtgagcaaaaggccagcaaaaggccaggaaccgtaaaaaggccgcgttgctggcgtttttccataggctccgcccccctgacgagcatcacaaaaatc
gacgctcaagtcagaggtggcgaaacccgacaggactataaagataccaggcgtttccccctggaagctccctcgtgcgctctcctgttccgaccctgccgcttaccggatacctgt
ccgcctttctcccttcgggaagcgtggcgctttctcatagctcacgctgtaggtatctcagttcggtgtaggtcgttcgctccaagctgggctgtgtgcacgaaccccccgttcagcccga
ccgctgcgccttatccggtaactatcgtcttgagtccaacccggtaagacacgacttatcgccactggcagcagccactggtaacaggattagcagagcgaggtatgtaggcggtg
ctacagagttcttgaagtggtggcctaactacggctacactagaagaacagtatttggtatctgcgctctgctgaagccagttaccttcggaaaaagagttggtagctcttgatccggca
aacaaaccaccgctggtagcggtggtttttttgtttgcaagcagcagattacgcgcagaaaaaaaggatctcaagaagatcctttgatcttttctacggggtctgacgctcagtggaac

gaaaactcacgttaagggattttggtcatgagattatcaaaaaggatcttcacctagatccttttaaattaaaaatgaagttttaaatcaatctaaagtatatatgagtaaacttggtctgac
agttaccaatgcttaatcagtgaggcacctatctcagcgatctgtctatttcgttcatccatagttgcctgactccccgtcgtgtagataactacgatacgggagggcttaccatctggccc
cagtgctgcaatgataccgcgagacccacgctcaccggctccagatttatcagcaataaaccagccagccggaagggccgagcgcagaagtggtcctgcaactttatccgcctc
catccagtctattaattgttgccgggaagctagagtaagtagttcgccagttaatagtttgcgcaacgttgttgccattgctacaggcatcgtggtgtcacgctcgtcgtttggtatggcttca
ttcagctccggttcccaacgatcaaggcgagttacatgatcccccatgttgtgcaaaaaagcggttagctccttcggtcctccgatcgttgtcagaagtaagttggccgcagtgttatca
ctcatggttatggcagcactgcataattctcttactgtcatgccatccgtaagatgcttttctgtgactggtgagtactcaaccaagtcattctgagaatagtgtatgcggcgaccgagttgc
tcttgcccggcgtcaatacgggataataccgcgccacatagcagaactttaaaagtgctcatcattggaaaacgttcttcggggcgaaaactctcaaggatcttaccgctgttgagat
ccagttcgatgtaacccactcgtgcacccaactgatcttcagcatcttttactttcaccagcgtttctgggtgagcaaaaacaggaaggcaaaatgccgcaaaaaagggaataagg
gcgacacggaaatgttgaatactcatactcttcctttttcaatattattgaagcatttatcagggttattgtctcatgagcggatacatatttgaatgtatttagaaaaataaacaaataggggttccgcgcacatttccccgaaaagtgccacctgacgtcgacggatcgggagatctcccgatcccctatggtgcactctcagtacaatctgctctgatgccgcatagttaagccagtatc
tgctccctgcttgtgtgttggaggtcgctgagtagtgcgcgagcaaaatttaagctacaacaaggcaaggcttgaccgacaattgcatgaagaatctgcttagggttaggcgttttgcg
ctgcttcgcgatgtacgggccagatatacgcgttgacattgattattgactagttattaatagtaatcaattacggggtcattagttcatagcccatatatggagttccgcgttacataactta
cggtaaatggcccgcctggctgaccgcccaacgacccccgcccattgacgtcaataatgacgtatgttcccatagtaacgccaatagggactttccattgacgtcaatgggtggagt
atttacggtaaactgcccacttggcagtacatcaagtgtatcatatgccaagtacgccccctattgacgtcaatgacggtaaatggcccgcctggcattatgcccagtacatgaccttat
gggactttcctacttggcagtacatctacgtattagtcatcgctattaccatggtgatgcggttttggcagtacatcaatgggcgtggatagcggtttgactcacggggatttccaagtctcc
accccattgacgtcaatgggagtttgttttggcaccaaaatcaacgggactttccaaaatgtcgtaacaactccgccccattgacgcaaatgggcggtaggcgtgtacggtgggagg
tctatataagcagagctctctggctaactagagaacccactgcttactggcttatcgaaattaatacgactcactatagggagacccaagctggctagcGccacc

#### CCR5 (L352TAG) HA

Atggattatcaagtgtcaagtccaatctatgacatcaattattatacatcggagccctgccaaaaaatcaatgtgaagcaaatcgcagcccgcctcctgcctccgctctactcactggt
gttcatctttggttttgtgggcaacatgctggtcatcctcatcctgataaactgcaaaaggctgaagagcatgactgacatctacctgctcaacctggccatctctgacctgtttttccttcttactgtccccttctgggctcactatgctgccgcccagtgggactttggaaatacaatgtgtcaactcttgacagggctctattttataggcttcttctctggaatcttcttcatcatcctcctgacaa
tcgataggtacctggctgtcgtccatgctgtgtttgctttaaaagccaggacggtcacctttggggtggtgacaagtgtgatcacttgggtggtggctgtgtttgcgtctctcccaggaatca
tctttaccagatctcaaaaagaaggtcttcattacacctgcagctctcattttccatacagtcagtatcaattctggaagaatttccagacattaaagatagtcatcttggggctggtcctgc
cgctgcttgtcatggtcatctgctactcgggaatcctaaaaactctgcttcggtgtcgaaatgagaagaagaggcacagggctgtgaggcttatcttcaccatcatgattgtttattttctctt
ctgggctccctacaacattgtccttctcctgaacaccttccaggaattctttggcctgaataattgcagtagctctaacaggttggaccaagctatgcaggtgacagagactcttgggatg
acgcactgctgcatcaaccccatcatctatgcctttgtcggggagaagttcagaaactacctcttagtcttcttccaaaagcacattgccaaacgcttctgcaaatgctgttctattttccag
caagaggctcccgagcgagcaagctcagtttacacccgatccactggggagcaggaaatatctgtgggctagTACCCATACGATGTTCCAGATTACGCTtgata
actcgagtctagagggcccgtttaaacccgctgatcagcctcgactgtgccttctagttgccagccatctgttgtttgcccctcccccgtgccttccttgaccctggaaggtgccactccc
actgtcctttcctaataaaatgaggaaattgcatcgcattgtctgagtaggtgtcattctattctggggggtggggtggggcaggacagcaagggggaggattgggaagacaatagc
aggcatgctggggatgcggtgggctctatggcttctgaggcggaaagaaccagctggggctctagggggtatccccacgcgccctgtagcggcgcattaagcgcggcgggtgtg
gtggttacgcgcagcgtgaccgctacacttgccagcgccctagcgcccgctcctttcgctttcttcccttcctttctcgccacgttcgccggctttccccgtcaagctctaaatcgggggct
ccctttagggttccgatttagtgctttacggcacctcgaccccaaaaaacttgattagggtgatggttcacgtagtgggccatcgccctgatagacggtttttcgccctttgacgttggagtc
cacgttctttaatagtggactcttgttccaaactggaacaacactcaaccctatctcggtctattcttttgatttataagggattttgccgatttcggcctattggttaaaaaatgagctgattta
acaaaaatttaacgcgaattaattctgtggaatgtgtgtcagttagggtgtggaaagtccccaggctccccagcaggcagaagtatgcaaagcatgcatctcaattagtcagcaacc
aggtgtggaaagtccccaggctccccagcaggcagaagtatgcaaagcatgcatctcaattagtcagcaaccatagtcccgcccctaactccgcccatcccgcccctaactccgc
ccagttccgcccattctccgccccatggctgactaattttttttatttatgcagaggccgaggccgcctctgcctctgagctattccagaagtagtgaggaggcttttttggaggcctaggct
tttgcaaaaagctcccgggagcttgtatatccattttcggatctgatcaagagacaggatgaggatcgtttcgcatgattgaacaagatggattgcacgcaggttctccggccgcttgg
gtggagaggctattcggctatgactgggcacaacagacaatcggctgctctgatgccgccgtgttccggctgtcagcgcaggggcgcccggttctttttgtcaagaccgacctgtccg
gtgccctgaatgaactgcaggacgaggcagcgcggctatcgtggctggccacgacgggcgttccttgcgcagctgtgctcgacgttgtcactgaagcgggaagggactggctgct
attgggcgaagtgccggggcaggatctcctgtcatctcaccttgctcctgccgagaaagtatccatcatggctgatgcaatgcggcggctgcatacgcttgatccggctacctgcccat
tcgaccaccaagcgaaacatcgcatcgagcgagcacgtactcggatggaagccggtcttgtcgatcaggatgatctggacgaagagcatcaggggctcgcgccagccgaactg
ttcgccaggctcaaggcgcgcatgcccgacggcgaggatctcgtcgtgacccatggcgatgcctgcttgccgaatatcatggtggaaaatggccgcttttctggattcatcgactgtg
gccggctgggtgtggcggaccgctatcaggacatagcgttggctacccgtgatattgctgaagagcttggcggcgaatgggctgaccgcttcctcgtgctttacggtatcgccgctcc
cgattcgcagcgcatcgccttctatcgccttcttgacgagttcttctgagcgggactctggggttcgaaatgaccgaccaagcgacgcccaacctgccatcacgagatttcgattccac
cgccgccttctatgaaaggttgggcttcggaatcgttttccgggacgccggctggatgatcctccagcgcggggatctcatgctggagttcttcgcccaccccaacttgtttattgcagctt
ataatggttacaaataaagcaatagcatcacaaatttcacaaataaagcatttttttcactgcattctagttgtggtttgtccaaactcatcaatgtatcttatcatgtctgtataccgtcgacc
tctagctagagcttggcgtaatcatggtcatagctgtttcctgtgtgaaattgttatccgctcacaattccacacaacatacgagccggaagcataaagtgtaaagcctggggtgcctaa
tgagtgagctaactcacattaattgcgttgcgctcactgcccgctttccagtcgggaaacctgtcgtgccagctgcattaatgaatcggccaacgcgcggggagaggcggtttgcgtat
tgggcgctcttccgcttcctcgctcactgactcgctgcgctcggtcgttcggctgcggcgagcggtatcagctcactcaaaggcggtaatacggttatccacagaatcaggggataac
gcaggaaagaacatgtgagcaaaaggccagcaaaaggccaggaaccgtaaaaaggccgcgttgctggcgtttttccataggctccgcccccctgacgagcatcacaaaaatc
gacgctcaagtcagaggtggcgaaacccgacaggactataaagataccaggcgtttccccctggaagctccctcgtgcgctctcctgttccgaccctgccgcttaccggatacctgt
ccgcctttctcccttcgggaagcgtggcgctttctcatagctcacgctgtaggtatctcagttcggtgtaggtcgttcgctccaagctgggctgtgtgcacgaaccccccgttcagcccga
ccgctgcgccttatccggtaactatcgtcttgagtccaacccggtaagacacgacttatcgccactggcagcagccactggtaacaggattagcagagcgaggtatgtaggcggtg
ctacagagttcttgaagtggtggcctaactacggctacactagaagaacagtatttggtatctgcgctctgctgaagccagttaccttcggaaaaagagttggtagctcttgatccggca
aacaaaccaccgctggtagcggtggtttttttgtttgcaagcagcagattacgcgcagaaaaaaaggatctcaagaagatcctttgatcttttctacggggtctgacgctcagtggaac
gaaaactcacgttaagggattttggtcatgagattatcaaaaaggatcttcacctagatccttttaaattaaaaatgaagttttaaatcaatctaaagtatatatgagtaaacttggtctgac
agttaccaatgcttaatcagtgaggcacctatctcagcgatctgtctatttcgttcatccatagttgcctgactccccgtcgtgtagataactacgatacgggagggcttaccatctggccc
cagtgctgcaatgataccgcgagacccacgctcaccggctccagatttatcagcaataaaccagccagccggaagggccgagcgcagaagtggtcctgcaactttatccgcctc
catccagtctattaattgttgccgggaagctagagtaagtagttcgccagttaatagtttgcgcaacgttgttgccattgctacaggcatcgtggtgtcacgctcgtcgtttggtatggcttca
ttcagctccggttcccaacgatcaaggcgagttacatgatcccccatgttgtgcaaaaaagcggttagctccttcggtcctccgatcgttgtcagaagtaagttggccgcagtgttatca
ctcatggttatggcagcactgcataattctcttactgtcatgccatccgtaagatgcttttctgtgactggtgagtactcaaccaagtcattctgagaatagtgtatgcggcgaccgagttgc
tcttgcccggcgtcaatacgggataataccgcgccacatagcagaactttaaaagtgctcatcattggaaaacgttcttcggggcgaaaactctcaaggatcttaccgctgttgagat
ccagttcgatgtaacccactcgtgcacccaactgatcttcagcatcttttactttcaccagcgtttctgggtgagcaaaaacaggaaggcaaaatgccgcaaaaaagggaataagg
gcgacacggaaatgttgaatactcatactcttcctttttcaatattattgaagcatttatcagggttattgtctcatgagcggatacatatttgaatgtatttagaaaaataaacaaatagggg
ttccgcgcacatttccccgaaaagtgccacctgacgtcgacggatcgggagatctcccgatcccctatggtgcactctcagtacaatctgctctgatgccgcatagttaagccagtatc
tgctccctgcttgtgtgttggaggtcgctgagtagtgcgcgagcaaaatttaagctacaacaaggcaaggcttgaccgacaattgcatgaagaatctgcttagggttaggcgttttgcg
ctgcttcgcgatgtacgggccagatatacgcgttgacattgattattgactagttattaatagtaatcaattacggggtcattagttcatagcccatatatggagttccgcgttacataactta
cggtaaatggcccgcctggctgaccgcccaacgacccccgcccattgacgtcaataatgacgtatgttcccatagtaacgccaatagggactttccattgacgtcaatgggtggagt
atttacggtaaactgcccacttggcagtacatcaagtgtatcatatgccaagtacgccccctattgacgtcaatgacggtaaatggcccgcctggcattatgcccagtacatgaccttat
gggactttcctacttggcagtacatctacgtattagtcatcgctattaccatggtgatgcggttttggcagtacatcaatgggcgtggatagcggtttgactcacggggatttccaagtctcc
accccattgacgtcaatgggagtttgttttggcaccaaaatcaacgggactttccaaaatgtcgtaacaactccgccccattgacgcaaatgggcggtaggcgtgtacggtgggagg
tctatataagcagagctctctggctaactagagaacccactgcttactggcttatcgaaattaatacgactcactatagggagacccaagctggctagcGccacc

#### Empty vector

An empty pcDNA3.1(+) vector was purchased from Invitrogen (V79020).

#### GLP-1R

Human GLP-1R cDNA was obtained from Addgene (plasmid #213253; GLP-1R-DuET).

#### FLAG-GLP-1R (F61TAG)-1D4

ttgcgcaacgttgttgccattgctacaggcatcgtggtgtcacgctcgtcgtttggtatggcttcattcagctccggttcccaacgatcaaggcgagttacatgatcccccatgttgtgcaa
aaaagcggttagctccttcggtcctccgatcgttgtcagaagtaagttggccgcagtgttatcactcatggttatggcagcactgcataattctcttactgtcatgccatccgtaagatgctt
ttctgtgactggtgagtactcaaccaagtcattctgagaatagtgtatgcggcgaccgagttgctcttgcccggcgtcaatacgggataataccgcgccacatagcagaactttaaaa
gtgctcatcattggaaaacgttcttcggggcgaaaactctcaaggatcttaccgctgttgagatccagttcgatgtaacccactcgtgcacccaactgatcttcagcatcttttactttcac
cagcgtttctgggtgagcaaaaacaggaaggcaaaatgccgcaaaaaagggaataagggcgacacggaaatgttgaatactcatactcttcctttttcaatattattgaagcatttat
cagggttattgtctcatgagcggatacatatttgaatgtatttagaaaaataaacaaataggggttccgcgcacatttccccgaaaagtgccacctgacgtcgacggatcgggagatc
tcccgatcccctatggtgcactctcagtacaatctgctctgatgccgcatagttaagccagtatctgctccctgcttgtgtgttggaggtcgctgagtagtgcgcgagcaaaatttaagcta
caacaaggcaaggcttgaccgacattgcatgaagaatctgcttagggttaggcgttttgcgctgcttcgcgatgtacgggccagatatacgcgttgacattgattattgactagttatta
atagtaatcaattacggggtcattagttcatagcccatatatggagttccgcgttacataacttacggtaaatggcccgcctggctgaccgcccaacgacccccgcccattgacgtca
ataatgacgtatgttcccatagtaacgccaatagggactttccattgacgtcaatgggtggagtatttacggtaaactgcccacttggcagtacatcaagtgtatcatatgccaagtacg
ccccctattgacgtcaatgacggtaaatggcccgcctggcattatgcccagtacatgaccttatgggactttcctacttggcagtacatctacgtattagtcatcgctattaccatggtgat
gcggttttggcagtacatcaatgggcgtggatagcggtttgactcacggggatttccaagtctccaccccattgacgtcaatgggagtttgttttggcaccaaaatcaacgggactttcc
aaaatgtcgtaacaactccgccccattgacgcaaatgggcggtaggcgtgtacggtgggaggtctatataagcagagctctctggctaactagagaacccactgcttactggcttat
cgaaattaatacgactcactatagggagacccaagctggctagcgtttaaacttaagcttagcgccaccatgaagacgatcatcgccctgagctacatcttctgcctggtattcgccg
actacaaggacgatgatgacgccagcatcgatagacctcagggcgctaccgtgagtctttgggagaccgtccagaagtggcgggagtatcgccgccagtgtcaacgcagcctga
cagaagatcctcctccagccaccgacttgtagtgcaataggacatttgatgagtatgcctgctggccagacggcgagcccggctcattcgtgaatgtgtcatgtccatggtacctgcc
atgggcatcatctgtgcctcaaggacacgtctacaggttctgcacagctgagggcctctggctgcaaaaagataacagttcccttccatggcgcgatcttagcgagtgcgaggaatct
aagaggggcgagcgcagttcccccgaggaacagctgctgtttctgtacattatctacaccgtcggctacgcgctgtcctttccgctctggttatcgctagtgccatccttttgggtttcgc
caccttcactgcacaagaaattacatccacttgaatctctttgcaagttttattctgcgggccttgagtgtgttcattaaggacgctgccctgaagtggatgtactctaccgcggcccaaca
gcaccaatggatggactcctcagctatcaggattctctttcctgtcgactcgtgtttctgcttatgcagtactgtgtggcagccaattattattggctgctggtcgaaggagtatatctttaca
cactgctcgcattctccgtgctgagtgaacagtggattttcaggctttacgtgtccattggatggggtgtgcctctgctgttcgtggtgccttggggcatcgtcaagtacctgtatgaagatg
aaggctgttggaccagaaactctaacatgaactactggctcatcatacgcctgcccatactcttcgcgattggagtcaactttcttatatttgtgagagtgatctgtatcgtcgtgtcaaag
ctgaaagctaaccttatgtgcaaaaccgacattaagtgcagactggctaagtccactctcaccctcatcccgctgctcggaacacatgaagtgatatttgccttcgtgatggacgagc
atgcgagaggcacactgcggtttatcaaactgttcacagagctgtctttcactagtttccaaggattgatggtggctattctctactgcttcgtgaacaatgaggtccaacttgaatttcgga
agtcatgggaacggtggcggcttgaacacctccatattcagcgcgatagttctatgaagccacttaaatgtcccacttccagcctgagttccggagccaccgccggctcttcaatgtat
acagccacttgtcaggcgtcctgctctatcgataccggtggacgcaccggtgaattcgatgaggccagcaccaccgtgtctaagaccgagacttctcaggtggcccctgcctaactc
gagtctagagggcccgtttaaacccgctgatcagcctcgactgtgccttctagttgccagccatctgttgtttgcccctcccccgtgccttccttgaccctggaaggtgccactcccactgt
cctttcctaataaaatgaggaaattgcatcgcattgtctgagtaggtgtcattctattctggggggtggggtggggcaggacagcaagggggaggattgggaagacaatagcaggc
atgctggggatgcggtgggctctatggcttctgaggcggaaagaaccagctggggctctagggggtatccccacgcgccctgtagcggcgcattaagcgcggcgggtgtggtggtt
acgcgcagcgtgaccgctacacttgccagcgccctagcgcccgctcctttcgctttcttcccttcctttctcgccacgttcgccggctttccccgtcaagctctaaatcgggggctcccttta
gggttccgatttagtgctttacggcacctcgaccccaaaaaacttgattagggtgatggttcacgtagtgggccatcgccctgatagacggtttttcgccctttgacgttggagtccacgtt
ctttaatagtggactcttgttccaaactggaacaacactcaaccctatctcggtctattcttttgatttataagggattttgccgatttcggcctattggttaaaaaatgagctgatttaacaaa
aatttaacgcgaattaattctgtggaatgtgtgtcagttagggtgtggaaagtccccaggctccccagcaggcagaagtatgcaaagcatgcatctcaattagtcagcaaccaggtgt
ggaaagtccccaggctccccagcaggcagaagtatgcaaagcatgcatctcaattagtcagcaaccatagtcccgcccctaactccgcccatcccgcccctaactccgcccagtt
ccgcccattctccgccccatggctgactaattttttttatttatgcagaggccgaggccgcctctgcctctgagctattccagaagtagtgaggaggcttttttggaggcctaggcttttgca
aaaagctcccgggagcttgtatatccattttcggatctgatcaagagacaggatgaggatcgtttcgcatgattgaacaagatggattgcacgcaggttctccggccgcttgggtgga
gaggctattcggctatgactgggcacaacagacaatcggctgctctgatgccgccgtgttccggctgtcagcgcaggggcgcccggttctttttgtcaagaccgacctgtccggtgcc
ctgaatgaactgcaggacgaggcagcgcggctatcgtggctggccacgacgggcgttccttgcgcagctgtgctcgacgttgtcactgaagcgggaagggactggctgctattgg
gcgaagtgccggggcaggatctcctgtcatctcaccttgctcctgccgagaaagtatccatcatggctgatgcaatgcggcggctgcatacgcttgatccggctacctgcccattcga
ccaccaagcgaaacatcgcatcgagcgagcacgtactcggatggaagccggtcttgtcgatcaggatgatctggacgaagagcatcaggggctcgcgccagccgaactgttcg
ccaggctcaaggcgcgcatgcccgacggcgaggatctcgtcgtgacccatggcgatgcctgcttgccgaatatcatggtggaaaatggccgcttttctggattcatcgactgtggcc
ggctgggtgtggcggaccgctatcaggacatagcgttggctacccgtgatattgctgaagagcttggcggcgaatgggctgaccgcttcctcgtgctttacggtatcgccgctcccgat
tcgcagcgcatcgccttctatcgccttcttgacgagttcttctgagcgggactctggggttcgaaatgaccgaccaagcgacgcccaacctgccatcacgagatttcgattccaccgcc
gccttctatgaaaggttgggcttcggaatcgttttccgggacgccggctggatgatcctccagcgcggggatctcatgctggagttcttcgcccaccccaacttgtttattgcagcttataa
tggttacaaataaagcaatagcatcacaaatttcacaaataaagcatttttttcactgcattctagttgtggtttgtccaaactcatcaatgtatcttatcatgtctgtataccgtcgacctctag
ctagagcttggcgtaatcatggtcatagctgtttcctgtgtgaaattgttatccgctcacaattccacacaacatacgagccggaagcataaagtgtaaagcctggggtgcctaatgagt
gagctaactcacattaattgcgttgcgctcactgcccgctttccagtcgggaaacctgtcgtgccagctgcattaatgaatcggccaacgcgcggggagaggcggtttgcgtattggg
cgctcttccgcttcctcgctcactgactcgctgcgctcggtcgttcggctgcggcgagcggtatcagctcactcaaaggcggtaatacggttatccacagaatcaggggataacgcag
gaaagaacatgtgagcaaaaggccagcaaaaggccaggaaccgtaaaaaggccgcgttgctggcgtttttccataggctccgcccccctgacgagcatcacaaaaatcgacg
ctcaagtcagaggtggcgaaacccgacaggactataaagataccaggcgtttccccctggaagctccctcgtgcgctctcctgttccgaccctgccgcttaccggatacctgtccgc
ctttctcccttcgggaagcgtggcgctttctcatagctcacgctgtaggtatctcagttcggtgtaggtcgttcgctccaagctgggctgtgtgcacgaaccccccgttcagcccgaccgc
tgcgccttatccggtaactatcgtcttgagtccaacccggtaagacacgacttatcgccactggcagcagccactggtaacaggattagcagagcgaggtatgtaggcggtgctaca
gagttcttgaagtggtggcctaactacggctacactagaagaacggtatttggtatctgcgctctgctgaagccagttaccttcggaaaaagagttggtagctcttgatccggcaaaca
aaccaccgctggtagcggtggtttttttgtttgcaagcagcagattacgcgcagaaaaaaaggatctcaagaagatcctttgatcttttctacggggtctgacgctcagtggaacgaaa
actcacgttaagggattttggtcatgagattatcaaaaaggatcttcacctagatccttttaaattaaaaatgaagttttaaatcaatctaaagtatatatgagtaaacttggtctgacagtta
ccaatgcttaatcagtgaggcacctatctcagcgatctgtctatttcgttcatccatagttgcctgactccccgtcgtgtagataactacgatacgggagggcttaccatctggccccagt
gctgcaatgataccgcgagacccacgctcaccggctccagatttatcagcaataaaccagccagccggaagggccgagcgcagaagtggtcctgcaactttatccgcctccatc
cagtctattaattgttgccgggaagctagagtaagtagttcgccagttaatagt

**Table S6**: Primers used for cloning

| Name | Sequence (5'-3') |
|---|---|
| **ACTB primers** | |
| AgeI restriction site_F | AAATTTACCGGTCGCCACCATGGATGATGATATCGCCGCGCTC |
| BamHI restriction site_R | AAATTTGGATCCCTAGAAGCATTTGCGG |
| ACTB_118K*_F | tcatgtttgagaccttcaacacccc |
| ACTB_118K*_R | tctgggtcatCTActcgcggttgg |
| ACTB_Cterm_HA_F | CCAGATTACGCTtagggatccaccggatctagataa |
| ACTB_Cterm_HA_R | AACATCGTATGGGTAgaagcatttgcggtgga |
| **NONO primers** | |
| NONO_258A*_F | gcagcctggctcctttgagtatg |
| NONO_258A*_R | taaaatctgggtggctgctctc |
| **CCR5 primers** | |
| CCR5_F96*_F | gcccagtgggactagggaaatacaatgtgtcaac |
| CCR5_F96*_R | ggcagcatagtgagcccagaagg |
| CCR5_L352*_F | CTAGTACCCATACGATGTTCCAG |
| CCR5_L352*_R | cccacagatatttcctgc |
| Cterm_HA_F | AGATTACGCTtgataactcgagtctagaggg |
| Cterm_HA_R | GGAACATCGTATGGGTAcaagcccacagatatttcct |
| **GLP-1R primers** | |
| GLP1R_61F*_F | tgcaataggacatttgatgagtatgcctgctg |
| GLP1R_61F*_R | ctacaagtcggtggctggaggaggatc |

#### Chemical structures

Iridium (G2)-PEG16-tetrazine (**1**)

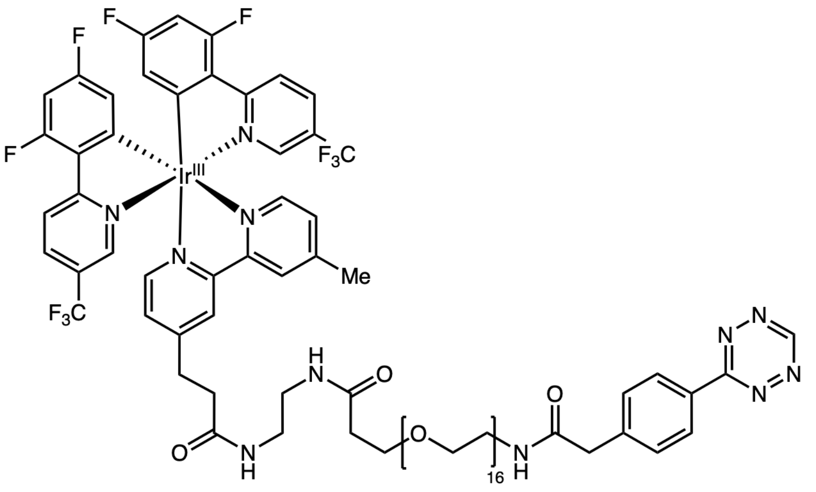

Biotin-PEG3-Diazirine (**S1**)

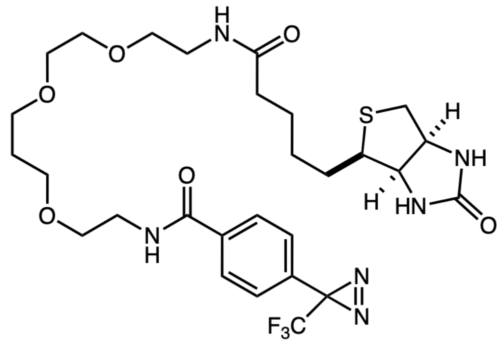

Tetrazine-PEG$_4$-TAMRA (**S2**)

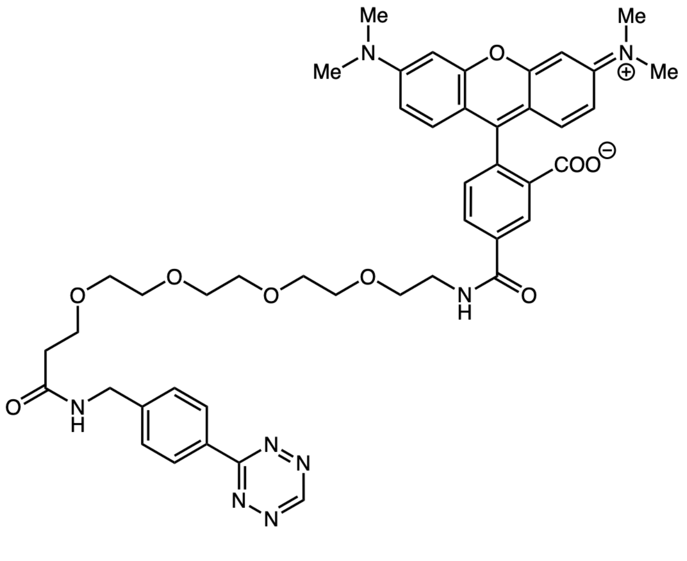

**Syntheses**

Tetrazine-PEG16-COOH (**S3**)

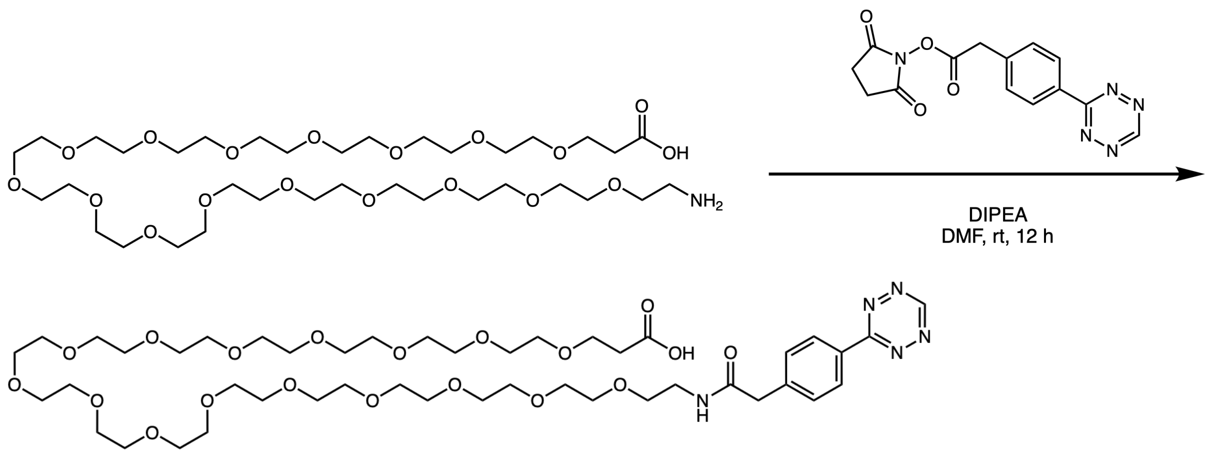

To a stirred solution of the amine-PEG16-COOH (126.7 mg, 0.16 mmol, 1.0 equiv, MedChemExpress, cat. HY-140179) in anhydrous DMF (0.2 M) was added tetrazine-NHS ester (50 mg, 0.16 mmol, 1.0 equiv, Vector Laboratories, cat. CCT-1127-25), followed by DIPEA (83 µL, 0.48 mmol, 3.0 equiv). The reaction mixture was stirred at room temperature for 12 h. The crude mixture was then concentrated under reduced pressure to afford the desired tetrazine-$PEG_{16}$-COOH as a pink oil (95 mg, 60% yield) and was directly used for the next step without further purification.

Tetrazine-$PEG_{16}$-NHS ester (**S4**)

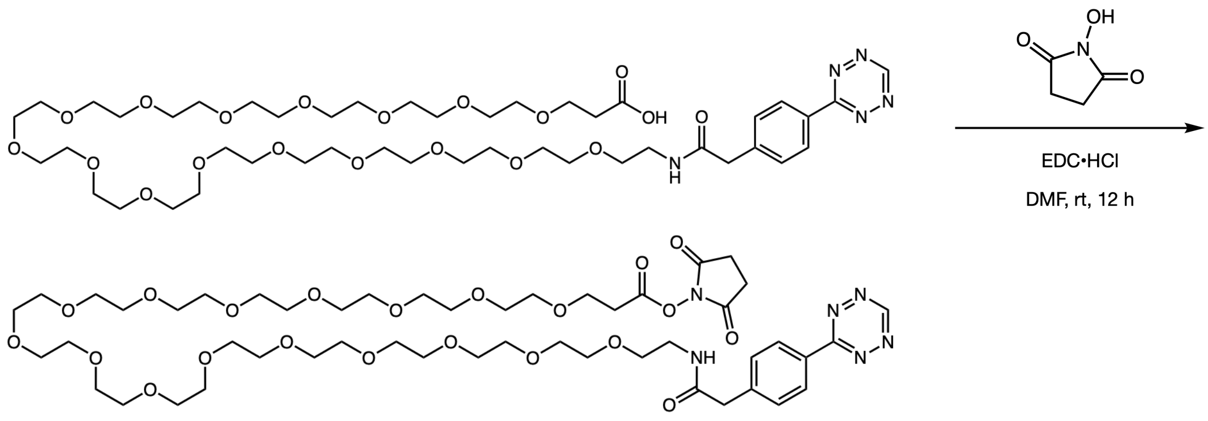

S3 and EDC hydrochloride (1.4 equiv.), and *N*-hydroxysuccinimide (1.2 equiv.) were added to a 40 mL vial equipped with a magnetic stir bar and then dissolved in DMF (0.2 M). The reaction mixture was stirred at room temperature for 12 h. The crude mixture was then concentrated under reduced pressure to afford the desired tetrazine-PEG$_{16}$-COOH as a pink oil (95 mg, 60% yield) and was directly used for the next step without further purification.

Tetrazine-PEG$_{16}$-Ir (**1**)

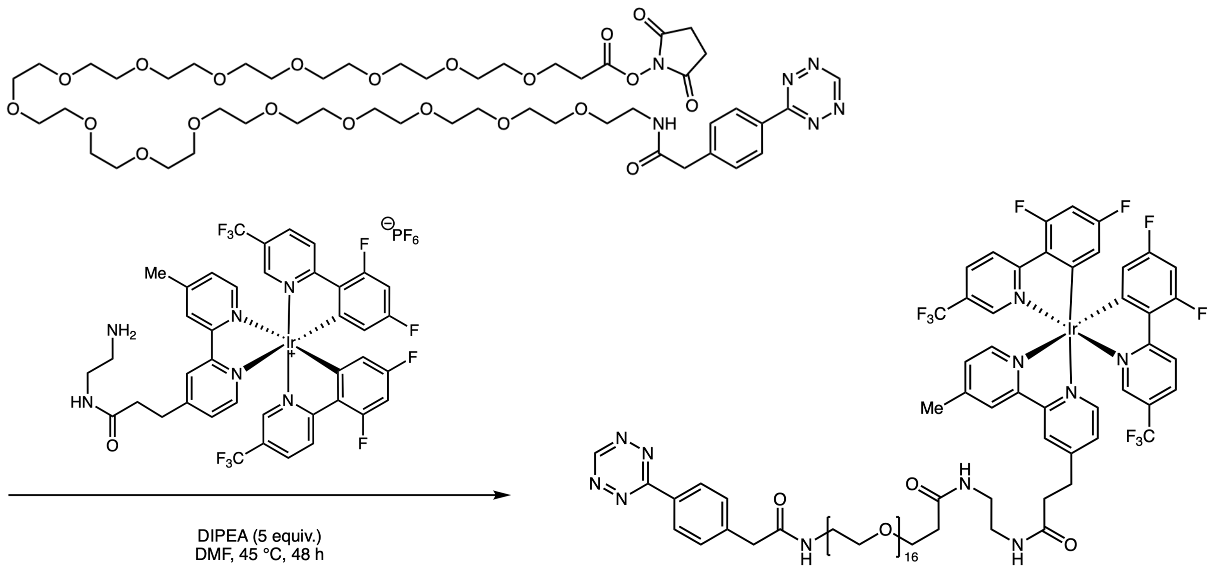

Tetrazine-PEG$_{16}$-NHS ester (63.7 mg, 58.5 µmol, 1.0 equiv) was dissolved in anhydrous DMF (0.2 M). Ir–ENH$_2$ (116.2 mg, 116.9 µmol, 2.0 equiv), which was prepared as previously reported,[2,3] was added, followed by DIPEA (51 µL, 292 µmol, 5.0 equiv). The reaction mixture was stirred at 45 °C for 48 h. The crude reaction mixture was purified via prep-HPLC (10-100% ACN/H$_2$O w/ 0.1% formic acid) to afford Ir–tetrazine (**1**) (25.3 mg, 12.9 µmol, 22%) as a bright orange-red oil.

**[1]H-NMR** (500 MHz, MeOD) δ 10.32 (s, 1H), 8.71 (s, 2H), 8.60 – 8.49 (m, 4H), 8.36 – 8.27 (m, 2H), 7.93 (dd, *J* = 22.8, 5.8 Hz, 2H), 7.72 (d, *J* = 19.4 Hz, 2H), 7.64 – 7.50 (m, 4H), 6.87 – 6.75 (m, 2H), 5.82 – 5.73 (m, 2H), 3.72 – 3.54 (m, 69H), 3.40 (t, *J* = 5.4 Hz, 2H), 3.27 – 3.21 (m, 4H), 3.18 (t, *J* = 7.4 Hz, 2H), 2.69 – 2.62 (m, 5H), 2.41 (t, *J* = 6.0 Hz, 2H).

**[13]C-NMR** (126 MHz, MeOD) δ 174.3, 173.9, 173.2, 169.19, 169.18, 167.6, 167.4, 167.3, 165.3, 165.2, 165.1, 163.1, 163.0, 159.3, 157.8, 156.8, 156.7, 156.4, 155.1, 151.7, 151.3, 146.5, 142.6, 138.4, 133.4, 132.0, 131.30, 131.25, 131.0, 130.2, 129.3, 127.8, 127.4, 127.1, 126.8, 125.22, 125.18, 125.1, 125.0, 124.5, 122.3, 115.3, 115.2, 115.1, 100.8, 100.6, 100.4, 71.6, 71.5, 71.4, 71.31, 71.26, 70.5, 68.2, 43.7, 40.7, 40.3, 39.7, 37.7, 36.2, 31.8, 21.5.

**[19]F NMR** (282 MHz, MeOD) δ -64.32 – -64.42 (m), -74.66 (d, *J* = 707.5 Hz), -104.40 – -104.60 (m), -107.95 – -108.13 (m).

**HRMS (ESI-TOF)**: m/z calculated for $C_{85}H_{105}F_{10}IrN_{11}O_{19}$ ($[M]^+$) 1966.7058, found 1966.7351

#### NMR Spectroscopy Data

Iridium-tetrazine (**1**) $^1$H NMR in MeOD

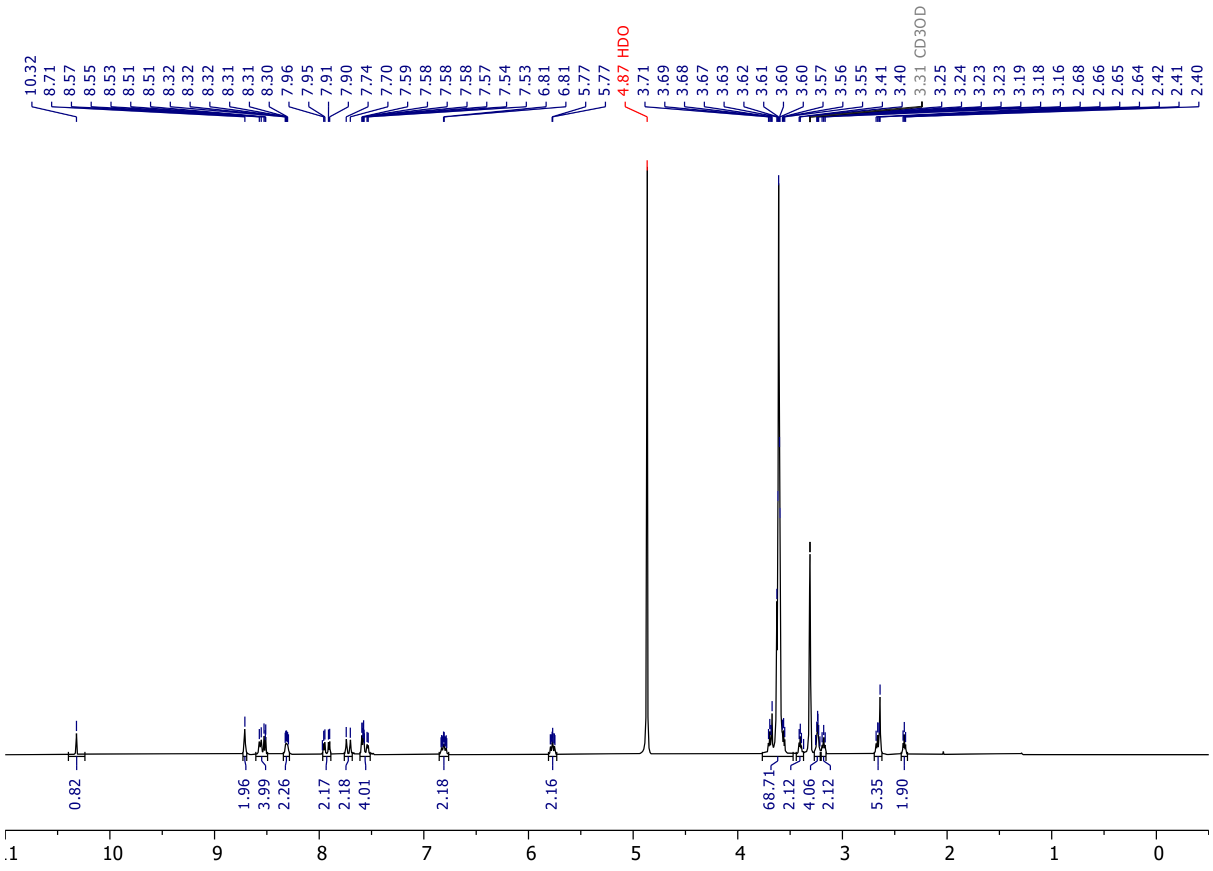

Iridium-tetrazine (**1**) $^{13}$C NMR in MeOD

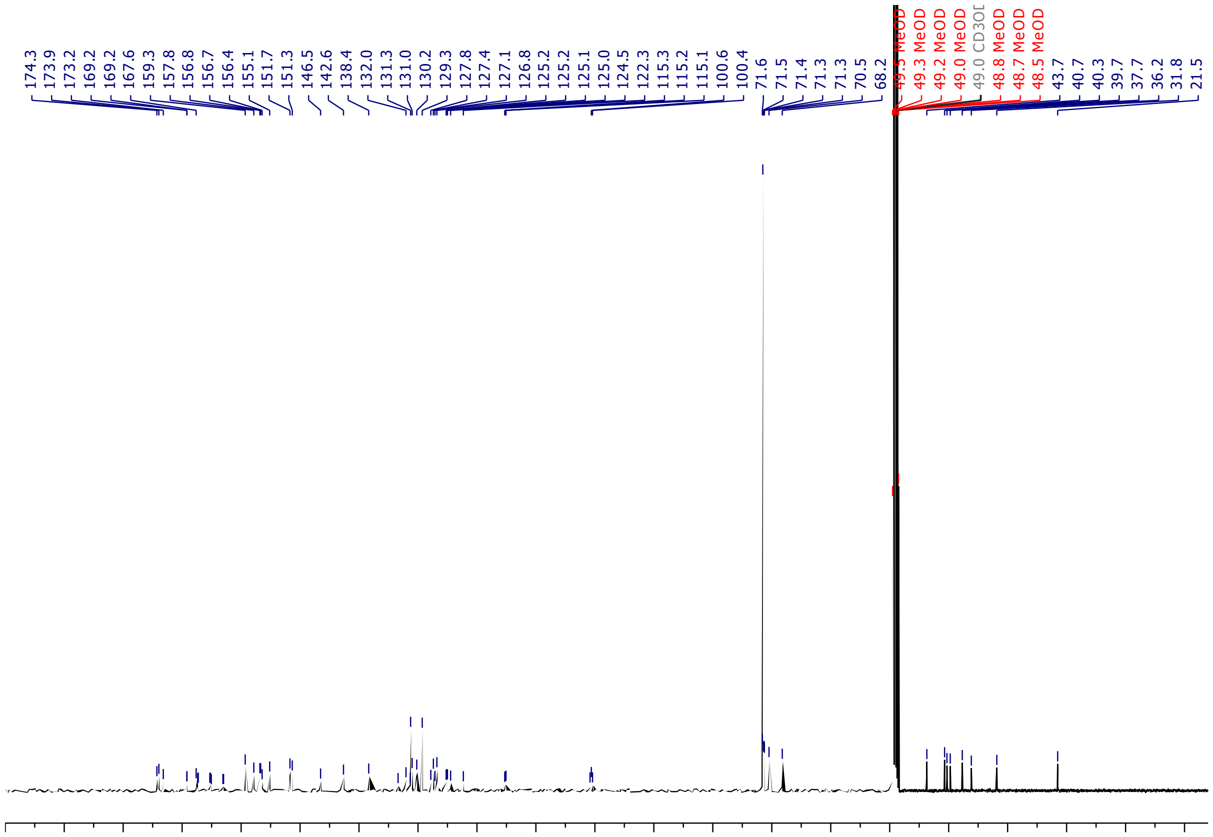

Iridium-tetrazine (**1**) $^{19}F$ NMR in MeOD

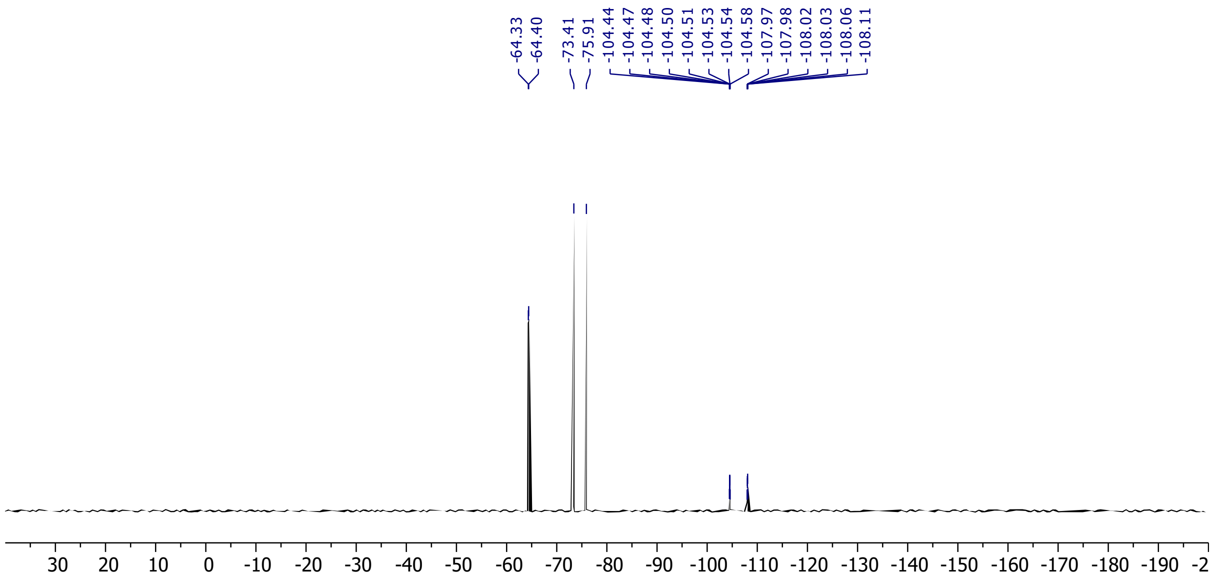

**References**

(1) Mihaila, T. S.; Bäte, C.; Ostersehlt, L. M.; Pape, J. K.; Keller-Findeisen, J.; Sahl, S. J.; Hell, S. W. Enhanced Incorporation of Subnanometer Tags into Cellular Proteins for Fluorescence Nanoscopy via Optimized Genetic Code Expansion. *Proc. Natl. Acad. Sci. U.S.A.* **2022**, *119* (29), e2201861119. https://doi.org/10.1073/pnas.2201861119.

(2) Huth, S. W.; Oakley, J. V.; Seath, C. P.; Geri, J. B.; Trowbridge, A. D.; Parker, D. L.; Rodriguez-Rivera, F. P.; Schwaid, A. G.; Ramil, C.; Ryu, K. A.; White, C. H.; Fadeyi, O. O.; Oslund, R. C.; MacMillan, D. W. C. μMap Photoproximity Labeling Enables Small Molecule Binding Site Mapping. *J. Am. Chem. Soc.* **2023**, *145* (30), 16289–16296. https://doi.org/10.1021/jacs.3c03325.

(3) Trowbridge, A. D.; Seath, C. P.; Rodriguez-Rivera, F. P.; Li, B. X.; Dul, B. E.; Schwaid, A. G.; Buksh, B. F.; Geri, J. B.; Oakley, J. V.; Fadeyi, O. O.; Oslund, R. C.; Ryu, K. A.; White, C.; Reyes-Robles, T.; Tawa, P.; Parker, D. L.; MacMillan, D. W. C. Small Molecule Photocatalysis Enables Drug Target Identification via Energy Transfer. *Proc. Natl. Acad. Sci. U.S.A.* **2022**, *119* (34), e2208077119. https://doi.org/10.1073/pnas.2208077119.

(4) Geri, J. B.; Oakley, J. V.; Reyes-Robles, T.; Wang, T.; McCarver, S. J.; White, C. H.; Rodriguez-Rivera, F. P.; Parker, D. L.; Hett, E. C.; Fadeyi, O. O.; Oslund, R. C.; MacMillan, D. W. C. Microenvironment Mapping via Dexter Energy Transfer on Immune Cells. *Science* **2020**, *367* (6482), 1091–1097. https://doi.org/10.1126/science.aay4106.

(5) Zhou, Y.; Zhou, B.; Pache, L.; Chang, M.; Khodabakhshi, A. H.; Tanaseichuk, O.; Benner, C.; Chanda, S. K. Metascape Provides a Biologist-Oriented Resource for the Analysis of Systems-Level Datasets. *Nature Communications* **2019**, *10* (1), 1523. https://doi.org/10.1038/s41467-019-09234-6.
